# Decoding the chemical language of RiPPs from the untapped Archaea domain

**DOI:** 10.1101/2024.10.07.616454

**Authors:** Zhi-Man Song, Cunlei Cai, Ying Gao, Xiaoqian Lin, Qian Yang, Dengwei Zhang, Gengfan Wu, Haoyu Liang, Qianlin Zhuo, Junliang Zhang, Peiyan Cai, Haibo Jiang, Wenhua Liu, Yong-Xin Li

## Abstract

Chemical communication is crucial in ecosystems with complex microbial communities. However, the difficulties inherent to the cultivation of archaea have led to a limited understanding of their chemical language, especially regarding the structure diversity and function of secondary or specialized metabolites (SMs). Our comprehensive investigation into the biosynthetic potential of archaea, combined with metabolic analyses and the first report of heterologous expression in archaea, has unveiled the previously unexplored biosynthetic capabilities and chemical diversity of archaeal ribosomally synthesized and post-translationally modified peptides (RiPPs). We have identified twenty-four new lanthipeptides of RiPPs exhibiting unique chemical characteristics, including a novel subfamily featuring an unexplored type with diamino-dicarboxylic (DADC) termini, largely expanding the chemical landscape of archaeal SMs. This sheds light on the chemical novelty of archaeal metabolites and emphasizes their potential as an untapped resource for natural product discovery. Additionally, archaeal lanthipeptides demonstrate specific antagonistic activity against haloarchaea, mediating the unique biotic interaction in the halophilic niche. Furthermore, they showcase a new ecological role of RiPPs in enhancing the host’s motility by inducing the rod-shaped cell morphology and upregulating the archaellin gene expression, facilitating the archaeal interaction with abiotic environments. These discoveries broaden our understanding of archaeal chemical language and provide promising prospects for future exploration of SM-mediated interaction.

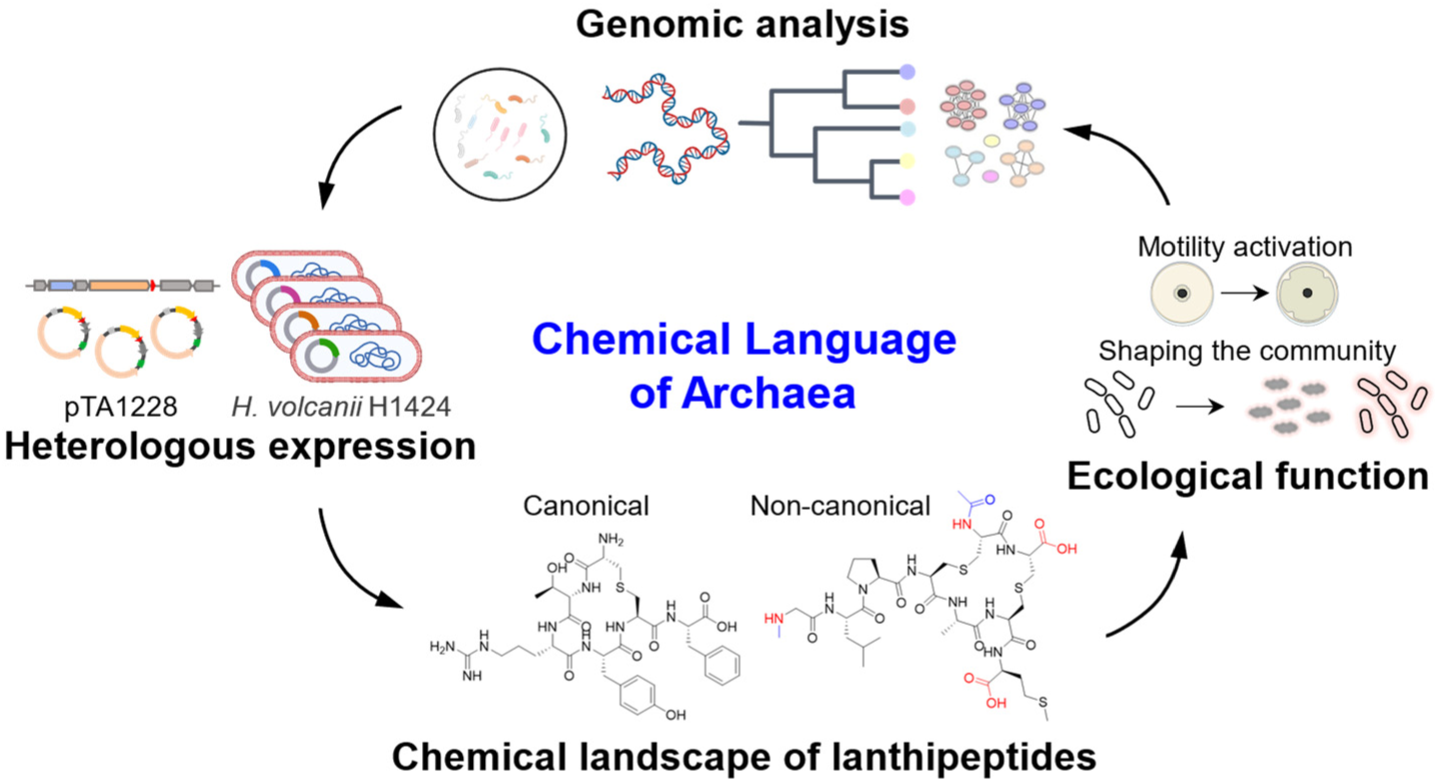

## Main

Archaea, the third domain of life, have evolved distinctly from bacteria and eukaryotes and are universally present in varied ecological niches^1^. They are particularly abundant in extreme environments, where their cultivation often poses challenges despite their significant roles in ecosystems^2^. Archaea have developed various mechanisms to thrive in different environments, involving direct and indirect interactions with biotic and abiotic factors^3–9^. These interactions extend to symbiosis, predation, antagonism, mutualism, and participation in essential processes like carbon, nitrogen, and sulfur cyclings. Bacteria and fungi are renowned for their production of a wide range of secondary or specialized metabolites (SMs), which include signaling molecules, antibiotics, and siderophores that aid in their interaction with their biotic and abiotic environments. However, our understanding of chemical-mediated interactions in archaea lags far behind that of bacteria, except for certain UV-protective carotenoids^10^, putative siderophores^11^ and quorum-sensing molecules^12,13^.

Archaea possess a variety of enzymes and metabolic pathways distinct from those in bacteria^14–18^, marking them as a promising and largely untapped reservoir of novel enzymes and SMs. However, due to the challenges in archaeal cultivation and genetic manipulation, the chemical landscape, biosynthesis, and ecological function of archaeal metabolites remain largely unexplored^2,19,20^. Haloarchaea, easily lab-cultured and known for producing antagonistic proteins like halocins^21,22^, are ideal models for studying archaeal interactions, cell biology, and metabolic pathways^6,23–25^. Despite the presence of hundreds of biosynthetic gene clusters (BGCs)^26–29^ and numerous attempts of wild-type strain fermentation, previous research identified only two ribosomally synthesized and post-translationally modified peptides (RiPPs) from Haloarchaea^26^. Expressing archaeal biosynthetic pathways heterologously in bacterial hosts can be problematic due to distinct metabolic features, potentially leading to improper folding, low expression levels, or inactive archaeal enzymes^30^. The scarcity of SMs from even well-defined archaeal BGCs predicted bioinformatically *in silico*^26–29,31^, alongside the challenges involved in their heterologous expression in model bacterial hosts, underscores the necessity for new strategies in discovering archaeal metabolites.

Uncovering the yet unexplored chemistry of archaeal SMs will provide crucial insights into their chemically mediated interactions, invigorating the relatively underexplored field of archaeal chemical biology. Here, our study reports the systematic investigation and the first heterologous expression-based discovery of SMs in archaea, aiming to uncover their biosynthetic potential, chemical landscape, and ecological functions. We successfully validated twenty-four classic and noncanonical lanthipeptides of RiPPs originating from eleven BGCs by combining the heterologous expression with biosynthetic rule-based metabolomic analysis^32^. These archaeal lanthipeptides exhibited a specific antagonistic activity against Haloarchaea. Intriguingly, our investigation unveiled a new ecological function of lanthipeptides in stimulating the heterologous host’s motility, by promoting a rod-like morphology and upregulating archaellin gene expression. These observations indicate that lanthipeptides potentially enhance the motility of archaea, thereby improving nutrient uptake or space occupation for environmental adaptation. Our study represents a significant advancement in understanding the archaeal chemical landscape and paves the way for archaeal natural product discovery and chemical ecology study.

## Results

### Genome mining reveals the largely untapped biosynthetic potential of archaeal RiPPs

Our previous genomic analysis indicated that archaea contain a relatively small number of BGCs per genome compared to the extensively studied and well-documented bacterial counterparts^26^. They encode a broad range of known SM classes, such as RiPPs, terpenes, nonribosomal peptides (NRPs), and polyketides (PKs). Our initial fermentation-based discovery identified a new lanthipeptide from Haloarchaea, marking the first discovery of lantibiotic and anti-archaeal SM in the archaeal domain^26^. In the current study, we further applied the antiSMASH 7.0 tool^33^, which is capable of analyzing both bacterial and archaeal BGCs, to examine 11,644 archaeal genomes from the NCBI database to delve deeper into exploring the biosynthetic landscape, diversity, and novelty of archaeal RiPPs. From 3,451 representative archaeal genomes, 7,731 BGCs were identified, mainly from phyla Halobacteriota (3,379, 43.7%), Thermoplasmatota (1,359, 17.6%), Methanobacteriota (1,271, 16.4%), Thermoproteota (962, 12.4%) and Nanoarchaeota (221, 2.9%) (Fig. 1a and Supplementary Data 1). Among them, RiPPs (3,599, 46.6%), Terpenes (1,990, 25.7 %), and NRPs (581, 7.5 %) emerged as dominant BGCs in the domain of Archaea (Fig.1a). These BGCs exhibited significant novelty compared to known BGCs from MIBiG^34^, indicating a largely unexplored source for natural product discovery.

**Figure 1.**
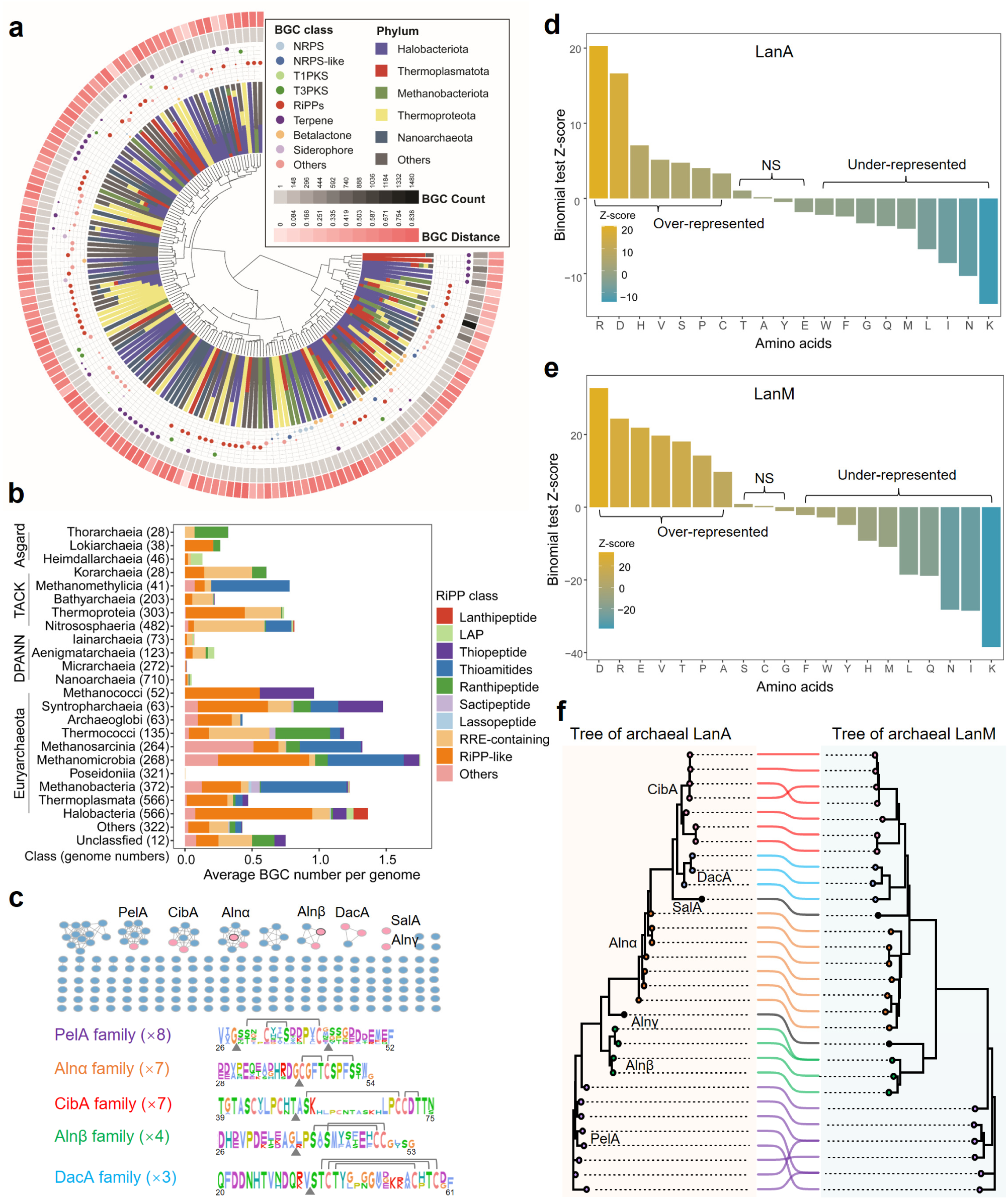
Uncovering the biosynthetic potential of archaeal RiPPs. **a**, The novelty, diversity, and distribution of archaeal BGCs. Layers from innermost to outermost show hierarchical clustering of the 148 gene cluster clans (GCCs) based on the BGC distances, phylum distribution for each clan, the proportion of different BGC classes for each clan, BGC count, and average cosine distance to known BGCs. **b**, The distribution of RiPP BGCs in the domain of Archaea at the class level. **c**, The SSN analysis of putative archaeal LanAs and sequence logos of experimentally verified LanA families. Sequences with 100% identity are grouped into a single node. Pink nodes represent the characterization of heterogeneous expression products or wild-type natural products in this study. Lines show the validated crosslinks between C and S/T residues. The main cleavage sites observed between the leader and core peptides are displayed as triangles below the LanAs. **d-e**, Binomial tests were conducted to compare the proportions of all 20 amino acids in LanAs (**d**) and LanMs (**e**) between archaeal and bacterial counterparts. *Z*-scores were calculated relative to archaeal LanAs and LanMs, where *Z* > 2.0 indicated significant enrichment of a given amino acid in Archaea (’Over-represented’), *Z* < −2.0 indicated significant depletion (’Under-represented’), and some amino acids showed no significant biases (’NS’). **f**, Co-phylogenomic analysis of archaeal LanAs (left) and LanMs (right) from verified SSN families. The nodes and lines are colored according to their LanA families in **c**. Two black notes represent verified orphan notes in the SSN analysis, SalA and Alnβ.

Phylogenetic distribution analysis unveiled a remarkable diversity within RiPP BGCs, widely distributed across the Asgard, TACK, DPANN superphyla, and predominantly within Euryarchaeota (Fig. 1a). Among these abundant RiPP classes are YcaO-related RiPPs (thioamitides and thiopeptides), radical S-adenosylmethionine (rSAM)-dependent enzyme-modified RiPPs (ranthipeptides and sactipeptides), lanthipeptides, and lassopeptides (Fig. 1b). While RiPPs are widely distributed throughout different phyla/classes, the scarcity of identified archaeal RiPPs hinders us from elucidating their unique chemical structure and putative ecological functions. Our attempts to activate YcaO- and rSAM-related BGCs in the wild-type strain or through heterologous expression have been unsuccessful, prompting a shift in focus towards lanthipeptides. Class II lanthipeptides, unique among other RiPPs, are solely present in Haloarchaea (refer to the class Halobacteria), implying halophilic niche-specific metabolites (Fig. 1b, Supplementary Data 1 and 2). Consequently, an updated library was constructed *in silico* to decipher the chemical landscape of archaeal lanthipeptides. This library contains 353 putative precursors (LanAs) and 110 lanthipeptide synthase LanMs from 102 archaeal lanthipeptide BGCs, through antiSMASH-based genome mining method^26,33^ (Supplementary Data 2). Among these, 67 deduplicated archaeal LanMs, exclusively present in Haloarchaea, are mainly from the genera *Halorussus* (26, 38.8%) and *Haloferax* (14, 20.9%), followed by *Haladaptatus* (5, 7.5%), *Halobacteriales* (5, 7.5%), *Natrinema* (5, 7.5%) (Supplementary Fig. 1a and Supplementary Data 2). Moreover, 196 unique LanA sequences distinctive from bacterial LanA have been categorized into 157 families through sequence similarity network (SSN) analysis (Fig. 1c), highlighting untapped chemical space for lanthipeptide discovery.

### Co-evolved archaeal LanMs and LanAs exhibit environmental adaptation

Our collective analysis of the phylogenetic tree, amino acid composition bias, SSN, and sequence logo indicates a unique chemical distinctiveness of archaeal lanthipeptides compared to their bacterial counterparts (Fig. 1 and Supplementary Figs. 1-2). The phylogenetic tree of archaeal LanMs highlighted their evolutionary divergence from bacterial LanMs (Supplementary Fig. 1b and Supplementary Data 2). The tree of archaeal LanAs exhibited two primary clades: one encompassing our previously characterized LanAs, Alnα and Alnβ, and another unexplored clade that offers prospects for further investigation (Supplementary Fig. 2). Additionally, archaeal LanAs are predicted to be shorter and comprise a higher portion of C and S instead of T residues than bacterial LanAs (Fig. 1c, d), which are crucial conserved residues for thioether ring formation in lanthipeptide biosynthesis^35^. More importantly, the distinctive amino acid composition bias of archaeal LanAs and LanMs supports the hypothesis that they underwent eco-evolution in response to hypersaline environments (Fig. 1d, e). Specifically, archaeal LanMs and LanAs evolved to over-represent D, V, R, and P residues while under-representing K, I, N, Q, M, L, W, and F residues (Fig. 1d, e). The increased abundance of D/E residues and the reduced portion of K residue is linked to the "salt-in" strategy of Haloarchaea, enabling interactions with cations in their high-salinity environment^6,36^. Furthermore, the reduced occurrence of hydrophobic amino acids (I, L, M, W) may increase random-coil structure formation, thereby preventing protein aggregation or inactivation in low-water environments. The prevalence of basic R residue in archaeal LanMs and LanAs, as pronounced as acidic D residue, may be a unique evolutionary trait of archaeal lanthipeptide biosynthetic pathways that have not been observed in other protein analyses to the best of our knowledge^36^. Considering the evolutionary divergence of LanMs and their amino acid composition biases, the co-evolutionary relationship between archaeal LanMs and LanAs hints at a unique evolutionary path for archaeal lanthipeptides, which occupy distinct chemical realms (Fig. 1f). These results imply that archaeal LanAs, having co-evolved with LanMs as a part of environmental adaptation, could potentially enhance the chemical diversity of lanthipeptides exclusive to Haloarchaea, thereby playing significant ecological roles in halophilic environments.

### Heterologous expression-based discovery of archaeal lanthipeptides

Moving beyond the traditional method of natural product discovery through wild-type cultivation, we successfully heterologously expressed eleven BGCs from various SSN families in *Haloferax volcanii* H1424 using the pTA1228 vector^37,38^ (Fig. 2a, Supplementary Table 2 and Supplementary Data 3). The procedure was initiated by cloning our previously verified lanthipeptide BGCs^26^, *alnα* and *alnβ*, which resulted in the production of corresponding analogs of the natural lanthipeptides, archalans α2 (**7**), β1 (**9**), and β2 (**10)** (Fig. 2b, c and Supplementary Figs. 3-4). It marks the first endeavor to authenticate archaeal RiPP BGCs through heterologous expression (Fig. 2), laying the groundwork for the subsequent expression of other archaeal lanthipeptide BGCs. Based on this approach, we heterologously expressed fifteen lanthipeptide BGCs, covering all seven major lanthipeptide families in SSN (Fig.1). This endeavor linked nine BGCs from five SSN families and two orphan BGCs with their lanthipeptide products, representing the first comprehensive exploration of archaeal SMs so far (Figs.1, 2 and Supplementary Table 1).

**Figure 2.**
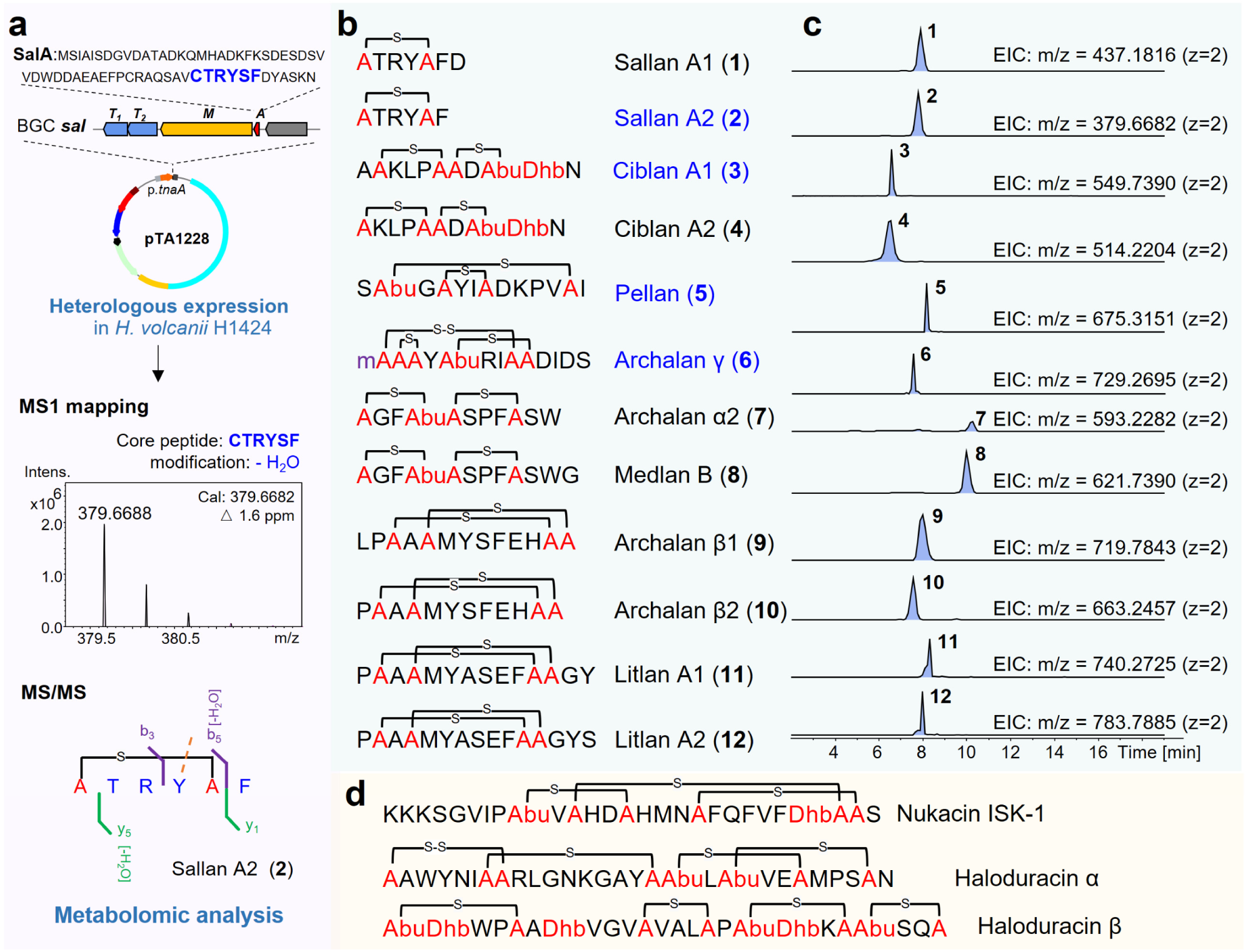
Deciphering novel archaeal class II lanthipeptides through heterologous expression and biosynthetic rule-based metabolic analysis. **a**, Workflow for the heterologous expression-based discovery of eleven lanthipeptide BGCs in *H. volcanii* H1424, in conjunction with metabolic analysis, is illustrated through the example of sallan A2 (**2**). Classical lanthipeptides (**b-c**) with different modifications (’m’ in purple represents methylation) were identified via HR LC-MS. Structures confirmed through NMR analysis or chemical derivatization are marked in blue. **d**, Representative bacterial class II lanthipeptides.

Except for the unusual peptides from DcaA family, nine other classical lanthipeptides were pinpointed by biosynthetic rule-based metabolic analysis^32^(Fig. 2 and Supplementary Figs. 5-18), including sallans (**1** and **2**, from BGC *sal*), ciblans (**3** and **4**, from BGCs *cib1* and *cib2*), pellan (**5**, from BGC *pel*), archalan γ (**6**, from BGC *alnγ*), medlan B (**8**, from BGC *medb*) and litlans (**11** and **12**, from BGC *lit*). It’s important to note that sallans (**1** and **2**), pellan (**5**), and litlans (**11** and **12**) were not detected in their corresponding wild-type strains. Others were detected in relatively low titers compared to the results from heterologous expression. Most of these lanthipeptides are detected in low quantities, either in the wild type or the heterologous host. Over the past decades, this has been a major obstacle in discovering natural products from Archaea.

To pinpoint the low-yield BGC-encoded peptides within the metabolic profile, we mapped the calculated mass data, which was predicted based on lanthipeptide post-translational modification rules^32^, with the HRMS data to detect corresponding hits. For instance, a UPLC-HRMS analysis of the culture extract from the heterologous expression strain of the BGC *sal* (*Halorussus salinus* YJ-37-H) exposed numerous low peptidic signals (Supplementary Fig. 5). The mass matching process revealed a hit of [M + 2H]^2+^ = 379.6688 (**2**) that matched the calculated mass data well of "CTRYSF" with one dehydration (calculated [M + 2H]^2+^ = 379.6682), linking this low-yield peptide to BGC *sal* (Fig. 2a). This underscores the benefit of using heterologous expression coupled with rule-based metabolic analysis to uncover the hidden chemistry in Archaea, especially those with low yields.

### Identification of new classical lanthipeptides from Archaea

The structures of classical lanthipeptides **1**-**4** were elucidated through a comprehensive approach that integrated High-Resolution Mass Spectrometry (HRMS), nuclear magnetic resonance (NMR) analysis, and advanced Marfey’s analysis. Following biosynthetic rule-based metabolic analysis, sallans A1 (**1**, observed [M+2H]^2+^ = 437.1816, calculated [M+2H]^2+^ = 437.1816, Δ = 0.0 ppm) and A2 (**2**, observed [M+2H]^2+^ = 379.6688, calculated [M+2H]^2+^ = 379.6682, Δ = 1.6 ppm) were found to be analogs encoded by BGC *sal*, which were predicted with one dehydration (-H_2_O, −18 Da) on their corresponding core peptides (CTRYSFD for **1** and CTRYSF for **2**, Fig. 2a, Supplementary Fig. 5). The MS/MS data confirmed a C-S crosslink between Cys1 and Ser5 in all of them (b_5_, −H_2_O, −18 Da), with a distinction at the C-terminus due to the presence of an additional D residue in **1** (y_2_, ‘FD’ residue, Supplementary Fig. 5). Furthermore, the 2D NMR data of purified **2** showed HMBC correlation of H_β_ to C_β_ (*δ*_H_ = 2.96, 3.31 ppm to *δ*_C_ = 35.9 ppm and *δ*_H_ = 3.07, 3.24 ppm to *δ*_C_ = 34.3 ppm) in two Ala residues confirming the thioether ring in Cys1-Ser5 (Supplementary Fig. 6 and Supplementary Table 3). Ciblan A1 (**3**, BGC *cib1*, *Haladaptatus cibarius* JCM19505) and ciblan A2 (**4**, BGC *cib2*, *H. denitrificans* DSM4425) were predicted with a similar modification on the core peptides of CibA1 (ASKLPCCDTTN) and CibA2 (SKLPCCDTTN). The modification includes two bicycle ring structures (Ser2-Cys6 and Cys7-Thr9 in A1, Ser1-Cys5 and Cys6-Thr8 in A2) and one unsaturated amino acid Dhb residue derived from the last Thr residue of each, supported by a series of b ions (b_6_/b_5_ for the first ring and b_9_/b_8_ for the second ring in ciblan A1/A2) and z_2_ fragment (Supplementary Fig. 7). The NMR data of purified **3** exhibited the presence of Dhb residue (C_α_= 131.0 ppm, C_β_ = 130.9 ppm, H_β_ = 6.66 ppm, C_γ_ = 13.4 ppm, H_γ_ = 1.86 ppm) derived from Thr10, along with two thioether rings in Ser2-Cys6 (*δ*_H_ = 2.97, 2.92 ppm of Ala6 to *δ*_C_ = 35.0 ppm of Ala2) and Cys7-Thr9 (*δ*_H_ = 2.98 ppm of Abu9 to *δ*_C_ = 41.2 ppm of Ala7) (Supplementary Fig. 8 and Supplementary Table 3). The stereochemistry of **2** and **3** were confirmed with L configuration in all unmodified amino acids and DL-Lan (2*S*,6*R*) or DL-MeLan (2*S*,3*S*,6*R*) residue in thioether rings (Supplementary Fig. 9 and Supplementary Table 4). Notably, **2** with only six amino acids represents the smallest natural lanthipeptide to date.

Due to their low yield, compounds **5**-**12** were primarily identified using HRMS and biosynthetic analysis. The MS/MS analysis and full reduction-desulfurization study of pellan (**5**, BGC *pel, H. pelagicus* RC-68) confirmed the presence of two dehydrations within residues Thr2, Cys4, Ser7 and Cys12, and proved interwind ring structures in the core peptide of PelA (S<u>T</u>G<u>C</u>Y<u>IS</u>DKPV<u>C</u>I, Supplementary Fig. 10). The partial desulfurization product of archalan γ (**6**, BGC *alnγ*, *H. salinus* YJ-37-H) indicated one methylation on Cys1, residues Ser2, Cys3, and Cys8 reduced to be Ala, and an intact C-S crosslinking in Thr5-Cys9 (<u>CSC</u>Y<u>T</u>RI<u>CC</u>DIDS, Supplementary Figs. 11-15). Subsequent dithiothreitol reduction and MS^n^ analysis determined a disulfide bond between Cys1 and Cys8 and a thioether ring between Ser2 and Cys3 in **6** (Supplementary Figs. 16 and 17). Altogether, archalan γ (**6)** featured N-terminal methylation, a Cys1-Cys8 crosslink, and thioether crosslinks in Ser2-Cys3 and Thr5-Cys9. It expands the number of the smallest thioether ring containing lanthipeptides which only has a few reported cases (such as mersacidin, ltnα, and sacAα)^35^.Archalan α2 (**7**, BGC *alnα*, *H. salinus* YJ-37-H) and medlan B (**8**, BGC *medb*, *H. mediterranei* ATCC33500) contained two thioether rings, supported by their key fragments of b_4_ (-H_2_O, C-S crosslink in Cys1-Thr4) and b_9_ (−2H_2_O, another crosslink in Cys5-Ser9) in MS/MS analysis (Supplementary Fig. 3), in line with the previously reported structure of archalan α^26^. Archalan β1-2 (**9** and **10**, BGC *alnβ*, *H. salinus* YJ-37-H) and litlan A1-2 (**11** and **12,** BGC *lit*, *H. litoreus* JCM31109) exhibited similar crosslinks to the previously reported archalan β^26^, featuring two interwind thioether rings that resulted in MS/MS fragments from the N/C-terminal overhang amino acids outside of the rings (Supplementary Figs. 4 and 18).

The chemical structures of mature archaeal lanthipeptides have a high appearance of acidic residues which may facilitate adaptation to high-cation cellular environments and maintain a stable compound structure to function efficiently in high-salt surroundings. Moreover, archaeal lanthipeptides revealed that they are typically shorter than bacterial lanthipeptides, such as nukacin ISK-1 and haloduracins (Fig. 2d). Despite being short in length, their bicycle and interwind crosslinking patterns and modifications including disulfide bond, methylation, dehydration, hydroxylation, are as intricate as those found in bacterial lanthipeptides.

### Discovery of unique lanthipeptides featuring unprecedented maturation

Apart from the classical lanthipeptides mentioned above, several unusual peptide signals from BGCs *meda* and *lar* engrossed our attention for the unexpected modification (Fig. 3 and Supplementary Figs. 19-29). They were detected in the extracts of both wild-type and recombinant strains, suggesting that they were likely encoded by these two BGCs, albeit with unexpected modifications as revealed by MS/MS data. While the MS analysis of classical lanthipeptides allows for the accurate prediction of core peptide sequences and putative modifications, the MS/MS data of these peptide signals only provided short amino acid fragments without consistent patterns, such as ‘FD’ fragment from the C-terminus of LarA (CPTC<u>DF</u>), ‘Y’ residue also from C-terminus and ‘S’ residue from N-terminus (<u>S</u>TCT<u>Y</u>) (Fig. 3b and Supplementary Fig. 24).

**Figure 3.**
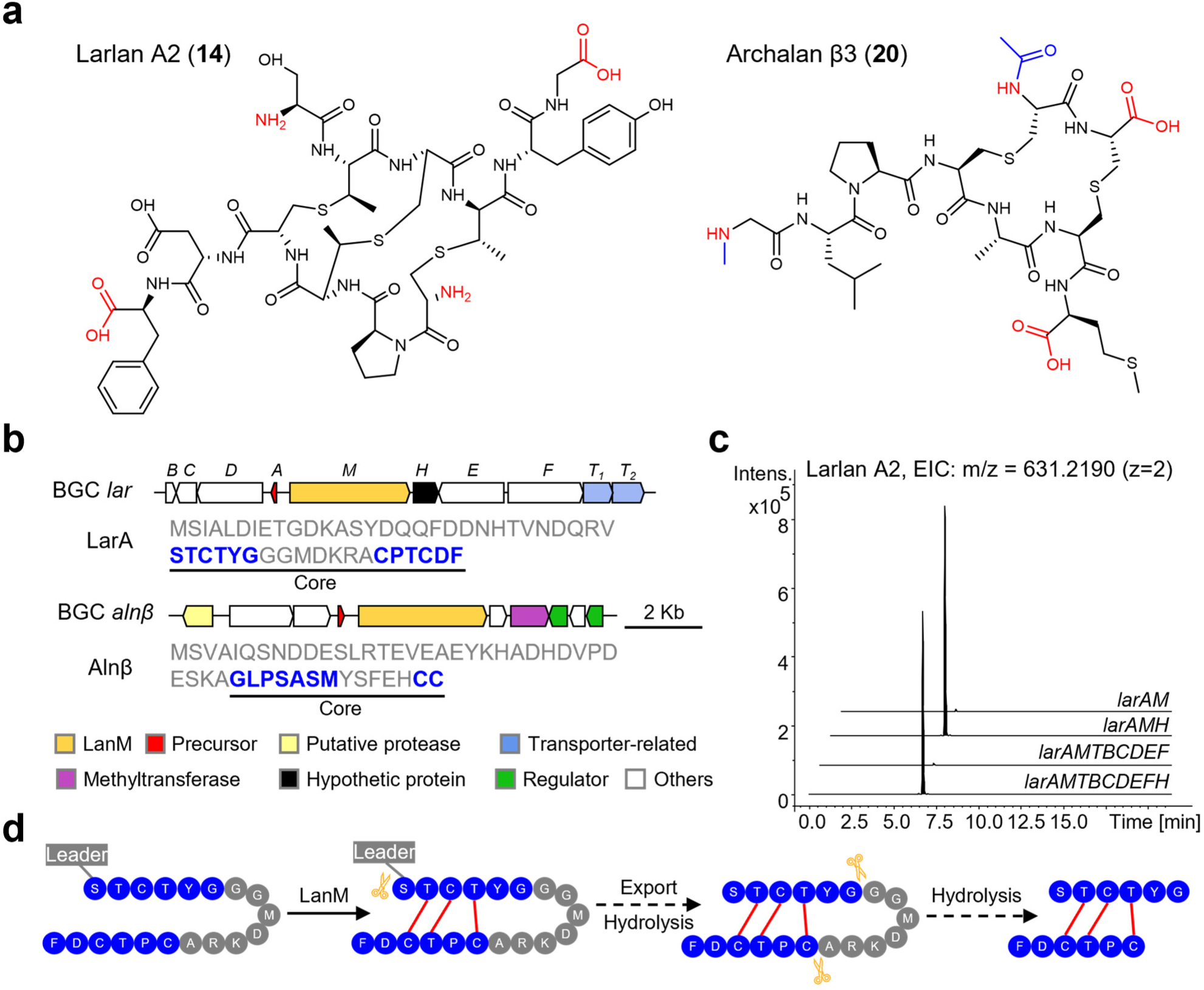
Novel chemistry and biosynthesis of archaeal lanthipeptides. **a**, Chemical structure of DADC lanthipeptides. Diamino-dicarboxylic termini are in red, and specific modifications at the N-terminus are in blue. **b**, BGCs *lar* and *alnβ* with their corresponding precursors. **c**, LC-MS analysis of in vivo reconstitution of BGC *lar* with/without gene *larH*. **d**, Proposed biosynthetic pathway of DADC lanthipeptides, demonstrated by larlan A2.

To fully characterize their structures, 3.2 mg of **13** ([M+3H]^3+^ = 415.4769, calculated [M+3H]^3+^ = 415.4772, Δ = 0.7 ppm) from the wild-type strain, 3.5 mg of **14** ([M+2H]^2+^ = 631.2194, calculated [M+2H]^2+^ = 631.2198, Δ = 0.6 ppm), and 2.0 mg of **15** ([M+2H]^2+^ = 652.2301, calculated [M+2H]^2+^ = 652.2251, Δ = 7.7 ppm) from the recombinant strain were purified, which were named medlan A1 (**13**), larlan A2 (**14**) and larlan A5 (**15**), respectively. Further NMR spectroscopy analysis revealed that these lanthipeptides comprised two parallel but opposite-direction peptide chains crosslinked by thioether bonds, characterizing them as a novel lanthipeptide subfamily (Supplementary Figs. 20-22 and Supplementary Table 3). These crosslinked patterns expose diamino-dicarboxylic termini, leading to the name of DADC lanthipeptides, with certain amino acids within the thioether rings being partially cleaved (Fig. 3). In addition, medlan A2 (**16**), larlans A1 (**17**), A3 (**18**), and A4 (**19**) are analogs of medlan A1 (**13**) or larlan A2 (**14**) and have been deduced to exhibit longer or shorter overhang amino acids at the N/C terminus through LC-MS/MS analysis, possibly due to hydrolysis by unknown aminopeptidase (Supplementary Figs. 23-29). As for three MeLan residues in medlans and larlans increasing the complexity of stereochemistry configuration, two additional single mutations of Thr to Ser in precursor LarA were constructed. By purifying 0.3 mg of each mutant (T2S and T4S of larlan A2) for the advanced Marfey’s analysis, a clear stereo structure annotation of each MeLan residue was achieved when compared with four MeLan standards^39,40^ (Supplementary Fig. 9 and Supplementary Table 4). The thioether rings between Thr2 and Cys10, Thr9 and Cys3 are characterized in the LL-MeLan (2*R*,3*R*,6*R*) configuration, while another ring between Thr4 and Cys7 in DL-MeLan (2*S*,3*S*,6*R*) configuration in larlan A2 (Fig. 3).

To find out whether other lanthipeptides also employ a similar maturation strategy, the LC-MS/MS data of both wild-type and recombinant strains were meticulously analyzed. The metabolic profile of *H. salinus* YJ-37-H, harboring six lanthipeptide BGCs, displayed mass signals with a fragmental pattern resembling larlans and medlans (Supplementary Fig. 30). To characterize their structures, 1.0 mg of compound **20** ([M+2H]^2+^ = 453.6800, calculated [M+2H]^2+^ = 453.6796, Δ = 0.9 ppm) was obtained for NMR and advanced Marfey’s analysis (Supplementary Figs. 31 and 32). As anticipated, archalan β3 (**20**), derived from BGC *alnβ*, shares the same characteristics as DADC lanthipeptides. Other derivatives, archalans β4-7 (**21**-**24**), were also detected and elucidated by LC-MS/MS with additional modifications or overhang amino acids (Supplementary Figs. 33-36). The advanced Marfey’s analysis revealed that two Lan residues are LL (2*R*,6*R*) configurations, and other unmodified amino acids are proteinogenic L configurations in archalan β3 (Fig. 3, Supplementary Fig. 9 and Supplementary Table 4).

Identifying this particular lanthipeptide subfamily exclusively in Archaea highlights the existence of a unique biosynthetic pathway and enzymes that are distinct from those in bacteria. Synteny analysis^41^ of the DacA family revealed the presence of three conserved genes: *A*, *M*, and *H*, which might serve as the boundary of DADC lanthipeptide BGCs (Supplementary Fig. 37 and Supplementary Data 3). The *lar* BGC was then selected for the in vivo reconstitution study which resulted in the production of a trace amount of larlan A2 when co-expressed genes *larA* and *larM* (Fig. 3b, c and Supplementary Fig. 37). This observation suggested the potential presence of generic peptidases from the heterologous host. On the contrary, a significantly increased production of larlan A2 was observed when co-expressed with the unannotated gene *larH* (Fig. 3c). This implied that LarH may play a role as a regulator, cofactor, chaperon, or a yet unidentified peptidase involved in the biosynthesis of larlan A2. In another case, we observed the production of classical lanthipeptide archalan β1-2 but didn’t detect DADC lanthipeptide archalan β3-7 through heterologous expression, despite including the putative peptidase within the BGC (Supplementary Figs. 4 and 37). Although both classical and DADC lanthipeptides were identified in the wild-type strain, heterologous expression only yielded classical lanthipeptides of archalan β1-2, supporting the hypothesis that putative proteases outside the BGC are involved in biosynthesis and suggesting a complex biosynthetic pathway for DADC lanthipeptides. This pathway is predicted to include additional hydrolysis steps compared to classical lanthipeptides, facilitated by unidentified generic proteases to cleave amino acids within the thioether rings (Fig. 3d).

### Archaeal lanthipeptides exhibit anti-archaeal and motility regulation activities

Given the chemical diversity and novelty of archaeal lanthipeptides, we next wanted to investigate their ecological functions. Six lanthipeptide crude extracts obtained from heterologous expression and seven purified lanthipeptides were collected for antimicrobial activity screening against nine haloarchaeal and six bacterial strains (Fig. 4). To exclude the potential influence of halocin produced by the heterologous host^42^, the same gradient fractions from the host containing an empty vector were tested for comparison. Specifically, extracts from the heterologous expression of BGCs *medb, cib2,* and *alnα-γ* exhibited significant anti-haloarchaeal activity at 100 μg mL^-1^, even though the lanthipeptides were in low proportion (Fig. 4a). The mature products from BGCs *alnα* and *medb*, archalan α (**25**), archalan α2 (**7**) and medlan B (**8**), were analogs exhibiting a similar anti-haloarchaeal spectrum (Fig. 4a). The purified lanthipeptide, archalan α (**25**), demonstrated strong inhibition against *H. larsenii* JCM 13917, *Haladaptatus cibarius* DSM 19505, *H. amylolyticus* JCM 18367, and *Halomicroarcula salina* JCM 18369 at 100 μg mL^−1^ (Fig. 4b). Notably, the MIC value of **25** against *H. salina* JCM 18369 was 6.25 μg mL^−1^ (Supplementary Fig. 38). On the contrary, the DADC lanthipeptide archalan β3 (**20**) exhibited lower activity than the classical lanthipeptide archalan β1-2 (**9**, **10**) produced by heterologous expression of BGC *alnβ*. The specific anti-haloarchaeal activity observed in archaeal lanthipeptides may be attributed to the characteristics of their habitats, which predominantly host Haloarchaea.

**Figure 4.**
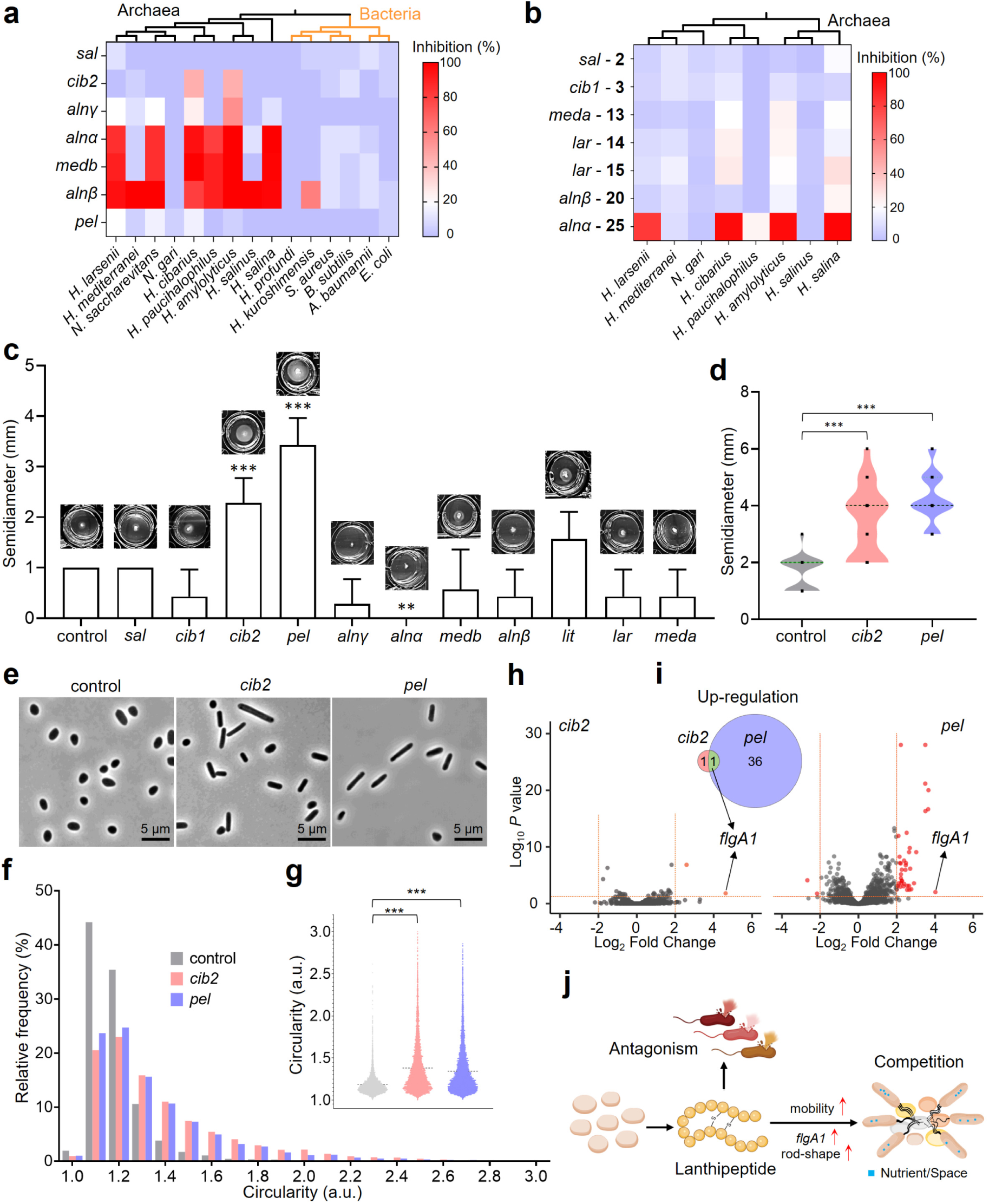
Ecological functions of archaeal lanthipeptides. The antimicrobial activity of the fractionated crude extracts of recombinant strains with different BGCs (**a**) and purified archaeal lanthipeptides at 100 μg mL^−1^ (**b**). Compounds **2**, **3**, **13**, **14**, **15**, **20**, and **25** correspond to sallan A2, ciblan A1, medlan A1, larlan A2, larlan A5, archalan β3, and archalan α, which are purified lanthipeptides encoded by the BGCs labeled in front of them. **c**, The motility assay screening of *H. volcanii* H1424 with empty vector (control) or different lanthipeptide BGCs in 24 well plates with a diameter of 1.56 cm/well. *** *P* < 0.001, ** *P* < 0.05. Each group has three biological replicates. Bars show the mean ± standard deviation (SD), n = 7. **d**, The violin plot of the motility assay of *H. volcanii* H1424 with empty vector, BGCs *cib2* and *pel*, respectively. The median and quartile ranges are shown as horizontal lines in black and green respectively. Black dots are the aligned experiment data, *** *P* < 0.001, n = 21. Each group has six biological repeats. **e**, Phase-contrast images of *H. volcanii* H1424 with empty vector, BGCs *cib2* and *pel*, respectively. Scale bars are 5 μm. Cell circularity (**g**) and frequency distribution (**f**) were calculated from the phase-contrast microscopy of recombinant strains. Each group has three biological replicates. Lines display the means of each group (n = 5,690, 5,553, and 6,264 for control, *cib2*, and *pel* groups, respectively), *** *P* < 0.001. **h**, Volcano plots illustrate the differential expression genes (DEGs) in strains with BGC *cib2* (left) or *pel* (right) compared to the strain with an empty vector. **i**, Venn diagram depicting the overlap DEGs detected in both strains. **j**, A proposed mode for the ecological function of archaeal lanthipeptides in antagonistic interaction and nutrient/space competition.

To study the in vivo function of lanthipeptide BGCs, stab-inoculation screens were conducted to assess their impact on the host’s motility. Surprisingly, three recombinant strains were found to modulate the motility as either activator or inhibitor (Fig. 4c). Strains containing BGCs *cib2* and *pel*, responsible for the production of ciblan A2 (**4**) and pellan (**5**), exhibited a significantly larger motility zone than the control (Fig. 4c, d, Supplementary Fig. 39 and Supplementary Data 4). Subsequently, strain with BGC *cib2* was selected for time-course motility assays to gain a deeper insight, indicating that BGC *cib2* significantly activated the host’s motility zone in the beginning without affecting growth and all strains displayed the Bull’s eye pattern in the late stage (Supplementary Fig. 39). Conversely, BGC *alnα*, responsible for the production of archalan α (**25**), inhibited the motility may be attributed to its antagonistic activity, which could impede host’s growth (Fig. 4b, c, and Supplementary Fig. 39). No obvious motility regulation assay in BGC *cib1*, in contrast to BGC *cib2* with identical precursor peptides, may be attributed to structure difference of their key products (Fig. 2 and Supplementary Fig. 4). These findings point to a new ecological role of lanthipeptides, potentially reducing the host’s reaction time or increasing that of other strains in response to stimuli, by regulating motility activity^23,24,43^.

A comprehensive investigation combining morphological and transcriptomic analysis was conducted to elucidate the mechanisms by which archaeal lanthipeptides enhance motility. Microscopic analysis of recombinant strains demonstrated that lanthipeptides could elevate the frequency of rod-shaped cell morphology, a characteristic associated with hypermotility^23,24,43^ (Fig. 4 e-g and Supplementary Data 5). Additionally, the transcriptomic analysis revealed that the archaellin gene *flgA1*, a major flagellin gene responsible for the motility of *H. volcanii*^44^, was upregulated in strains possessing BGCs *cib2* or *pel* (Fig. 4h, i and Supplementary Data 6). These studies support the hypothesis that lanthipeptides may promote motility by inducing a rod-shaped cell morphology and upregulating the expression of the archaellin gene *flgA1*. These lanthipeptide-mediated regulations could potentially confer advantages to their host in environmental adaptation, such as occupying space, enhancing nutrient acquisition, and evading predation (Fig. 4j).

## Discussion

Archaea, which predominantly survive in extreme environments, are reported to have unique metabolic pathways with eco-evolutionary strategies distinct from bacteria and fungi^2,6^. A notable portion of archaeal BGCs consists of RiPPs and terpenes^26^, as identified *in silico*, contrasting with the prevalent NRPs and PKs BGCs in bacteria^45^. However, given the lack of identified SMs, our understanding of archaeal chemical language, remains markedly limited compared to our comprehensive knowledge of bacteria and fungi, particularly regarding their biosynthesis and ecological roles^46,47^. While RiPPs exhibit a high diversity and widespread distribution in archaea, class II lanthipeptides have been found to exist exclusively in halophilic archaea^26–28^. This observation implies archaeal lanthipeptides to be niche-specific metabolites adapted to hypersaline environments, as evidenced by the amino acid composition bias and phylogenetic analysis of the hallmark enzyme LanMs and their corresponding precursor LanAs in this study. We then employed a combined strategy involving the biosynthetic rule-based metabolomic analysis and the first use of heterologous expression of archaeal BGCs in Haloarchaea to unveil the chemical landscape of archaeal lanthipeptides. The heterologous expression approach substantially expands the repertoire of archaeal SMs and encourages further exploration of other unique and silent BGCs from the domain of Archaea, such as diverse and widely distributed YcaO/rSAM-modified RiPPs and NRPs. Our discovery of diverse lanthipeptides from archaea expedites the discovery and biosynthesis study of archaeal chemical languages, laying the stage for their chemical ecology study.

The discovery of a new lanthipeptide subfamily, DADC lanthipeptides, underscores the distinct characteristics of archaeal metabolic pathways and enzymes. These lanthipeptides are marked by the unprecedented diamino-dicarboxylic termini, resulting from the unknown cleavage of amino acids within the thioether rings. We propose that DADC lanthipeptide undergoes a multiple-step maturation process, initially forming classical lanthipeptides and then cleaving amino acids within the thioether rings through a generic protease outside the BGCs. This hypothesis is corroborated by the findings from the in vivo reconstitution of BGCs *lar* and *alnβ* and their corresponding natural products. The precise biosynthetic pathways and enzymes of DADC lanthipeptides remain elusive due to the challenges encountered in the enzymatic study of Haloarchaea. Nonetheless, our research sheds light on the complexity and novelty of archaeal SMs, emphasizing the untapped potential of archaea in natural products and enzyme discovery.

To gain first insight into the ecological function of archaeal lanthipeptides, we assessed in vitro antimicrobial tests and in vivo motility assay. Their specific antagonism against haloarchaea may stem from the prevalence of haloarchaea over normal bacteria in halophilic habitats^48^. However, it is important to note that this study, with a panel of limited indicators, potentially overlooks antagonistic interactions with other archaeal or bacterial species. Subsequently, we carried out the motility regulation assay, which is believed to contribute to nutrient acquisition, reproductive efficiency, antibiotic resistance, and evasion of predation^23,24,43^. The discovery that lanthipeptide BGCs *cib2* and *pel* were involved in activating motility reveals a previously unknown ecological function of RiPPs. This newfound role may assist the host organism in adapting to environmental stimuli. Notably, our microscopic and transcriptomic analyses provide evidence that lanthipeptides enhanced the host’s motility by inducing a shift in cell shape towards a rod-like morphology and upregulating the expression of archaellin gene *flgA1*. These findings underscore the potential role of lanthipeptides in influencing two crucial factors for archaeal motility^23,24,44^, despite our limited understanding of regulation mechanisms due to the difficulties in genetically manipulating the wild-type host.

In conclusion, this study presents the first heterologous expression of archaeal BGCs to reveal the chemical landscape of lanthipeptides in archaea. Furthermore, we have demonstrated a newfound ecological function of archaeal lanthipeptides in regulating in vivo motility, with the mechanism involving the transformation of rod-shaped cell morphology and the upregulation of archaellin expression. Our discovery of biofunctional lanthipeptides from Archaea provides new insights into their ecological functions, which are anticipated to spur research into the less-explored field of archaeal chemical biology and chemical ecology. These findings not only signify a major advancement in decoding the archaeal chemical language but also highlight the potential of archaea as a rich source of bioactive SMs and unique enzymes.

## Methods

### BGC analysis

Using antiSMASH bacterial version 7.0^33^, archaeal BGCs were identified from the archaeal genomes available on NCBI RefSeq and Genbank databases (accessed in Aug. 2023). The genomes were initially deduplicated based on 99% average nucleotide identity using dRep v3.4.5^49^. The completeness and contamination of genomes were assessed using CheckM2^50^, and genomes that met the quality requirements (completeness >80% and contamination <5%) were selected for further analysis. Representative genomes were then selected prioritizing higher assembly levels and genome size. The BGC class was determined based on the product annotation provided by antiSMASH. BGC features were extracted using BiG-SLiCE v2.0.0 with default settings^51^. The similarity of archaeal BGCs to 2,502 BGCs from MiBiG 3.1^34^ was quantified by calculating the minimal cosine distance. Pairwise cosine distances between archaeal BGCs or between archaeal BGCs and MiBiG BGCs were computed using the SciPy library in Python 3.8^52^. Based on those cosine distances, hierarchical clustering with average linkage was adopted for clustering BGCs into GCCs at a distance cutoff of 0.8 using Scikit-learnv1.0.2^53^. The heatmap showing BGC distribution was plotted using *Complex Heatmap* package in R 4.1.2^54^. The BGC synteny was analyzed by Clinker^55^ with default parameters.

### Sequence similarity network analysis of archaeal LanAs

The SSN analysis was conducted as in our previous study^26^. Archaeal class II lanthipeptide BGCs were extracted according to the product annotation from antiSMASH. The precursor peptides of class II lanthipeptide, LanAs, were predicted from the ten open reading frames adjacent to *lanM*, with lengths ranging from 10 to 100 amino acids and containing Cys and Thr/Ser residues within 30 amino acids of the C-terminus. A total of 354 LanAs was submitted to the EFI - Enzyme Similarity Tool^56^ with an *E*-value of 1.0 × 10^−1^ and an Alignment Score Threshold of 7. Sequences with 100% identity were grouped into a single node, leading to 196 unique sequences. The resulting SSN was visualized using Cytoscape 4.1^57^.

### Phylogenomic and amino acid composition analysis of archaeal LanAs and LanMs

A total of 12,398 bacterial and 110 archaeal LanMs (Supplementary Data 2) were firstly deduplicated at 80% similarity using MMseqs^58^ with the following parameters: -c 0.8 --min-seq-id 0.8 -e 10 -- cluster-reassign 1. Alignments were then performed using MUSCLE^59^ and visualized with FastTree^60^ using default parameters. LanA sequences were aligned using MAFFT v7^61^ with parameter: -- maxiterate 1000. The phylogenomic trees were conducted by IQ-TREE v2^62^ with the following parameters: -m LG+G4 -B 1000 -T 4. The visualization of phylogenetic trees of archaeal LanAs and LanMs was done using iTOL v6^63^. The co-phylogenetic analysis of archaeal LanAs and LanMs was performed using Phytools 2.0^64^. To compare the compositional bias of each amino acid in archaeal and bacterial lanthipeptides at protein and substrate levels, the amino acid composition of LanAs and LanMs was calculated through an in-house Python v.3.12.7 script (https://github.com/GAOYingHKU/Binomial-test-Z-score/) and plotted with ggplot2 (v.3.5.1)^36,65^. *Z*-scores were calculated relative to archaeal LanAs and LanMs, where *Z* > 2.0 indicated significant enrichment of a given amino acid in archaea (’Over-represented’), *Z* < −2.0 indicated significant depletion (’Under-represented’), and some amino acids showed no significant biases (’NS’).

### Cultivation of haloarchaea strains and extraction of lanthipeptide metabolites

All the wild-type strains listed in Supplementary Table 1 were cultured in modified DSMZ medium 1520 (NaCl 250.0 g L^-1^, KCl 3.0 g L^-1^, trisodium citrate 3.0 g L^-1^, MgSO_4_.7 H_2_O 20.0 g L^-1^, yeast extract (Difco) 1.0 g L^-1^, casamino acids (Difco) 7.5 g L^-1^, glucose 1.0 g L^-1^, Na-succinate 10.0 g L^-1^, sucrose 20.0 g L^-1^) on a rotary shaker (37 °C and 200 r.p.m.). After cultivating for 2 days, 10% cell culture was inoculated to a large volume fermentation at 37 °C and 200 r.p.m. for another 5-7 days. The recombinant strains were cultured in complete Hv-YPC medium (yeast extract 5 g L^-1^, peptone 1.0 g L^-1^, casamino acids 1.0 g L^-1^, NaCl 160.0 g L^-1^, MgCl_2_·6H_2_O 20.0 g L^-1^, MgSO_4_·7H_2_O 23.0 g L^-1^, KCl 4.7 g L^-1^, 13.3 mM pH 7.5 Tris-HCl and other 3 mM CaCl_2_ adding after cooling) or selectivity Hv-Ca medium ( casamino acids 17.0 g L^-1^, NaCl 160.0 g L^-1^, MgCl_2_·6H_2_O 20.0 g L^-1^, MgSO_4_·7H_2_O 23.0 g L^-1^, KCl 4.7 g L^-1^, 13.3 mM pH 7.5 Tris-HCl, and other 3 mM CaCl_2_, thiamine 0.9 μg L^-1^ and biotin 0.1 μg L^-1^ adding after cooling) as previously reported^37,38^. The single colonies were initially cultivated in Hv-Ca medium at 45 °C for 24 h, and then 5 mL of the suspension was inoculated to a 250 mL flask containing 80 mL Hv-YPC medium. Culture broth of each strain was collected at −80 °C after 3 - 7 days of fermentation. For lanthipeptide extraction, 1 mL culture broth was extracted by Diaion^®^ HP-20 resin (Sigma-Aldrich), followed by washing three times with 1 mL of ddH_2_O and eluting with 0.5 mL MeOH. The resulting extracts were dried by a vacuum concentrator, redissolved in MeOH, and analyzed by HR LC-MS as detailed in Supplementary Information.

### Heterologous expression and reconstitution of lanthipeptide BGCs in *H. volcanii* H1424

Lanthipeptide BGCs were amplified from the genomic DNA of respective wild-type strains using primers designed with appropriate overhangs (Supplementary Table 3). The corresponding PCR products were inserted into the plasmid pTA1228 via Gibson assembly. The recombinant vectors were first propagated in *E. coli* DH5α (DE3) and then introduced into *H. volcanii* H1424 via PEG-mediated transformation approach guided by the Handbook^66^. The recombinant *H. volcanii* H1424 strains were selected in Hv-Ca agar plates and single colonies were grown in Hv-YPC liquid medium for 2 days (45 °C and 200 r.p.m.).

### Structure elucidation of archaeal lanthipeptides

^1^H, ^13^C, ^1^H-^1^H COSY, ^1^H-^13^C HSQC, ^1^H-^13^C HMBC, TOCSY, and NOESY NMR spectra for medlan A1 (**13**), larlan A2 (**14**), larlan A5 (**15**) and archalan β3 (**20**) were recorded on Avance DRX 600 FT-NMR spectrometer (600 and 150 MHz for ^1^H and ^13^C NMR, respectively) using DMSO-*d*_6_. Chemical shifts were reported using the DMSO-*d*_6_ resonance as the internal standard for ^1^H-NMR DMSO-*d*_6_: *δ* = 2.50 p.p.m. and ^13^C-NMR DMSO-*d*_6_: *δ* = 39.6 p.p.m. Sallan A2 (**2**) and ciblan A1 (**3**) used MeOH-*d*_4_ as the solvent and the internal standard were referenced to: ^1^H-NMR MeOH-*d*_4_: *δ* = 3.31, 4.87 p.p.m. and ^13^C-NMR MeOH-*d*_4_: *δ* = 49.0 p.p.m.

### Motility assay

The motility screening for recombinant strains was set up according to previously reported methodology^23,24^. Briefly, 2 μL of mid-log phase strain cultures (OD_600_ = 0.1, 50 μL in 96-well plates of Corning, Thermo Scientific Varioskan Flash) were stabbed into the plates containing modified Hv-Ca + 0.4% agar motility medium supplemented with additional trace elements ( per liter: 5.0 g EDTA, 0.8 g FeCl_3_, 0.05 g ZnCl_2_, 0.01 g CuCl_2_, 0.01 g CoCl_2_, 0.01 g H_3_BO_3_, 1.6 g MnCl_2_, 0.01 g Ni_2_SO_4,_ and 0.01 g H_2_MoO_4_). The plates were incubated in a sealed plastic bag at 45 ℃ for 1 day. The time-course motility assays were then conducted at 45 ℃ for 2 - 4 days. The containers used were 6/12/24 - well plates (NEST, China), with approximately 8/4/2 mL of medium added per well, respectively.

### Light microscopy

Phase-contrast microscopy was performed using a 100x oil immersion objective lens (Zeiss, LSM980 system, Germany) to examine the cell morphology of three recombinant strains (containing the empty vector pTA1228, pTA1228-BGC *cib2* or pTA1228-BGC *pel*) after 15 hours of cultivation in a shaker (45 °C, 200 r.p.m). Specifically, 1 μL of each strain was placed onto slides coated with a 1% agarose pad containing 18% BSW, and a coverslip was used to cap the samples. Each group has three biological replicates to ensure consistency. ImageJ software^67^ was used to distinguish and measure the perimeter and area values of different cells (Supplementary Data 4). The circularity values were calculated using the formula:

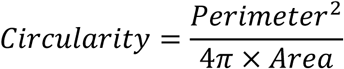

For the tolerance of ImageJ, the circularity values were filtered no greater than 3.0 and no less than 1.0. The frequency of circularity values was then visualized using GraphPad Prism 8 software.

### Transcriptomic analysis

The cell cultures of recombinant strains carrying the empty plasmid pTA1228, pTA1228-BGC *cib2* or pTA1228-BGC *pel* in Hv-Ca medium were harvested after 20 hours of incubation (45°C and 200 r.p.m.). Each group has three biological replicates. The total RNA of all the samples was extracted using RNAprep Pure Cell/Bacteria Kit (Tiangen Biotech, Beijing, China) according to the manufacturer’s instructions. RNA qualification was examined by 1% agarose gel electrophoresis and Aligent 5400 (Agilent Technologies, Santa Clara, CA, USA). RNA libraries were constructed using the Fast RNA-seq Lib Prep Kit V2 (ABclonal, Cat. RK20306) and sequenced with Illumina nova X Plus. Sequence reads were trimmed for adaptor sequence, low-quality sequence and rRNA sequence using kneaddata v0.12.0 (https://github.com/biobakery/kneaddata) with parameters: -db SILVA_128_LSUParc_SSUParc_ribosomal_RNA --trimmomatic-options ILLUMINACLIP:TruSeq3-PE.fa:2:30:10 SLIDINGWINDOW:4:20 MINLEN:60 --run-trim-repetitive --reorder --bowtie2-options ‘--very-sensitive --dovetail’ --bypass-trf. Trimmed sequence reads were mapped to the reference genome sequence of *H. volcanii* (GCF 010692905.1) along with GTF annotations using Bowtie 2.3.5.1 with default parameters^68^. The differential expression genes (DEGs; *p* < 0.05, absolute regulation fold change ≥ 2) between control and experiment groups were analyzed with DESeq2 (version 1.38.3)^69^. The DEGs were depicted through the utilization of volcano plots using the EnhancedVolcano in R 4.2.1.

## Ethics approval and Consent to participate

Not applicable.

## Consent for publication

Not applicable.

## Data availability

All genomes used in this research were obtained from the NCBI Assembly RefSeq database (https://www.ncbi.nlm.nih.gov/assembly) and Genebank database (https://www.ncbi.nlm.nih.gov/genbank/). The HRMS data generated in this study have been deposited in MassIVE under the accession code MSV000095989. The transcriptomic data of this study has been deposited in NCBI with the accession number GSE281021. Data supporting the results of this study is available in the paper, as well as supplementary information and supplementary data.

## Code availability

The code used in this study for amino acid composition bias analysis of LanAs and LanMs assigned a citable DOI through Zenodo (DOI10.5281/zenodo.13705935) and can be accessed at https://github.com/GAOYingHKU/Binomial-test-Z-score/.

## Competing interests

The authors declare no competing interests.

## Funding

This work is partially funded by a Shenzhen Basic Research General Programme (JCYJ20210324122211031), the Research Grants Council of Hong Kong (27107320 and 17115322), and the Hong Kong Branch of Southern Marine Science and Engineering Guangdong Laboratory (Guangzhou) (SMSEGL20SC02) to Y.-X.L. The funding sources had no role in study design, data collection and analysis, decision to publish, or manuscript preparation.

## Author contributions

Z.-M.S. and Y.-X.L. designed the research and prepared the manuscript. Z.-M.S., C.C., H.L., J.Z. and Q.Z. performed research. Z.-M.S. and Y.-X.L. characterized the structures of compounds. Z.-M.S. and C.C. performed the LC-MS analysis. Z.-M.S., Y.G., X.L., D.Z., and G.W. did the genomic analysis. Y.G. analyzed the transcriptomic data. Z.-M.S., Y.Q., and H.J. performed the phase-contrast microscopic analysis. Z.-M.S., C.C., Y.G., X.L., Y.Q., D.Z., G.W., H.L., Q.Z., J.Z., P.Y. C., H.J. and Y.-X.L. analyzed data and discussed the results. W.L. and Y.-X.L. founded the research. Y.-X.L. supervised the research. All authors read and approved the final manuscript.

## Acknowledgments

The authors would like to express their gratitude to Prof. Thorsten Allers (School of Life Sciences, University of Nottingham, UK) and Prof. Likui Zhang (College of Environmental Science and Engineering, Yangzhou University, China) for their generous sharing of the plasmid pTA1228, host strain *H. volcanii* H1424 and corresponding protocol. Thank Prof. Shihui Dong (College of Chemistry and Chemical Engineering, Lanzhou University, China) for kindly providing MeLan and Lan standard samples and Yi-Man Eva FUNG, Jo Yip, and Bonnie Yan from the Department of Chemistry, University of Hong Kong for their help in MS and NMR analysis. Thank the microscopic imaging service at the Centre for PanorOmic Sciences (University of Hong Kong) and Zhuohan Li (Department of Chemistry, University of Hong Kong) for their help in microscopic analysis. Some figures were created with BioRender.com.

## Notes

### Competing Interest Statement

The authors have declared no competing interest.

### Summary of Updates

A comprehensive investigation combining morphological and transcriptomic analysis was conducted to elucidate the mechanisms by which archaeal lanthipeptides enhance motility.

