## Supplementary Figures and Supplementary Tables for "Decoding the chemical language of RiPPs from the untapped Archaea domain"

### Experiment materials and methods

#### General materials and experimental procedures

Biochemicals and media components for bacterial cultures were purchased from standard commercial sources. Marfey's reagent *N*α-(5-fluoro-2,4-dinitrophenyl)-L-leucinamide (L-FDLA) and *N*α-(5-fluoro-2,4-dinitrophenyl)-D-leucinamide (D-FDLA) were purchased from the Tokyo Chemistry Industry. DNA polymerase Phanta Max Master Mix (Vazyme) was used for the polymerase chain reactions and NEBuilder® HiFi DNA assembly (NEB) was used for Gibson assembly. Nisin was purchased from Sigma (CAS: 1414-45-5, potency ≥ 900IU/mg). All strains used in this study are listed in Supplementary Table 1. 1D and 2D NMR spectra were recorded at 298K on a Bruker Avance DRX 600 FT-NMR spectrometer (600 and 150 MHz for $^{1}$H and $^{13}$C NMR, respectively). NMR solvents dimethyl sulfoxide-$d_6$ (CAS: 2206-27-1, purity: 99.5 %) and Methonal-$d_6$ (CAS:811-98-3, purity: 99.8 %) were purchased from Cambridge Isotope Laboratories Inc. (USA) and Aladdin (Shanghai, China), respectively.

High-resolution liquid chromatography-mass spectrometry (HR LC-MS) analyses were performed on UltiMate 3000 UHPLC Systems with a Waters Acquity UPLC BEH C18 column (1.7 µm, 300 Å, 2.1 × 150 mm) coupled to Bruker impact™ II Mass Spectrometer except for the heterologous expression analysis of DADC lanthipeptides with 100 mm column. The column was maintained at 40 °C and ran at a 0.2 mL min $^{-1}$ flow rate, using 0.1% formic acid in $H_2O$ as solvent A and 0.1% formic acid in ACN as solvent B. A gradient was employed for chromatographic separation starting at 5% B for 2 min, then 5% to 95% B for 15 min, washed with 95% B for 4 min and finally held at 5% B for 1 min. All the samples were analyzed in positive polarity, using data-dependent acquisition mode. The HR LC-MS data was analyzed with Bruker Compass DataAnalysis 4.3. Low-resolution (LR) LC-MS analyses were performed on the Waters ACQUITY H-Class UPLC system coupled with an ACQUITY SQ detector 2 mass spectrometer. Separations were carried out on the same column and methods as HR LC-MS unless otherwise stated.

#### Production, identification, and purification of archaeal lanthipeptides

Sallan A1-A2 (**1**, **2**)

The sallans were only produced by the heterologous expression of BGC *sal* in *H. volcanii* H1424 at 45 °C. The recombinant H1424 culture broth was collected after cultivation in Hv-YPC medium for 3 days at 45 °C and extracted overnight by 10% HP-20 resin (v/v). The resin was washed three times for desalting, followed by elution with MeOH to collect the crude extracts. This extraction method applies to all subsequent crude extracts, unless otherwise specified. The crude extract was evaporated and injected into an HPLC system (Waters, Parsippany, NJ, United States) equipped with a Phenomenex Kinetex XB-C18 column (250 mm × 10 mm, 5 µm, 100 Å) and eluted by isocratic 16% ACN/$H_2O$ containing 0.05% trifluoroacetic acid (TFA) at the flow rate of 4 mL min $^{-1}$. Sallan A1 (**1**) and A2 (**2**) appeared at 30.8-33.5 and 36.0-38.0 min respectively.

Ciblan A1 (**3**)

Ciblan A1 (**3**) was produced and extracted from the fermentation broth of the wild-type strain *H. cibarius* DSM 19505 after being cultivated in modified DSMZ medium 1520 for 5 days at 37 °C. The isolation of ciblan A1 (**3**) was eluted with Phenomenex Biphenyl column (250 mm × 10 mm, 5 µm, 100 Å) with a gradient elution of 10% to 14% ACN/$H_2O$ containing 0.05% TFA in 5.0-45.0 min at the flow rate of 3 mL min$^{-1}$. Ciblan A1 (**3**) appeared at 19.9-23.0 min.

Medlan A1 and A2 (**13**, **16**)

For larger-scale fermentation, wild-type strain *H. mediterranei* was cultured in modified DSMZ medium 1520 for 3 days at 37 °C. The crude extract passed through an open ODS (Octadecylsilane) column, eluting with a gradient concentration of MeOH-$H_2O$ (10-50% at intervals of 10%, and finally 100%). Following this, the fraction with target compounds detected by LC-MS analysis was evaporated and loaded to a Sephadex LH-20 column (Solarbio, China) using a solvent mixture of 60% MeOH/$H_2O$ system to separate and concentrate medlan A1-2 (**13**, **16**). The purification of melan A1-2 (**13**, **16**) was on an HPLC system with Phenomenex Biphenyl column (250 mm × 10 mm, 5 μm, 100 Å) with 40% MeOH/$H_2O$ containing 0.05% TFA at the flow rate of 4 mL $min^{-1}$. Medlan A1 (3.2 mg) and A2 appeared at 20.5-24.0 min and 14.8-15.6 min respectively.

Larlan A2 and A5 (**14**, **15**)

The larlans were purified from the crude extract of heterologously expressed genes *larAMTBCDEFHGIJ* in recombinant *H. volcanii* H1424 cultivated in Hv-YPC medium for 3 days. Larlan A2 was eluted with 15% ACN/$H_2O$ containing 0.05% TFA on the HPLC system with a Phenomenex Biphenyl column (250 mm × 10 mm, 5 μm, 100 Å) at the flow rate of 4 mL $min^{-1}$. Peak larlan A2 (**14**) appeared at 31.8-35.2 min. Larlan A5 (**15**) was eluted with 39% MeOH containing 0.05% TFA on an HPLC system with a Phenomenex Kinetex XB-C18 column (250 mm × 10 mm, 5 μm, 100 Å) at the flow rate of 3 mL $min^{-1}$. Peak larlan A5 (**15**) appeared at 56.0-62.0 min.

Archalan β3 (**20**)

The fermentation and production of archalan β3 (**20**) were performed as previously reported [1]. The fermentation broth of the wild-type strain *Haloferax salinus* was collected after cultivation in modified DSMZ medium 1520 for 5 days at 37 °C. The crude extract passed through an open ODS column, eluting with a gradient concentration of MeOH-$H_2O$ (10-50% at intervals of 10%, and finally 100%). The target compound was detected in the fraction of 40% MeOH through LC-MS analysis. Following this, the fraction was evaporated and loaded to a Sephadex LH-20 column (Solarbio, China) using a solvent mixture of 60% MeOH/$H_2O$ system to separate and concentrate archalan β3 (**20**) and its analogs. Those fractions with target compound were merged, evaporated, and subjected to an HPLC system on Phenomenex Kinetex XB-C18 column (250 mm × 10 mm, 5 μm, 100 Å) with 32% MeOH/$H_2O$ containing constant 0.05% TFA at the flow rate of 4 mL $min^{-1}$. Archalan β3 (1.0 mg) appeared at 40.0-44.0 min.

Archalan γ (**6**)

The above obtained crude extract of *H. salinus* was analyzed by an HPLC system (Waters, Parsippany, NJ, United States) with a Phenomenex Luna C18 column (250 mm × 10 mm, 5 μm, 100 Å). A gradient method was used ranging from 5%-95% ACN/$H_2O$ containing constant 0.1% TFA within 5-70 min at the flow rate of 3 mL $min^{-1}$. Archalan γ (**6**) appeared at 39.0-43.0 min.

Two different single mutants of LarA

Each MeLan residues in the larlan A2 (**14**) have four different possibilities, LL-Melan, DL-Melan, L-*allo*-L-MeLan, and D-*allo*-L-MeLan. Therefore, two single mutants of Thr to Ser (T34S and T46S) were constructed and transferred in *H. volcanii* H1424. Around 200 mL culture broth of each mutant strain was extracted by HP20 for HPLC separation. They were purified in different conditions. LarA2 T2S was purified in a Phenomenex Kinetex XB-C18 column (250 mm × 4.6 mm, 5 μm, 100 Å) with a gradient of 15%-20% ACN/$H_2O$ containing 0.05% TFA from 5.0-45.0 min at 1 mL $min^{-1}$ and appeared at 29.2-30.2 min. LarA2 T9S was purified in a Phenomenex Kinetex XB-C18 column (250 mm × 4.6 mm, 5 μm, 100 Å) with a gradient of 20%-60% ACN/$H_2O$ containing 0.05% TFA from 5.0-35.0 min at 1 mL $min^{-1}$ and appeared at 9.6-10.1 min.

#### Advanced Marfey's analysis for purified archaeal lanthipeptides

Amino acid configurations of purified archaeal lanthipeptides were determined using the advanced Marfey's method[2,3]. Briefly, lanthipeptides (~ 0.2 mg) were dissolved in 1 mL of 6 N HCl (with 5% (v/v)) heated at 100 °C for 16 h. The hydrolysates were evaporated, redissolved in 100 µL of water, and aliquoted into two portions. Each portion and standard sample were treated with 20 µL of $NaHCO_3$ (1 M) and 50 µL of L-FDLA or D-FDLA (1 M) at 40 °C for 2 h, then quenched with 20 µL HCl (1 M) and dried under compressed air. The mixtures were dissolved in 400 µL of MeOH for LR LC-MS or HR LC-MS analysis. Separations were carried out on a Waters Acquity UPLC BEH C18 column (2.1 mm × 150 mm ID, 1.7 µm) by using a gradient elution mode at a flow rate of 0.2 mL $min^{-1}$: 5%-80% ACN/$H_2O$ with constant 0.1% formic acid in 2.0-20.0 min, 80%-100% in 20.0-25.0 min, then isocratic 100% for 4 min. The stereochemistry was determined by comparing the retention time of L/D-FDLA derivatized amino acids. For the configuration determination of residue MeLan, hydrolysates were obtained and analyzed by HR LC-MS using the gradient from 40%-50% ACN/$H_2O$ with 0.1% formic acid in 2.0-25.0 min with a Waters ACQUITY Premier Peptide BEH C18 column (1.7 µm, 300 Å, 2.1 × 150 mm). The synthetic DL-Lan, LL-Lan, LL-Melan, L-*allo*-L-MeLan, and D-*allo*-L-MeLan from Dong's lab[4] and nisin were used as a reference to confirm the stereochemistry of the MeLan and Lan residues of these lanthipeptides by HR LCMS.

#### Reductive desulfurization, DTT reduction and LC-MS characterization of archalan γ (6)

The HPLC fraction containing archalan γ (**6**) (39-43 min, ~ 3.0 mg) was suspended in 4.0 mL of $CH_3OH/H_2O$ (1:1) or $CD_3OD/D_2O$ (1:1), to which 20 mg of $NiCl_2$ and 20 mg of $NaBH_4/NaBD_4$ were added. This mixture was stirred under 1 atm of $H_2$ at room temperature for 2 h for partial desulfurization and 8 h for total desulfurization. Then, the mixture was centrifuged, and the supernatant was collected. A mixed solvent of $CH_3OH/H_2O$ (ration=1:1, 1.0 mL) was added to the black nickel boride pellets, and the suspension was sonicated and re-centrifuged to recover any residual peptide. Combined supernatants were dried under vacuum and stored at -20 ºC before HR-LCMS and $MS^n$ analyses. Dithiothreitol (DTT) was dissolved in the buffer (50 mM Tris-HCl, 500 mM NaCl, 10% glycoses, pH = 8.0). A crude extract containing **6** was treated with 0.1 M DTT solution for 2 h at 55 °C. Then, the reduced sample was cooled down and purified by Zip Tip C18 tips (Millipore, Billerica, MA, USA) to do LC-MS analysis and stored at -20 °C. The partial reduction supernatants were analyzed on LTQ Orbitrap Velos mass spectrometer (Thermo Scientific, San Jose, CA, USA).

#### Bioassay test for archaeal lanthipeptides

Except for purified archaeal lanthipeptides, crude extracts containing the classical lanthipeptides identified through LC-MS/MS analysis were also subjected to a bioassay test using the extracts from the host strain *H. volcanii* H1424 (with empty plasmid pTA1228) for comparison. The crude extracts of recombinant strains with different lanthipeptide BGCs or the empty plasmid were initially cultivated in 200 mL YPC medium for 5 days. Subsequently, the extracts were subjected to extraction using HP20 resin with elution using 0%, 50%, and 100% MeOH. All the crude extracts were processed through a C18 solid-phase extraction (SPE) cartridge to accumulate the target lanthipeptides. The compounds in 100% MeOH fraction were further going through a 1 mL C18 SPE column eluted by 0, 5, 10, 20, 30, 50, 70, and 100% MeOH. The target lanthipeptides in 70% and 100% MeOH fractions were finally redissolved in dd$H_2O$, and subsequently subjected to centrifugation to prevent halocins.

Ten haloarchaea strains, including *H. larsenii* JCM 13917, *H. mediterranei* ATCC 33500, *Natrinema saccharevitans* JCM 12889, *N. gari* JCM 14663, *Haladaptatus cibarius* DSM 19505, *H. paucihalophilus* JCM 13897, *H. amylolyticus* JCM 18367, *H. salinus* YJ-37-H, *Halomicroarcula salina* JCM 18369, two halophilic bacterial strains, *Halobacillus profundi* JCM 14154 and *H. kuroshimensis* JCM 14155, and other four bacteria, Methicillin-resistance *Staphylococcus aureus* ATCC 43300, *Acinetobacter baumannii* ATCC 19606, *Bacillus subtilis* 168, *Escherichia coli*

ATCC 25922 were used as indicator strains for antimicrobial test. Haloarchaea were cultured in modified DSMZ medium 589. Their cell density was adjusted to $5.0 \times 10^5$ c.f.u. $mL^{-1}$ in the 96-well microtiter plates (Corning, USA). These cultures were then mixed with varying compounds and incubated in the shaking incubator (37 °C and 200 r.p.m.) for 2-7 days. According to the Japan Collection of Microorganisms, two halophilic bacterial strains were cultivated in 464 mediums. Other bacterial strains were grown in LB medium. After overnight (37 °C and 200 r.p.m.) cultivation, the cultures were adjusted to $5.0 \times 10^5$ c.f.u. $mL^{-1}$ in the 96-well microtiter plates, mixed with different compounds, and then incubated in the static incubator (37 °C and 200 r.p.m.) for 24 h. The volume was 100 μL in each well. The crude extract samples were prepared in a stock concentration of 10 μg $\mu L^{-1}$. The same volume of stock solutions, DMSO or $ddH_2O$, were added and tested as the negative control. The cell growth was monitored at $OD_{600}$ (Varioskan Flash, Thermo Scientific, USA). The vitality was calculated by subtracting the initial $OD_{600}$ value (at 0 h) from the final $OD_{600}$ value (at 24 h) for each group and then dividing this difference by the $OD_{600}$ value of the control group. The inhibition value was determined by subtracting the calculated viability value from 100%. Figures were performed by GraphPad Prism 8. The minimal inhibitory concentration (MIC) of the pure compound against each strain was defined as the lowest compound concentration at which archaeal growth was not observed.

#### Growth curve analysis

To investigate the potential correlation between growth regulation and motility regulation induced by lanthipeptide BGCs, the growth curves of recombinant strains with the empty vector pTA1228, pTA1228-BGC *cib2*, pTA1228-BGC *pel,* and pTA1228-BGC *alna* were tested. Three independent mid-log phase cell cultures of each strain were diluted to $OD_{600}$ = 0.1 (50 μL in Corning 96-well plates, Thermo Scientific Varioskan Flash), and 200 μL of each dilution was added to 15 mL cell culture tube with 3 mL liquid Hv. Ca medium for cultivation at 45 °C with continuous shaking at 200 r.p.m. The $OD_{600}$ readings for each strain were measured at various time points by taking 200 μL of culture medium in 96-well plates (Corning), and the data were visualized using GraphPad Prism 8 software.

#### Supplementary Figures

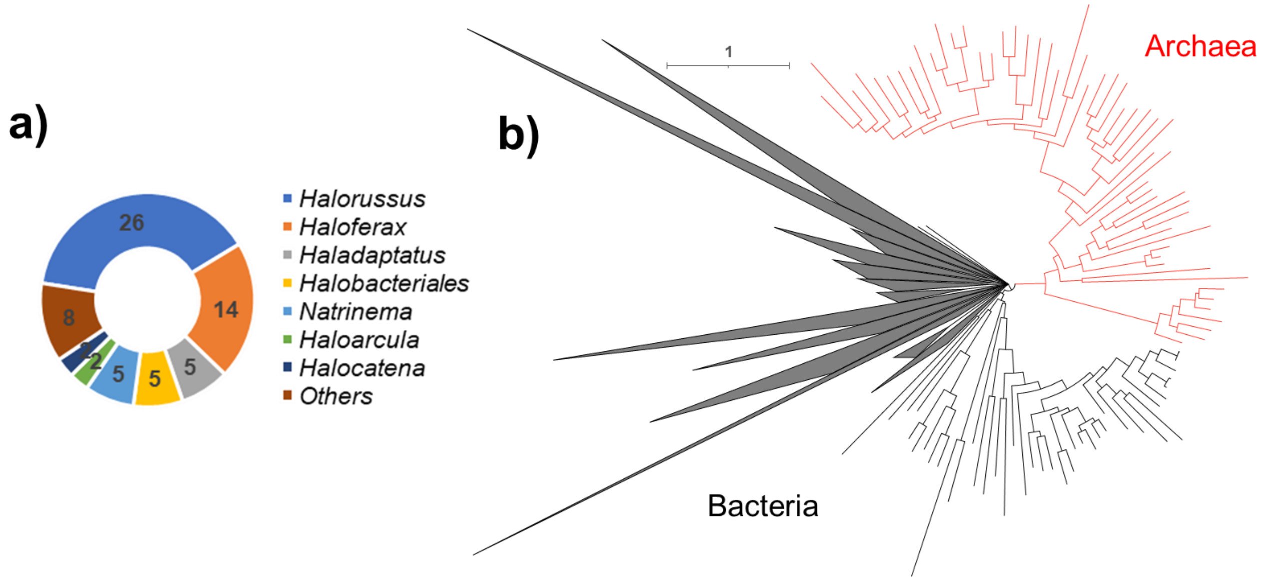

**Supplementary Figure 1. Genomic analysis of archaeal LanMs**

a, The distribution of 67 deduplicated archaeal LanMs at the genus level. b, The Maximum Likelihood (ML) tree of 110 archaeal and 12,398 bacterial LanMs (Supplementary Data 2). The sequences were deduplicated based on an 80% similarity threshold, and the tree was rooted at the midpoint for visualization. The distinct clustering of archaeal LanMs separate from bacterial LanMs suggests significant evolutionary divergence between these two groups of lanthipeptide synthetases.

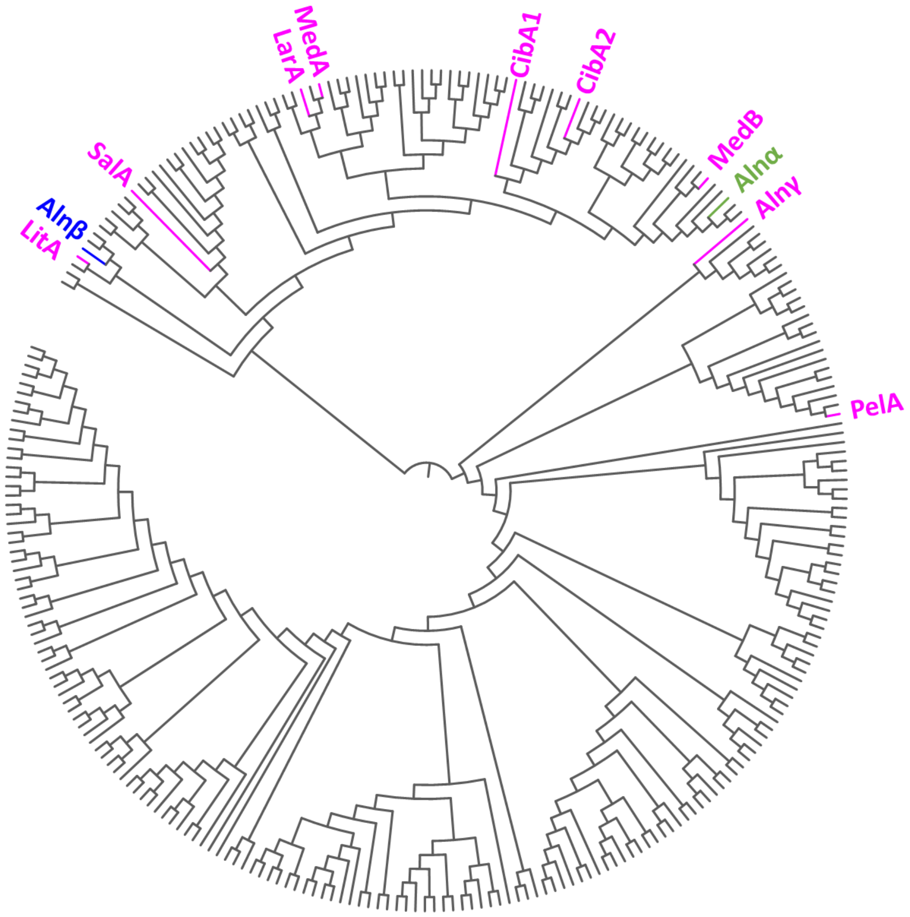

**Supplementary Figure 2. Maximum Likelihood (ML) tree of 353 archaeal LanAs**

Our previously reported lanthionines, archalans α and β, were derived from Alnα and Alnβ, appearing in green and blue, respectively. This study introduces newly characterized LanAs named Alnγ, LitA, SalA, LarA, MedA, CibA1, CibA2, MedB, and PelA, highlighted in rose red. The phylogenetic tree illustrates that LanAs within the same sequence similarity network (SSN) tend to cluster together, such as LitA with Alnβ, and CibA1 with CibA2. Additionally, the ML tree displays two main clades, one comprising most of the characterized LanAs and the other including Alnγ and PelA. Within the Alnγ and PelA clade, numerous unexplored archaeal LanAs are identified, with many of them belonging to the first cluster in Figure 1. This observation underscores the novelty of archaeal LanAs.

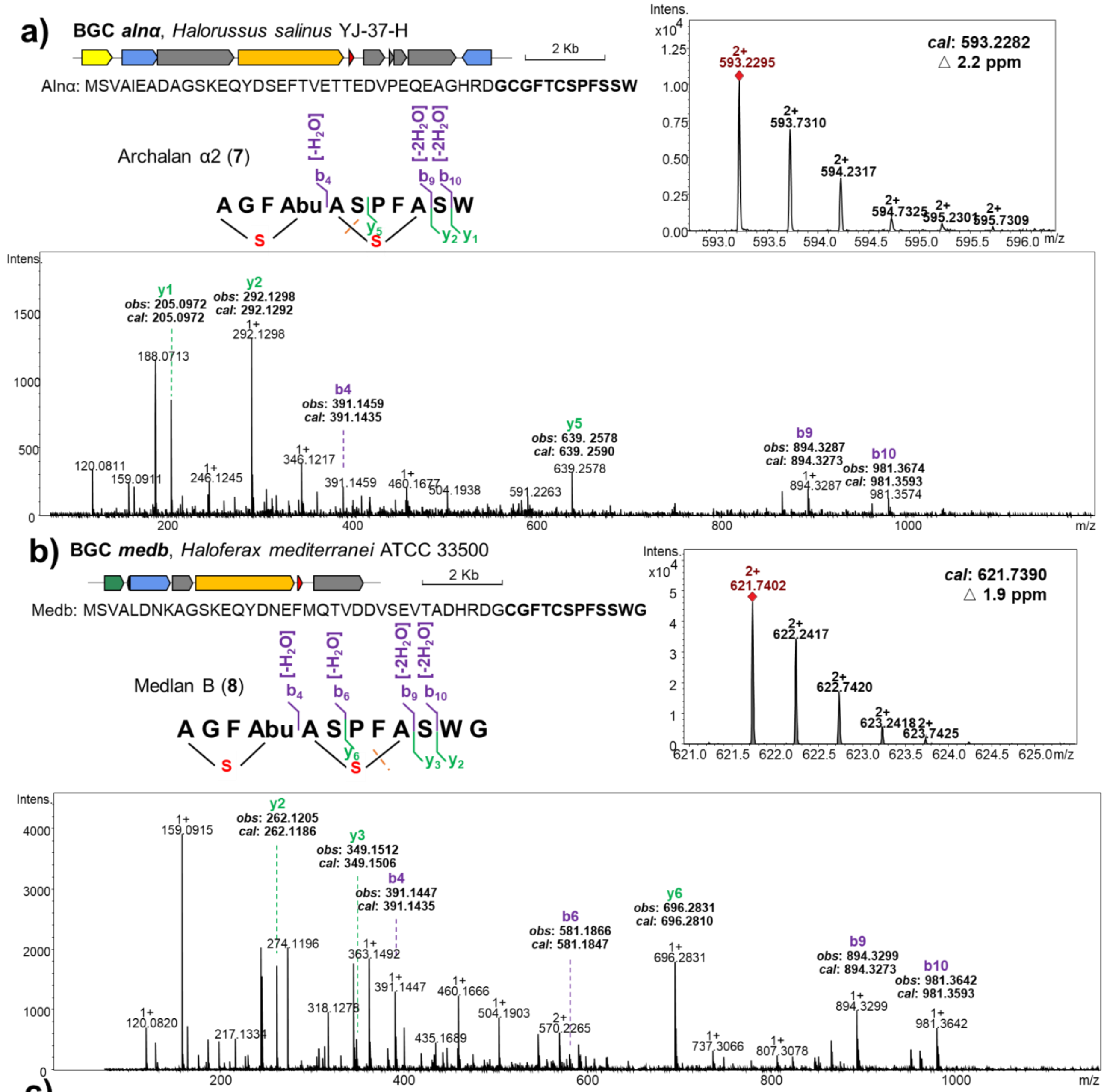

c)

| | Archalan α2 (7) | | | Medlan B (8) | | | | Archalan α2 (7) | | | Medlan B (8) | | |
|---|---|---|---|---|---|---|---|---|---|---|---|---|---|
| Ion | Obs m/z | Calc m/z | △ppm | Obs m/z | Calc m/z | △ppm | Ion | Obs m/z | Calc m/z | △ppm | Obs m/z | Calc m/z | △ppm |
| b4 | 391.1459 | 391.1435 | 6.1 | 391.1447 | 391.1435 | 3.1 | | | | | | | |
| b6 | | | | 581.1866 | 581.1847 | 3.3 | y6 | | | | 696.2831 | 696.2810 | 3.0 |
| | | | | | | | y5 | 639.2578 | 639.2590 | 1.9 | | | |
| b9 | 894.3287 | 894.3273 | 1.6 | 894.3299 | 894.3273 | 2.9 | y3 | | | | 349.1512 | 349.1506 | 1.7 |
| b10 | 981.3674 | 981.3593 | 8.3 | 981.3642 | 981.3593 | 5.0 | y2 | 292.1298 | 292.1292 | 2.1 | 262.1205 | 262.1186 | 7.2 |
| | | | | | | | y1 | 205.0972 | 205.0972 | 0.0 | | | |

**Supplementary Figure 3. LC-MS/MS analysis of heterologously expressed classical lanthipeptides archalan α2 (7) and medlan B (8)**

BGCs *alnα* (a) and *medb* (b) were identified from *H. salinus* YJ-37-H and *H. mediterranei* ATCC35500, respectively. The peptide signals, $[M+2H]^{2+}$ = 593.2295 (calculated $[M+2H]^{2+}$ = 593.2282, Δ = 2.2 ppm) and $[M+2H]^{2+}$ = 621.7402 (calculated $[M+2H]^{2+}$ = 621.7390, Δ = 1.9 ppm), were putative lanthipeptide products (a-c). LC-MS/MS analysis of these two lanthipeptides unveiled key ion masses of $b_4$ (-$H_2O$, thioether ring in Cys1-Thr4), $b_9$ (-2$H_2O$, thioether ring in Cys5-Ser9), and $y_5$ or $y_6$ (thioether ring in Cys5-Ser9) in archalan α2 (**7**) and medlan B (**8**). This analysis revealed the presence of C-S crosslinks in Cys1-Thr4 and Cys5-Ser9, consistent with the NMR-verified structure of archalan α (**25**). Both archalan α2 (a) and medlan B (b) were detected in the heterologous expression of BGC *medb*, possibly due to the cryptic peptidase from the host genome. This observation was further supported by the identification of archalan α2 (a) rather than the natural product archalan α in the heterologous expression result of BGC *alnα*. Gene annotations are colored as follows: LanM in orange, LanA in red, transporter-related genes in blue, regulator genes in green, putative peptidase genes in yellow, methyltransferase genes in purple, and other genes in grey. This legend applies to the subsequent LC-MS/MS analysis results, except for special statements.

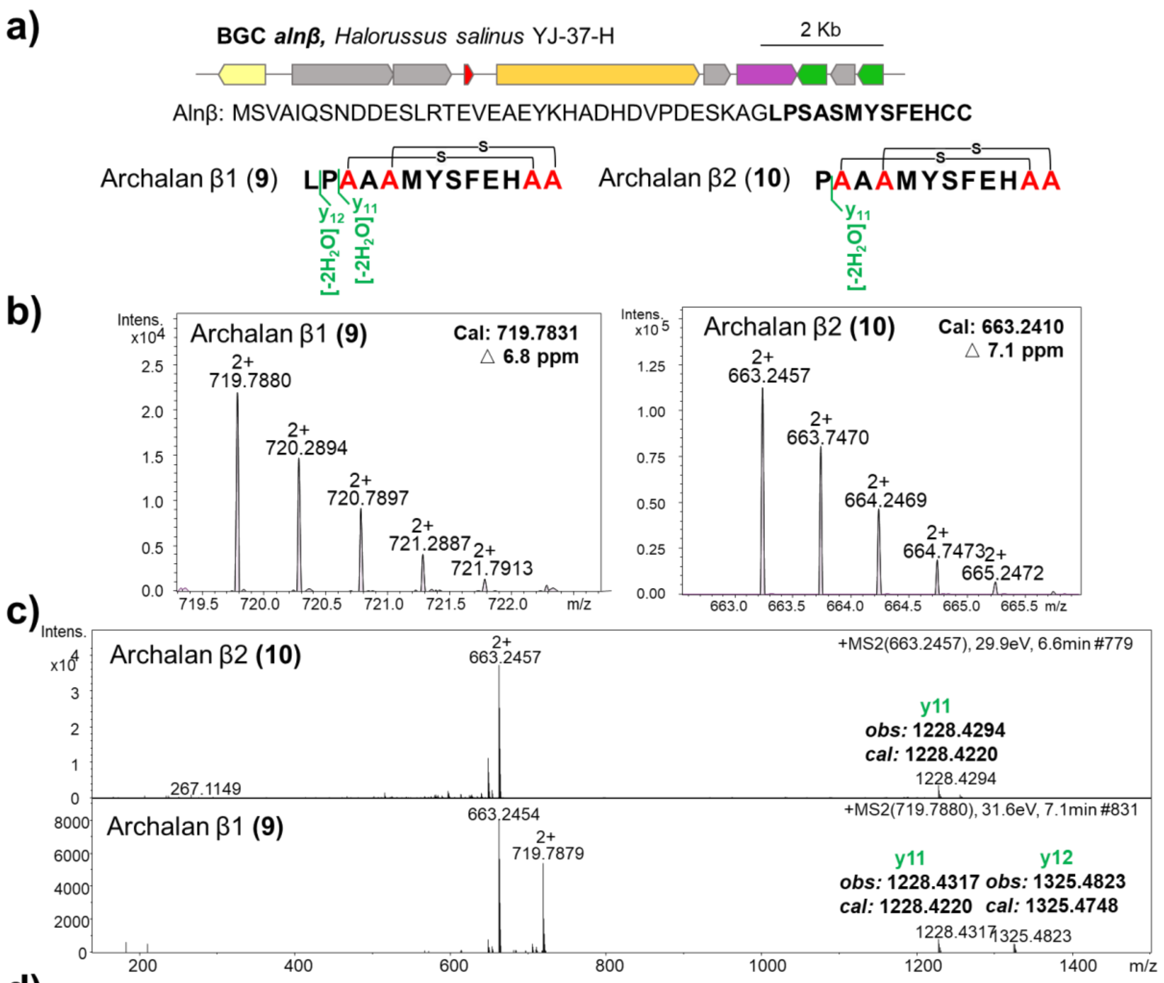

d)

| | Archalan β1 (**9**) | | | Archalan β2 (**10**) | | |
|---|---|---|---|---|---|---|
| Ion | Obs m/z | Calc m/z | △ppm | Obs m/z | Calc m/z | △ppm |
| y11 | 1228.4317 | 1228.4220 | 7.9 | 1228.4294 | 1228.4220 | 6.0 |
| y12 | 1325.4823 | 1325.4748 | 5.7 | | | |

**Supplementary Figure 4. LC-MS/MS analysis of heterologously expressed classical lanthipeptide archalan β1-2 (9-10)**

The BGC *alnβ* was identified from *H. salinus* YJ-37-H (a). b-d, Peptide signals indicative of lanthipeptide products were observed as $[M+2H]^{2+}$ = 719.7880 (calculated $[M+2H]^{2+}$ = 719.7831, Δ = 6.8 ppm) and $[M+2H]^{2+}$ = 663.2457 (calculated $[M+2H]^{2+}$ = 663.2410, Δ = 7.1 ppm). A modification of $-2H_2O$ on their corresponding core peptides of Alnβ (LPSASMYSFEHCC or PSASMYSFEHCC) was proposed based on MS1 analysis (a,b), with MS/MS fragments encompassing only amino acids outside the large cyclic rings (c,d), akin to the structure of archalan β as previously documented in our research[1]. Archalans β, β1 (**9**), and β2 (**10**) originate from the same precursor, differing in the N-terminal amino acid overhangs, as confirmed by the different y ions (c,d). The crosslinks were inferred according to the structure of archalan β elucidated through LC-MS/MS analysis in the previous study and the resolved archalan β3 (**20**) structure via NMR in this investigation (Supplementary Fig. 31). All variants displayed a Ser-Cys crosslink with an intertwined configuration (a). Notably, the recombinant strain produced classical lanthipeptides archalan β1-2 (**9-10**) while failing to yield the DADC lanthipeptide archalans β3-7 (**20-24**) as the wide-type strain despite the inclusion of a putative peptidase within the BGC. This suggests the mysterious proteases outside the BGC may be involved in the biosynthesis of DADC lanthipeptides.

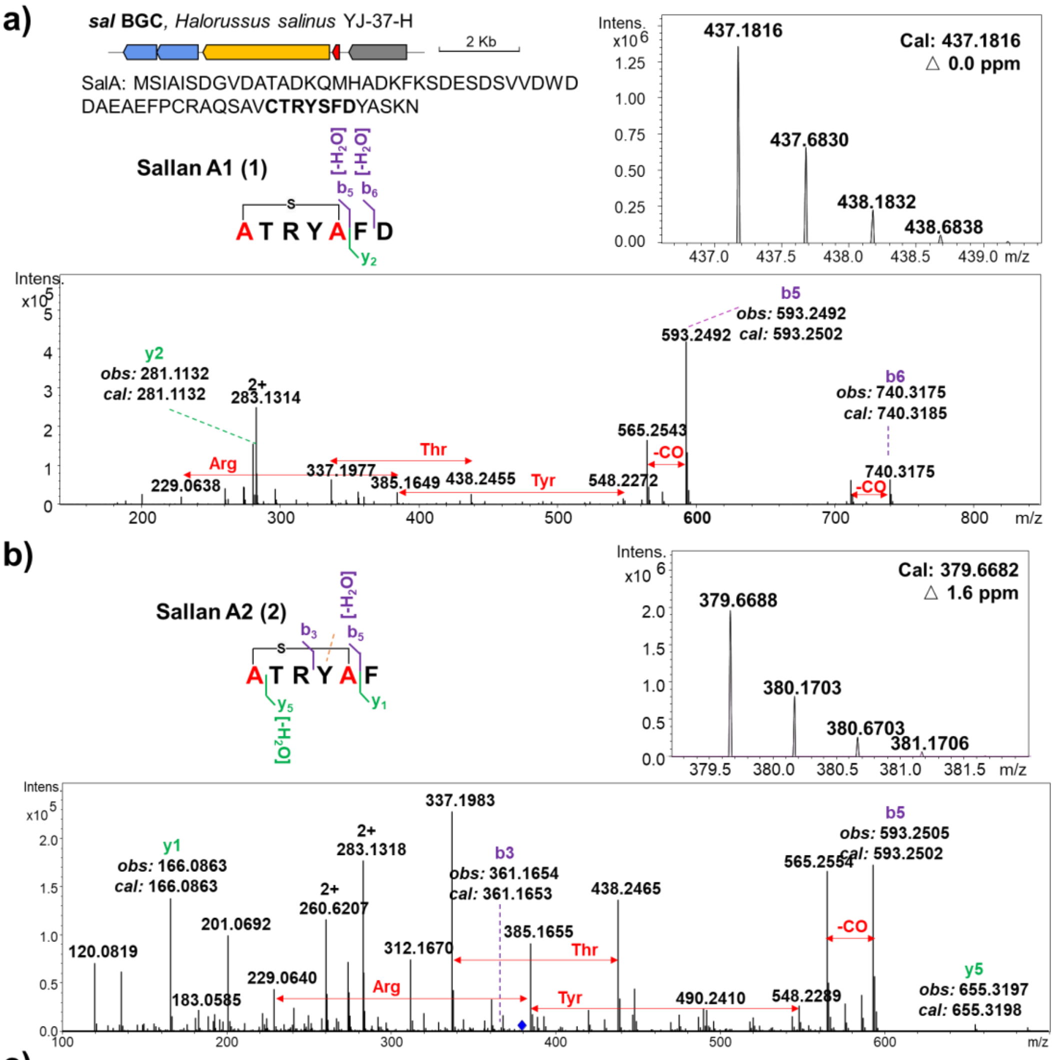

c)

| | Sallan A1 (1) | | | Sallan A2 (2) | | | | Sallan A1 (1) | | | Sallan A2 (2) | | |
|---|---|---|---|---|---|---|---|---|---|---|---|---|---|
| Ion | Obs m/z | Calc m/z | △ppm | Obs m/z | Calc m/z | △ppm | Ion | Obs m/z | Calc m/z | △ppm | Obs m/z | Calc m/z | △ppm |
| | | | | | | | y5 | | | | 655.3197 | 655.3198 | 0.2 |
| b3 | | | | 361.1654 | 361.1653 | 0.3 | | | | | | | |
| b5 | 593.2492 | 593.2502 | 1.7 | 593.2505 | 593.2502 | 0.5 | | | | | | | |
| b6 | 740.3175 | 740.3185 | 1.4 | | | | y2 | 281.1132 | 281.1132 | 0.0 | | | |
| | | | | | | | y1 | | | | 166.0863 | 166.0863 | 0.0 |

**Supplementary Figure 5. LC-MS/MS analysis of heterologously expressed classical lanthipeptide sallan A1-2 (1-2)**

BGC *sal*, identified from *Halorussus salinus* YJ-37-H, was heterologously expressed in *Haloferax volcanii* H1424. Mass spectrometry analysis revealed putative lanthipeptide products, $[M+2H]^{2+}$ = 437.1816 (calculated $[M+2H]^{2+}$ = 437.1816, Δ = 0.0 ppm) and $[M+2H]^{2+}$ = 379.6688 (calculated $[M+2H]^{2+}$ = 379.6682, Δ = 1.6 ppm), with one dehydration in their corresponding core peptides (a-c). Sallan A1 (**1**) and A2 (**2**) are derived from the same precursor SalA, differing by only one amino acid at the C-terminus which could be caused by a peptidase from the host genome. The amino acid sequences of sallans in the figure are annotated with b and y ions, indicating the putative thioether ring locations. NMR analysis (Supplementary Fig. 6) confirmed the thioether ring between Ser1 and Cys5 in sallan A2 (**2**), providing further support for the crosslink observed in sallan A1 (**1**) and A2 (**2**) during MS/MS analysis. Notably, these products were exclusively detected in the crude extract of the recombinant strain, emphasizing the benefits of heterologous expression in Haloarchaea.

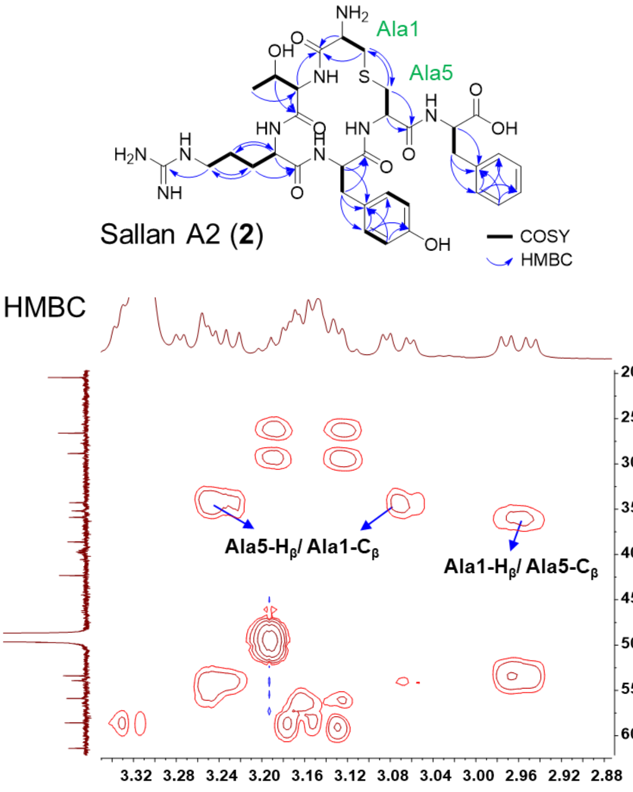

#### Supplementary Figure 6. Structure elucidation of purified archaeal lanthipeptide sallan A2 (2)

The mass signal of sallan A2 (**2**), observed as $[M+2H]^{2+}$ = 379.6688, was predicted with a molecular formula of $C_{34}H_{47}N_9O_9S$ (*cal*: $[M+2H]^{2+}$ = 379.6682, Δ = 1.6 ppm), matching with the predicted core peptide (CTRYSF) of BGC *sal* with one dehydration modification. To fully characterize its structure, 1.1 mg of **2** was purified from 5 L culture broth of the recombinant strain, and its structure was elucidated using extensive NMR analysis in MeOH-$d_4$. The observation of many carbonyl carbons ($\delta_C$ 168.0-174.0 ppm) in $^{13}$C-NMR spectra supported the peptidic nature of **2** (Supplementary Fig. 40 and Supplementary Table 3). Apart from the free carboxyl groups, the detection of five carbonyl carbons was consistent with the predicted presence of six amino acids in the core peptide (Supplementary Fig. 40 and Supplementary Table 3). Through HSQC, COSY, and HMBC data, compound **2** was further deduced to contain 1 × Thr, 1 × Arg, 1 × Tyr, 1 × Phe, and 1 × Lan. The HMBC correlation of Ala1-$H_β$ to Ala5-$C_β$ ($\delta_H$ = 2.96 ppm to $\delta_C$ = 35.9 ppm) and Ala5-$H_β$ to Ala1-$C_β$ ($\delta_H$ = 3.07, 3.24 ppm to $\delta_C$ = 34.3 ppm) suggested a C-S crosslink between Ala1 and Cys5 (Supplementary Fig. 40 and Supplementary Table 3). The stereochemistry of **2** was confirmed with L configuration in all unmodified amino acids and DL-Lan (2*S*,6*R*) residue in the thioether ring by advanced Marfey's analysis (Supplementary Fig. 9 and Supplementary Table 4). Notably, **2** with only six amino acids represents the smallest natural lanthipeptide to date.

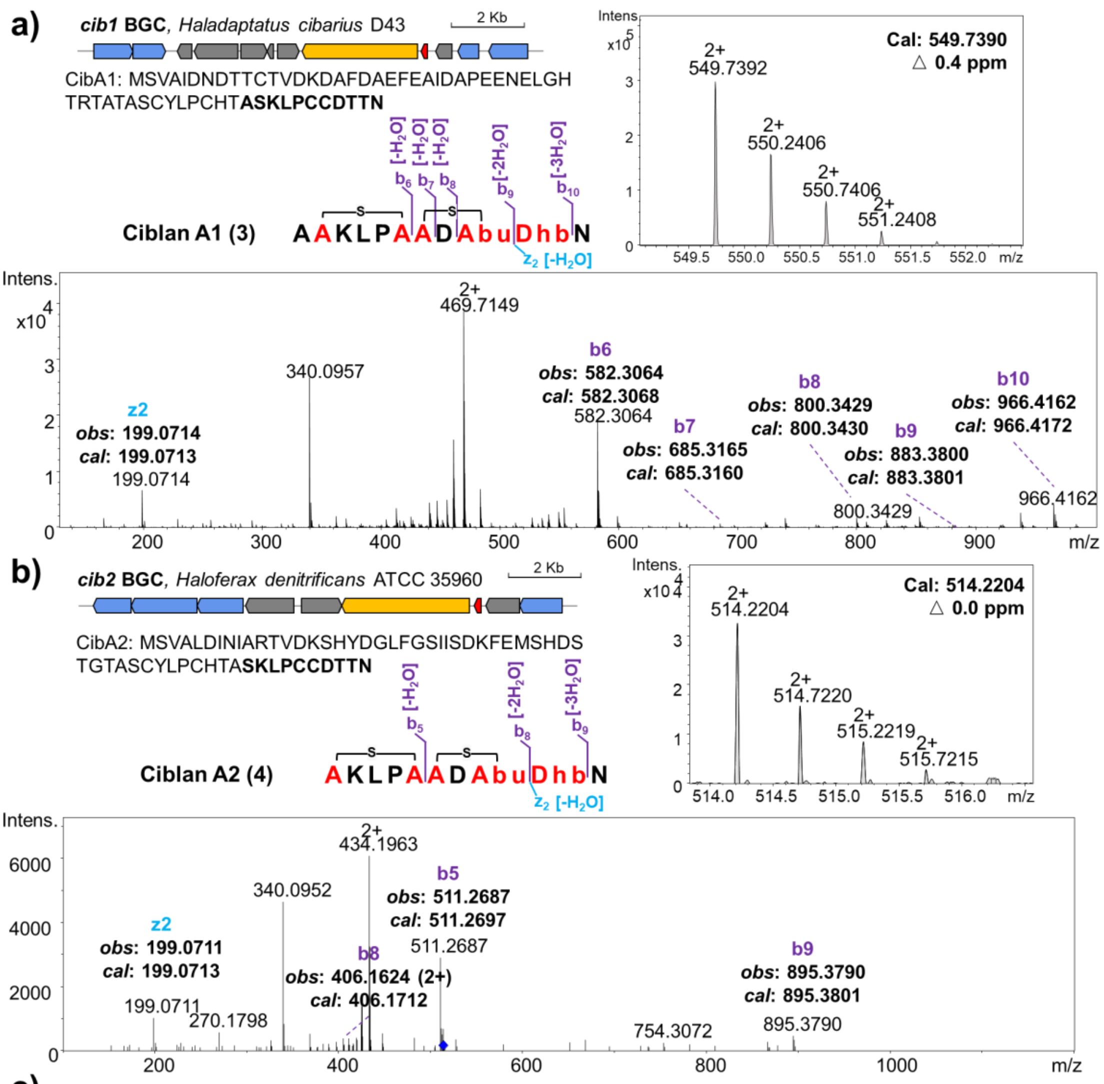

c)

| | Ciblan A1 (**3**) | | | Ciblan A2 (**4**) | | |
|---|---|---|---|---|---|---|
| Ion | Obs m/z | Calc m/z | △ppm | Obs m/z | Calc m/z | △ppm |
| b5 | | | | 511.2687 | 511.2697 | 2.0 |
| b6 | 582.3064 | 582.3068 | 0.7 | | | |
| b7 | 685.3165 | 685.3160 | 0.7 | | | |
| b8 | 800.3429 | 800.3430 | 0.1 | 406.1624 | 406.1712 | 21.7 |
| b9 | 883.3800 | 883.3801 | 0.1 | 895.3790 | 895.3801 | 1.2 |
| b10 | 966.4162 | 966.4172 | 1.0 | | | |
| z2 | 199.0714 | 199.0713 | 0.5 | 199.0711 | 199.0713 | 1.0 |

**Supplementary Figure 7. LC-MS/MS analysis of heterologously expressed classical lanthipeptide ciblan A1-2 (3-4)**

The BGCs *cib1* (a) and *cib2* (b) were identified from *Haloferax denitrificans* ATCC 35960 and *Haladaptatus cibarius* D43, respectively. Mass spectrometry analysis revealed putative lanthipeptide products with peptide signals $[M+2H]^{2+}$ = 549.7392 (calculated $[M+2H]^{2+}$ = 549.7390, Δ = 0.4 ppm) and $[M+2H]^{2+}$ = 514.2204 (calculated $[M+2H]^{2+}$ = 514.2204, Δ = 0.0 ppm), indicating a connection to BGCs *cib1* and *cib2* respectively (a-c). SSN analysis indicated that these two precursors, originating from the same cluster, might exhibit a similar structure, a finding corroborated by our results. Importantly, NMR analysis of ciblan A1 (**3**) confirmed the presence of two thioether rings between Ser2-Cys6, Cys7-Thr9, and one Dhb in Thr10 (Supplementary Fig. 8). The key MS/MS fragments, including $b_5$ ($-H_2O$, thioether ring in Ser1-Cys5), $b_8$ ($-2H_2O$, another ring in Cys6-Thr8), $b_9$ ($-3H_2O$, dehydration of Thr9) and $z_2$ ($-H_2O$, dehydration of Thr9) observed in ciblan A2 (**4**), supported identical crosslinks and dehydrations as seen in ciblan A1 (**3**).

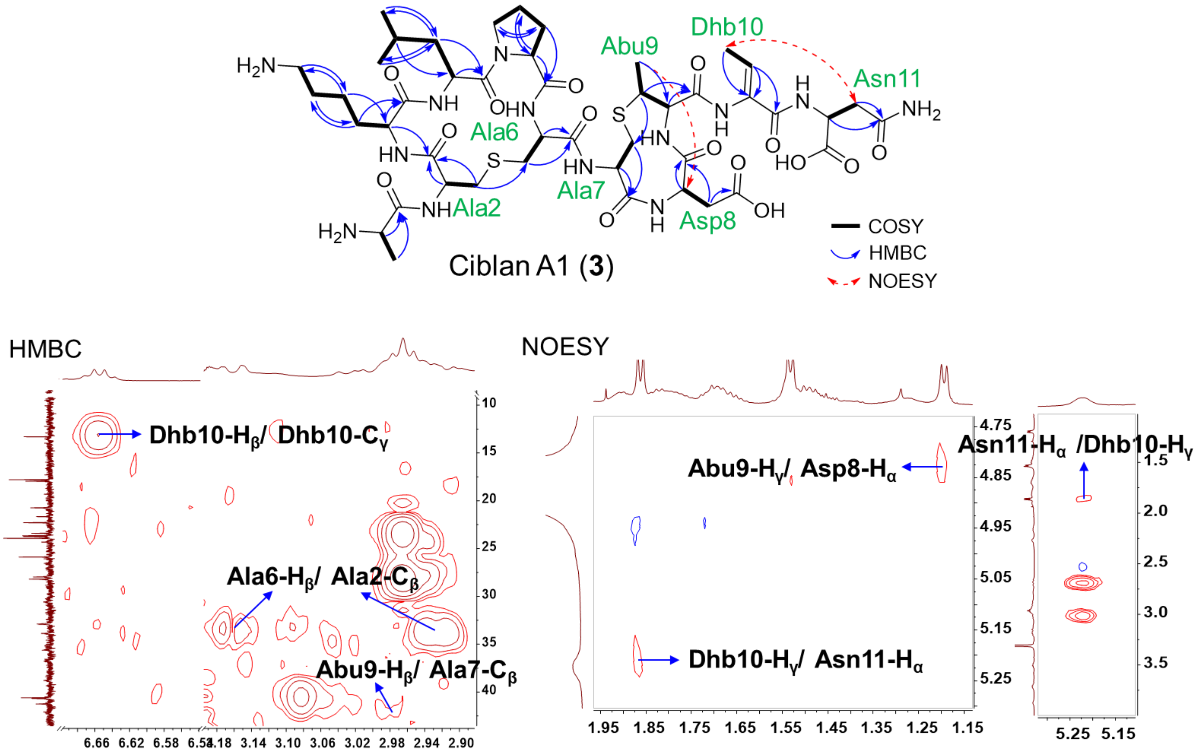

#### Supplementary Figure 8. Structure elucidation of purified archaeal lanthipeptide ciblan A1 (3)

The mass signal of ciblan A1 (**3**), observed as $[M+2H]^{2+}$ = 549.7392, was predicted with a molecular formula of $C_{45}H_{71}N_{13}O_{15}S_2$ (*cal*: $[M+2H]^{2+}$ = 549.7390, Δ = 0.4 ppm), matching a loss of three dehydrations on the predicted core peptide of BGC *cib1* (ASKLPCCDTTN). Ultimately, 1.0 mg of **3** was purified from 8 L culture broth of the wild-type strain and its structure was elucidated using extensive NMR analysis in MeOH-$d_4$. The observation of many carbonyl carbons ($\delta_C$ 170.0-175.0 ppm) in $^{13}$C-NMR spectra supported the peptidic nature of **3** (Supplementary Fig. 41 and Supplementary Table 3). Through HSQC, COSY, and HMBC data, compound **3** was further deduced to contain 1 × Ala, 1 × Lys, 1 × Leu, 1 × Pro, 1 × Asp, 1 × Asn, 1 × Dhb, 1 × Lan and 1 ×MeLan. The NMR data of purified **3** exhibited the presence of Dhb residue ($C_\alpha$= 131.0 ppm, $C_\beta$ = 130.9 ppm, $H_\beta$ = 6.66 ppm, $C_\gamma$ = 13.4 ppm, $H_\gamma$ = 1.86 ppm), which had NOESY correlation with Asn11 ($\delta_H$ = 1.20 ppm from Dhb10 and $\delta_H$ = 5.22 ppm from Asn11), thereby confirming its position as derived from Thr10. Moreover, the key HMBC correlations exhibited two thioether rings in Ser2-Cys6 ($\delta_H$ = 2.97, 2.92 ppm of Ala6 to $\delta_C$ = 35.0 ppm of Ala2) and Cys7-Thr9 ($\delta_H$ = 2.98 ppm of Abu9 to $\delta_C$ = 41.2 ppm of Ala7). Additionally, the *Z*-geometry of the double bond in Dhb residue was determined based on the corresponding $^3J$ coupling value of 7.1 Hz (Supplementary Fig. 41 and Supplementary Table 3). The stereochemistry of **3** was confirmed with L configuration in all unmodified amino acids, along with DL-Lan (2*S*,6*R*) and DL-MeLan (2*S*,3*S*,6*R*) configurations in thioether rings by comparing the standard samples in advanced Marfey's analysis (Supplementary Fig. 9 and Supplementary Table 4).

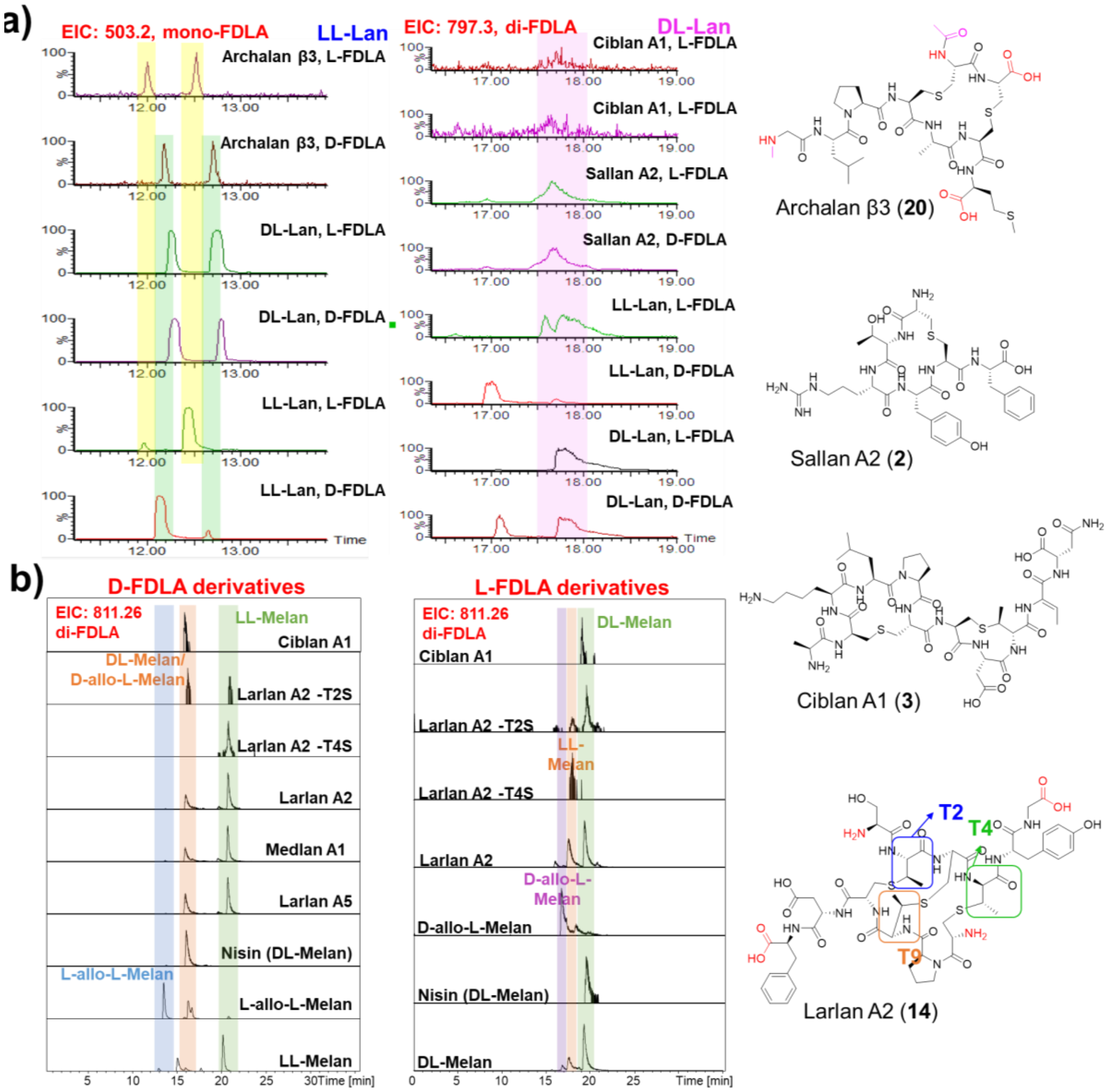

**Supplementary Figure 9. Advanced Marfey's analysis of archaeal lanthipeptides**

The advanced Marfey's analysis revealed that all the unmodified amino acids in isolated lanthipeptides are L configurations (Supplementary Table 4). However, the lanthionine and methyllanthionine rings are different in specific lanthipeptides. The two thioether rings are all LL (2*R*,6*R*) configurations in **20,** while DL (2*S*,6*R*) configurations in **2** and **3** (a). In addition, **3** harbored another DL-MeLan (2*S*,3*S*,6*R*) residue compared with the standard samples (b). As for three MeLan residues in larlans and medlan A1 increasing the complexity of stereochemistry configuration, two single mutations of Thr to Ser were isolated for comparative stereochemistry analysis (larlan A2-T2S and -T4S). The D-FDLA derivatives of larlan A2-T4S displayed a single peak that matched the standard sample LL-MeLan, representing that MeLan rings in Thr2 and Cys10, Thr9 and Cys3 are LL configurations. In comparison, larlan A2-T2S mutant displayed two peaks, with one peak confirmed to contain LL-MeLan residue, while another peak contained either DL-MeLan or D-allo-L-MeLan residue. The D-FDLA derivatives of DL-MeLan or D-allo-L-MeLan cannot be distinguished or separated as reported[4, 5]. Hence, L-FDLA derivatives confirmed another peak to be DL-MeLan, suggesting another thioether ring between Thr4 and Cys7 in a DL-MeLan configuration. Altogether, the first two thioether rings (Thr2-Cys10, Thr9-Cys3) are LL-MeLan (2*R*,3*R*,6*R*), the last is DL-MeLan (2*S*,3*S*,6*R*), and all other unmodified amino acids are proteinogenic L configurations in larlan A2 (**14**).

a)

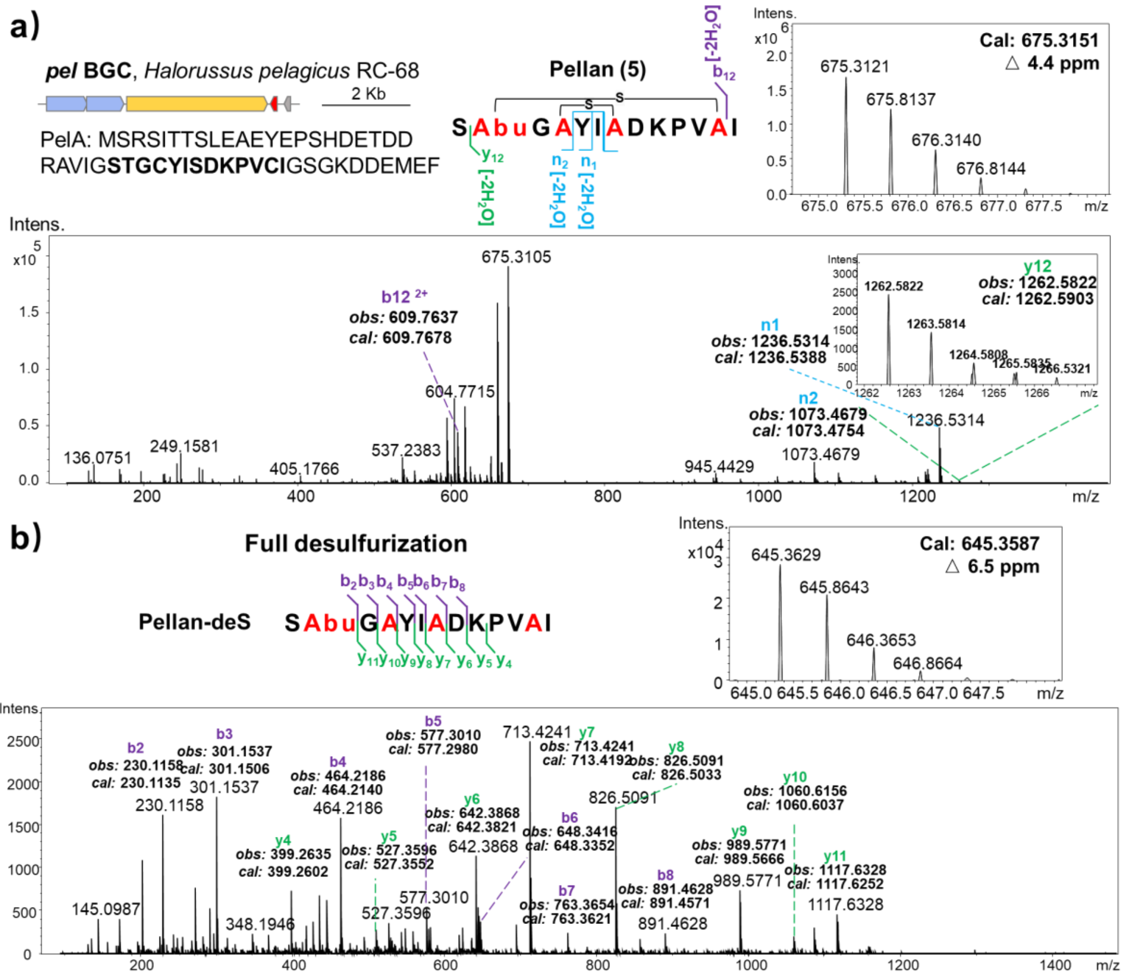

c)

| Ion | Pellan (5) Obs m/z | Pellan (5) Calc m/z | Pellan (5) △ppm | Pellan-deS Obs m/z | Pellan-deS Calc m/z | Pellan-deS △ppm | Ion | Pellan (5) Obs m/z | Pellan (5) Calc m/z | Pellan (5) △ppm | Pellan-deS Obs m/z | Pellan-deS Calc m/z | Pellan-deS △ppm |
|---|---|---|---|---|---|---|---|---|---|---|---|---|---|
| | | | | | | | y12 | 1262.5822 | 1262.5903 | 6.4 | | | |
| b2 | | | | 230.1158 | 230.1135 | 10.0 | y11 | | | | 1117.6328 | 1117.6252 | 6.8 |
| b3 | | | | 301.1537 | 301.1506 | 10.3 | y10 | | | | 1060.6156 | 1060.6037 | 11.2 |
| b4 | | | | 464.2186 | 464.2140 | 9.9 | y9 | | | | 989.5771 | 989.5666 | 10.6 |
| b5 | | | | 577.3010 | 577.2980 | 5.2 | y8 | | | | 826.5091 | 826.5033 | 7.0 |
| b6 | | | | 648.3416 | 648.3352 | 9.9 | y7 | | | | 713.4241 | 713.4241 | 0.0 |
| b7 | | | | 763.3654 | 763.3621 | 4.3 | y6 | | | | 642.3868 | 642.3821 | 7.3 |
| b8 | | | | 891.4628 | 891.4571 | 6.4 | y5 | | | | 527.3596 | 527.3552 | 8.3 |
| | | | | | | | y4 | | | | 399.2635 | 399.2602 | 8.3 |
| b12 | 609.7637 | 609.7678 | 6.7 | | | | | | | | | | |
| n1 | 1236.5314 | 1236.5388 | 6.0 | | | | | | | | | | |
| n2 | 1073.4679 | 1073.4754 | 7.0 | | | | | | | | | | |

**Supplementary Figure 10. LC-MS/MS analysis of heterologously expressed and desulfurized classical lanthipeptide pellan (5)**

The BGC *pel* was identified from *H. pelagicus* RC-68 (a). A peptide signal, $[M+2H]^{2+}$ = 675.3121 (calculated $[M+2H]^{2+}$ = 675.3151, Δ = 4.4 ppm), indicated putative lanthipeptide products (a, c). Based on MS1 analysis, the putative modification involves the loss of two molecules of $H_2O$ in the core region of PelA, indicating the presence of two thioether rings. The desulfurization product of pellan (**5**), pellan-deS, confirmed the entire core peptide and the modification residues of Thr2, Cys4, Ser7 and Cys12. Consequently, pellan (**5**) was hypothesized to have interwound or bicyclic rings, supported by MS1 and desulfurization analyses (S<u>T</u>G<u>C</u>YI<u>S</u>DKPV<u>C</u>I). However, MS/MS fragments of $n_1$ (m/z = 1236.5314 $[M+H]^+$, calculated $[M+H]^+$ = 1236.5388, Δ = 6.0 ppm) and $n_2$ (m/z = 1073.4679 $[M+H]^+$, calculated $[M+H]^+$ = 1073.4754, Δ = 7.0 ppm)) with the loss of 'I' and 'IY' residues within the rings (S<u>T</u>G<u>C</u>**YI**<u>S</u>DKPV<u>C</u>I) supported an interwind structure instead of a bicycle structure. Therefore, the thioether crosslinks of pellan (**5**) were deduced to be present in Thr2-Cys12, and Ser7-Cys4.

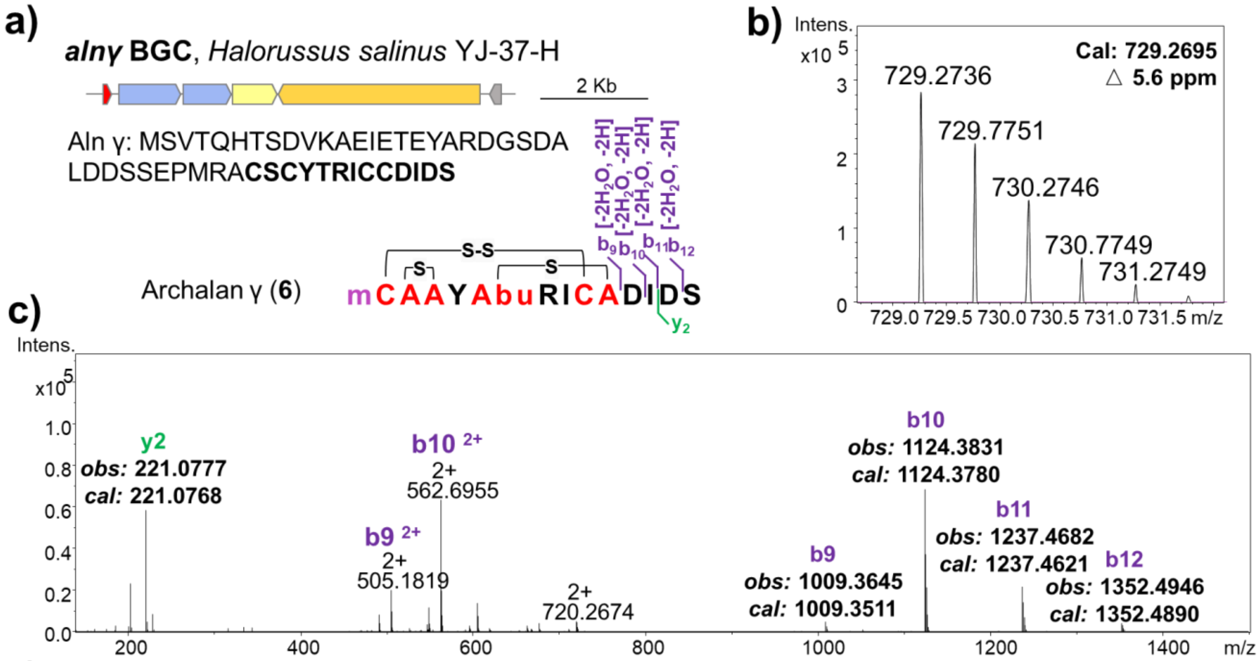

d)

| Archalan γ (**6**) | | | |
|---|---|---|---|
| Ion | Obs m/z | Calc m/z | △ppm |
| b12 | 1352.4946 | 1352.4890 | 4.1 |
| b11 | 1237.4682 | 1237.4621 | 4.9 |
| b10 | 1124.3831 | 1124.3780 | 4.5 |
| b9 | 1009.3645 | 1009.3511 | 13.3 |
| y2 | 221.0777 | 221.0768 | 4.1 |

**Supplementary Figure 11. LC-MS/MS analysis of heterologously expressed classical lanthipeptide archalan γ (6)**

The BGC *alnγ* was identified from *H. salinus* YJ-37-H (a), with a peptide signal of $[M+2H]^{2+}$ = 729.2736 (calculated $[M+2H]^{2+}$ = 729.2695, Δ = 5.6 ppm), indicating a putative lanthipeptide product (b-d). Archalan γ (**6**) was found to be produced by both recombinant and wild-type strains. The MS/MS fragments observed were limited to the amino acids outside the large cyclic rings ($b_9$-$b_{12}$), predicting with the fragment 'DIDS' of the core peptide Alnγ (CSCYTRICC<u>DIDS</u>). Through a combination of MS1 data, the modification likely involved one methylation and the loss of two $H_2O$ molecules and two hydrogens in the core region (b). The crosslinks were deduced by analyzing chemical bond breakdown reactions with tandem LC-MS/MS, as detailed in this study (Supplementary Figs. 12-17).

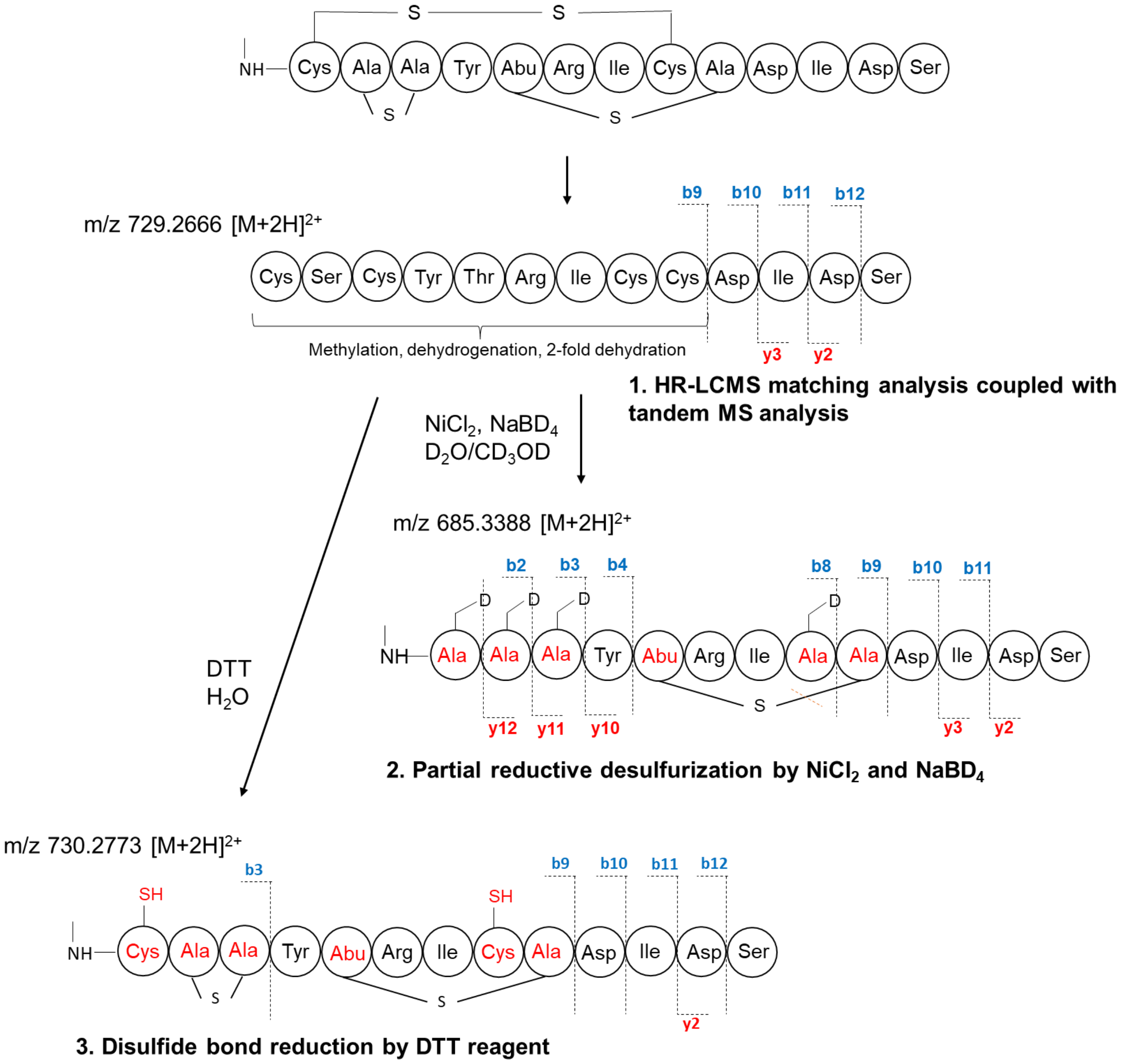

#### Supplementary Figure 12. Characterization of the C-S crosslinks in archalan γ (6)

The tandem mass spectrum of archalan γ displayed a linear DIDS motif in the C-terminus, consistent with the predicted core peptide sequence (CSCYTRICCDIDS). To figure out the C-S crosslinks in archalan γ, $NaBD_4$ and $D_2O/CD_3OD$ were employed for partial desulfurization. A mass signal of m/z 685.3388 $[M+2H]^{2+}$ was identified in the partially desulfurized product, indicating the reduction and desulfurization of one C-S crosslink while the other remained intact. Tandem mass analysis revealed that residues Cys1, Ser2, Cys3, and Cys8 were reduced to Ala, and a C-S crosslinking in Thr5 and Cys9. Subsequently, DTT reduction was employed to determine the position of a disulfide bond. The tandem mass spectrum of m/z 730.2773 $[M+2H]^{2+}$ showed the presence of a disulfide bond between Cys1 and another Cys residue. Further $MS^n$ analysis confirmed the S-S bond between Cys1 and Cys8. Altogether, these clues indicated the presence of two thioether rings in Thr5 and Cys9, Ser2 and Cys3, and a disulfide bond in Cys1 and Cys8.

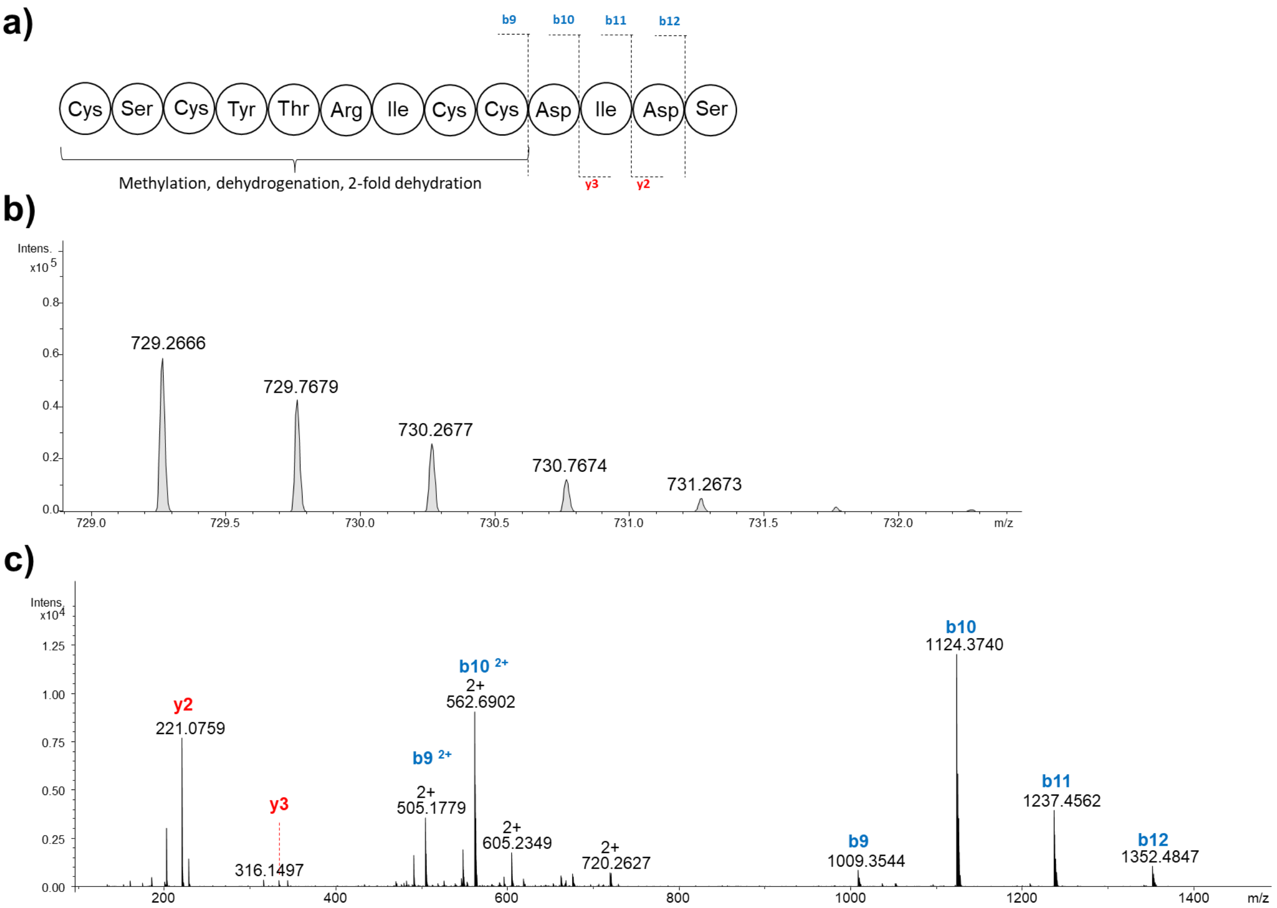

d)

| Ion | Obs m/z | Calc m/z | △ppm | Ion | Obs m/z | Calc m/z | △ppm |
|---|---|---|---|---|---|---|---|
| b9 | 1009.3544 | 1009.3511 | 3.3 | | | | |
| b10 | 1124.3740 | 1124.3780 | 3.6 | y3 | 334.1598 | 334.1609 | 3.3 |
| b11 | 1237.4562 | 1237.4621 | 4.8 | y2 | 221.0759 | 221.0768 | 4.1 |
| b12 | 1352.4847 | 1352.4890 | 3.2 | | | | |

#### Supplementary Figure 13. LC-MS/MS analysis of archalan γ (6)

a) The amino acid sequence of archalan γ is presented with b and y ions, along with the location of the thioether ring. LC-MS/MS was used to fragment archalan γ in order to confirm the amino acid sequence by examination of the fragmentation patterns. Major fragment ions of the doubly charged precursor ion, *m/z* 729.2666 (b), are annotated with their b/y ion identity (c), and the amino acid residues deduced from fragment ions are labeled in blue/red, respectively. d) The observed and calculated b/y ions are listed in the table.

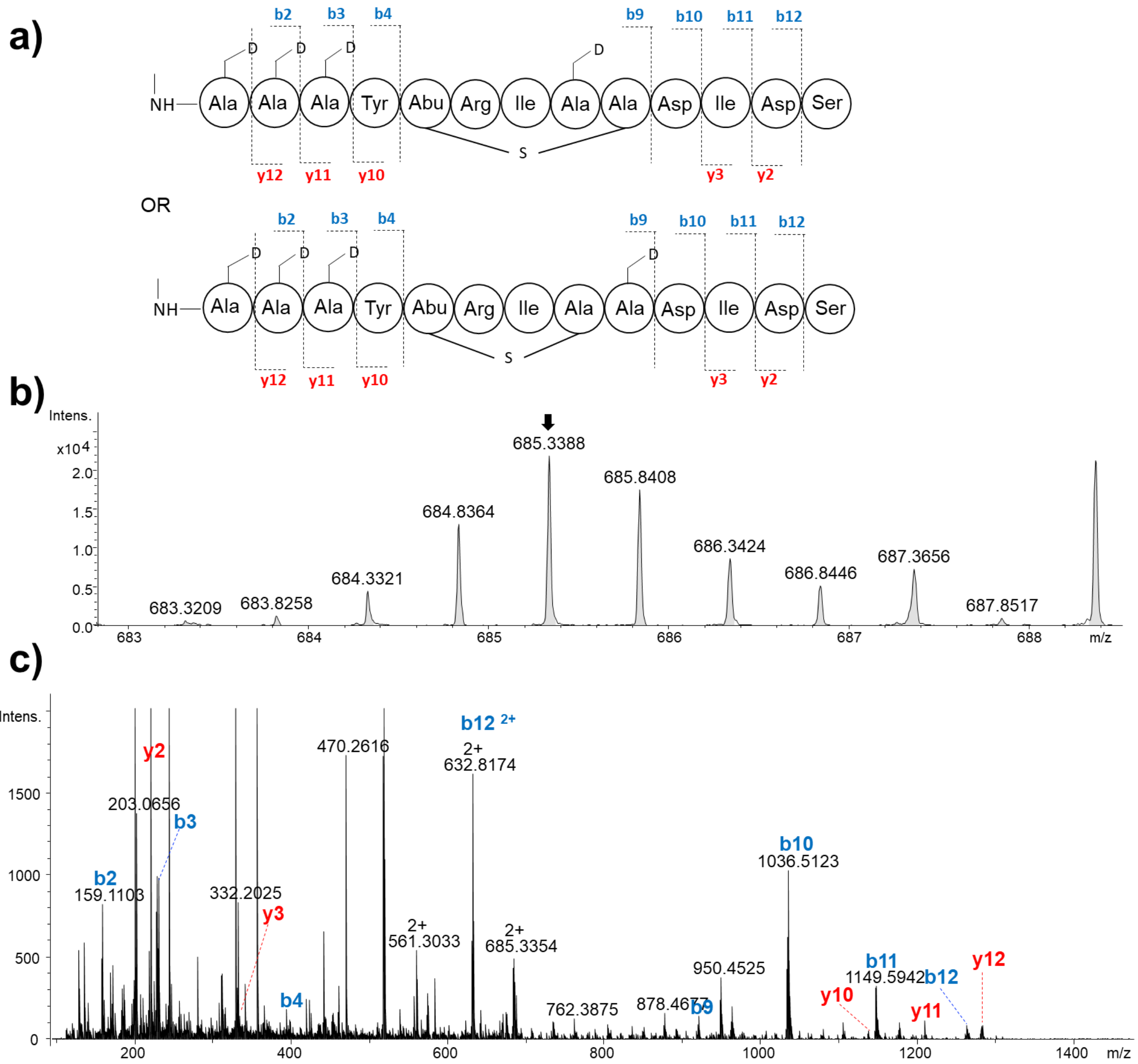

| Ion | Obs m/z | Calc m/z | △ppm | Ion | Obs m/z | Calc m/z | △ppm |
|---|---|---|---|---|---|---|---|
| | | | | y12 | 1283.6034 | 1283.6128 | |
| b2 | 159.1103 | 159.1097 | 3.8 | y11 | 1211.5621 | 1211.5694 | 6.0 |
| b3 | 231.1527 | 231.1531 | 1.7 | y10 | 1139.5159 | 1139.5260 | 8.9 |
| b4 | 394.2208 | 394.2164 | 11.2 | | | | |
| b9 | 921.4916 | 921.4913 | 0.3 | | | | |
| b10 | 1036.5123 | 1036.5182 | 5.7 | y3 | 334.1612 | 334.1609 | 0.9 |
| b11 | 1149.5942 | 1149.6023 | 7.0 | y2 | 221.0764 | 221.0768 | 1.8 |
| b12 | 1264.6161 | 1264.6292 | 10.4 | | | | |

**Supplementary Figure 14. LC-MS/MS analysis of partial desulfurized archalan γ (6)**

a) Two possibilities of the partial desulfurized archalan γ, which was generated from partial reductive desulfurization by $NiCl_2$ and $NaBD_4$ in $D_2O/CD_3OD$ (1:1, v:v), are presented with b and y ions marked as well as the presumed location of the thioether ring. LC-MS/MS was used to fragment the partially desulfurized product to confirm the ring position by examination of the fragmentation patterns. Major fragment ions of the doubly charged precursor ion, *m/z* 685.3388 (b), are annotated with their b/y ion identity (c), and the amino acid residues deduced from fragment ions are labeled in blue/red, respectively. d) The observed and calculated b/y ions are listed in the table.

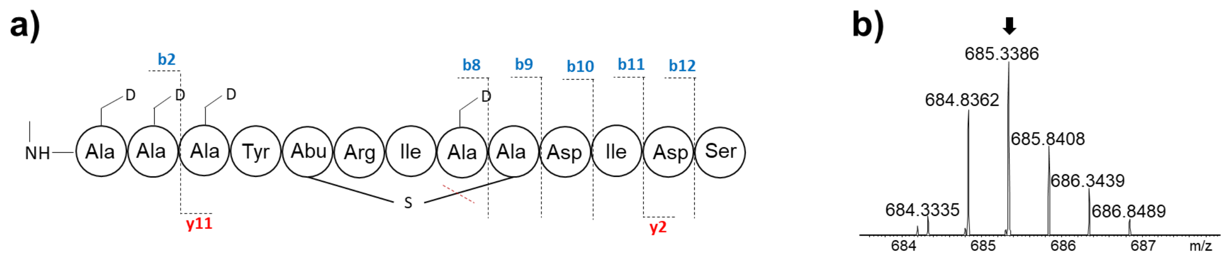

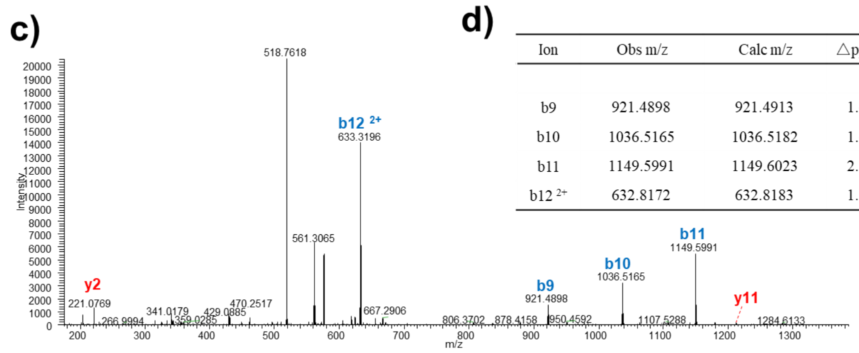

d)

| Ion | Obs m/z | Calc m/z | △ppm | Ion | Obs m/z | Calc m/z | △ppm |
|---|---|---|---|---|---|---|---|
| | | | | y11 | 1211.5686 | 1211.5694 | 0.7 |
| b9 | 921.4898 | 921.4913 | 1.6 | | | | |
| b10 | 1036.5165 | 1036.5182 | 1.6 | | | | |
| b11 | 1149.5991 | 1149.6023 | 2.8 | y2 | 221.0769 | 221.0768 | 0.5 |
| b12 $^{2+}$ | 632.8172 | 632.8183 | 1.7 | | | | |

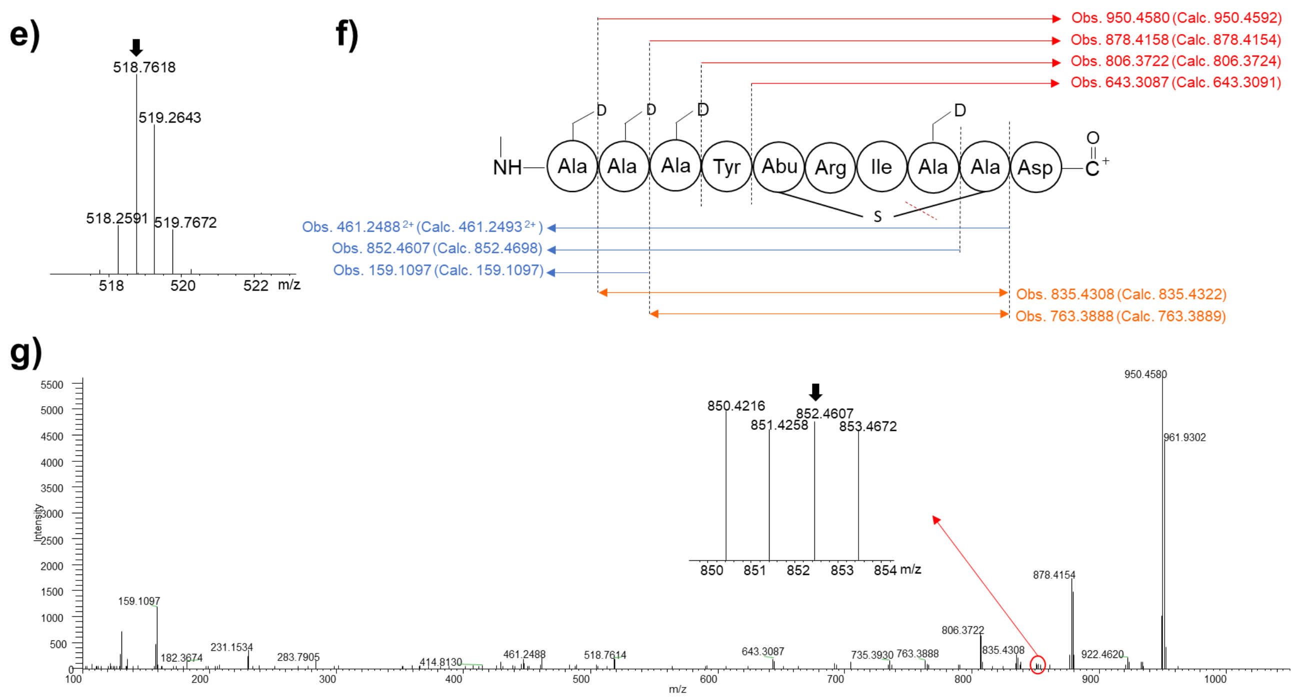

**Supplementary Figure 15. MS$^n$ spectrum of partial desulfurized archalan γ (6)**

a) The amino acid sequence of partial desulfurized archalan γ is presented with b and y ions marked as well as the location of the thioether ring which was generated from partial reductive desulfurization by $NiCl_2$ and $NaBD_4$ in $D_2O/CD_3OD$ (1:1, v:v). LC-MS/MS was used to fragment the partially desulfurized product to confirm the ring position by examination of the fragmentation patterns. Major fragment ions of the doubly charged precursor ion, *m/z* 685.3386 (b), are annotated with their b/y ion identity (c), and the amino acid residues deduced from fragment ions are labeled in blue/red, respectively. d) The observed and calculated b/y ions are listed in the table. The key MS$^3$ fragments of MS/MS ion *m/z* 518.7618 (e) are presented with observed and calculated mass (f) which existed in the MS$^3$ spectrum (g).

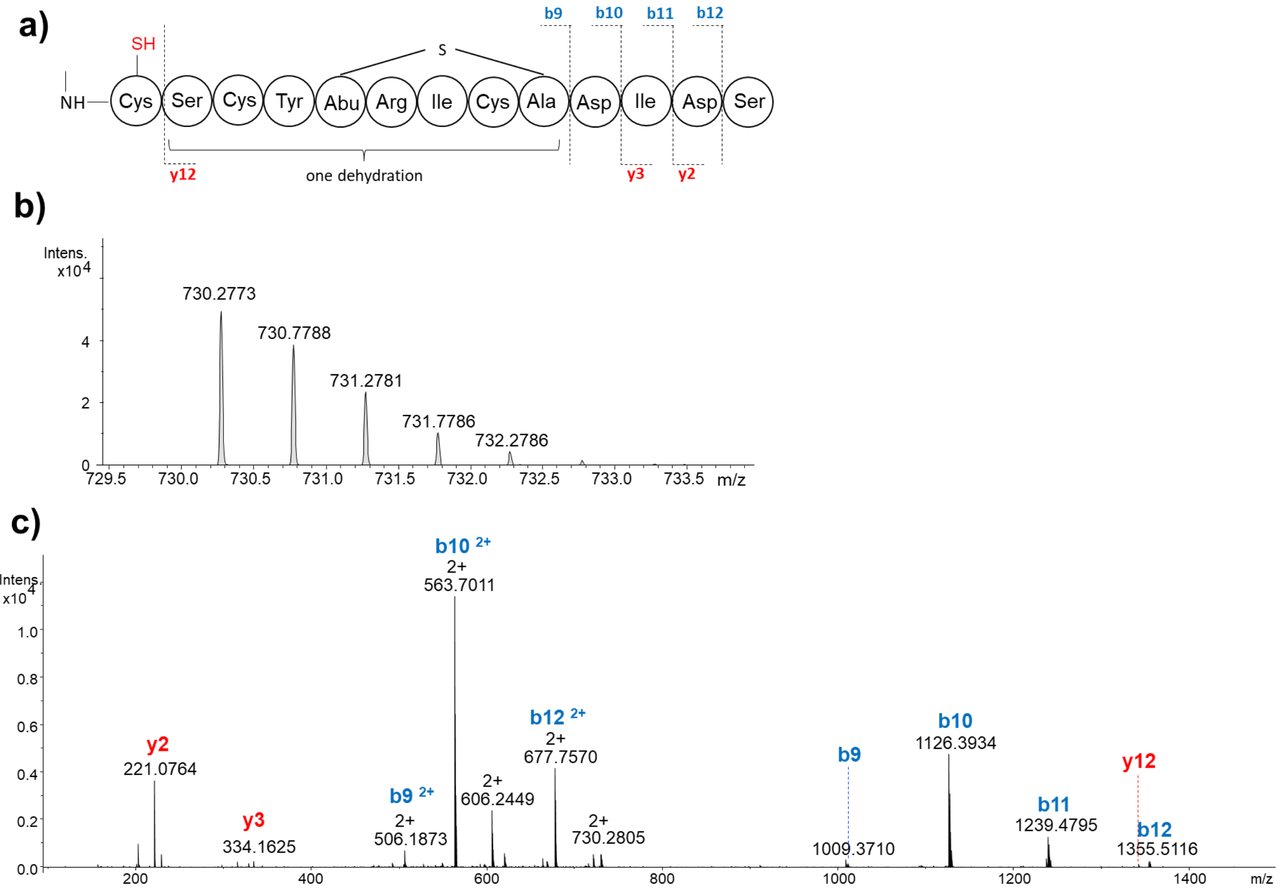

d)

| Ion | Obs m/z | Calc m/z | △ppm | Ion | Obs m/z | Calc m/z | △ppm |
|---|---|---|---|---|---|---|---|
| | | | | y12 | 1342.5239 | 1342.5225 | 1.0 |
| b9 | 1011.3667 | 1011.3667 | 0.0 | | | | |
| b10 | 1126.3934 | 1126.3937 | 0.3 | y3 | 334.1625 | 334.1609 | 4.8 |
| b11 | 1239.4795 | 1239.4778 | 1.4 | y2 | 221.0764 | 221.0768 | 1.8 |
| b12 | 1354.5116 | 1354.5047 | 5.1 | | | | |

**Supplementary Figure 16. LC-MS/MS analysis of DTT reduction product of archalan γ (6)**

a) The amino acid sequence of the DTT reduction product of archalan γ is presented with b and y ions, along with confirmed methylation and reduction of Cys1. LC-MS/MS was used to confirm the disulfide bond position by examination of the fragmentation patterns. Major fragment ions of the doubly charged precursor ion, *m/z* 730.2773 (b), are annotated with their b/y ion identity (c), and the amino acid residues deduced from fragment ions are labeled in blue/red, respectively. d) The observed and calculated b/y ions are listed in the table.

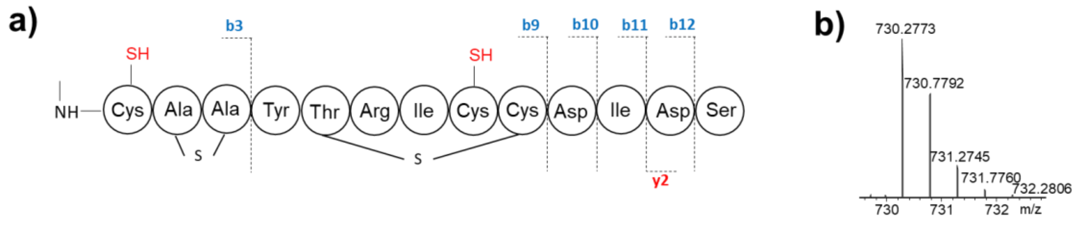

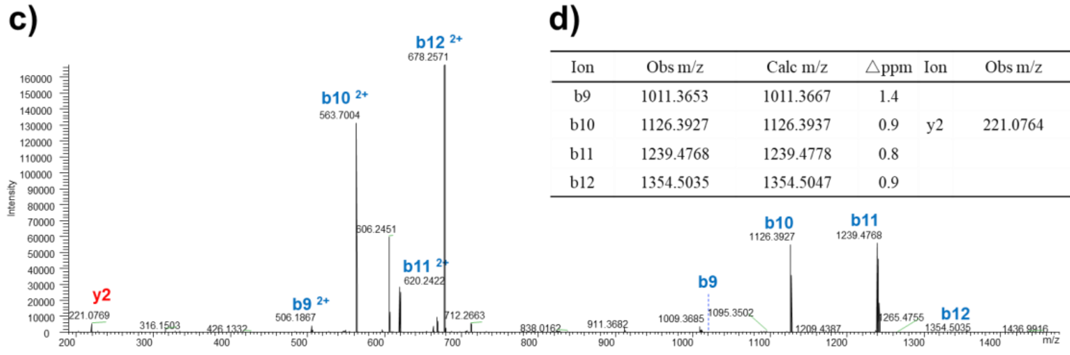

d)

| Ion | Obs m/z | Calc m/z | △ppm | Ion | Obs m/z | Calc m/z | △ppm |
|---|---|---|---|---|---|---|---|
| b9 | 1011.3653 | 1011.3667 | 1.4 | | | | |
| b10 | 1126.3927 | 1126.3937 | 0.9 | y2 | 221.0764 | 221.0768 | 1.8 |
| b11 | 1239.4768 | 1239.4778 | 0.8 | | | | |
| b12 | 1354.5035 | 1354.5047 | 0.9 | | | | |

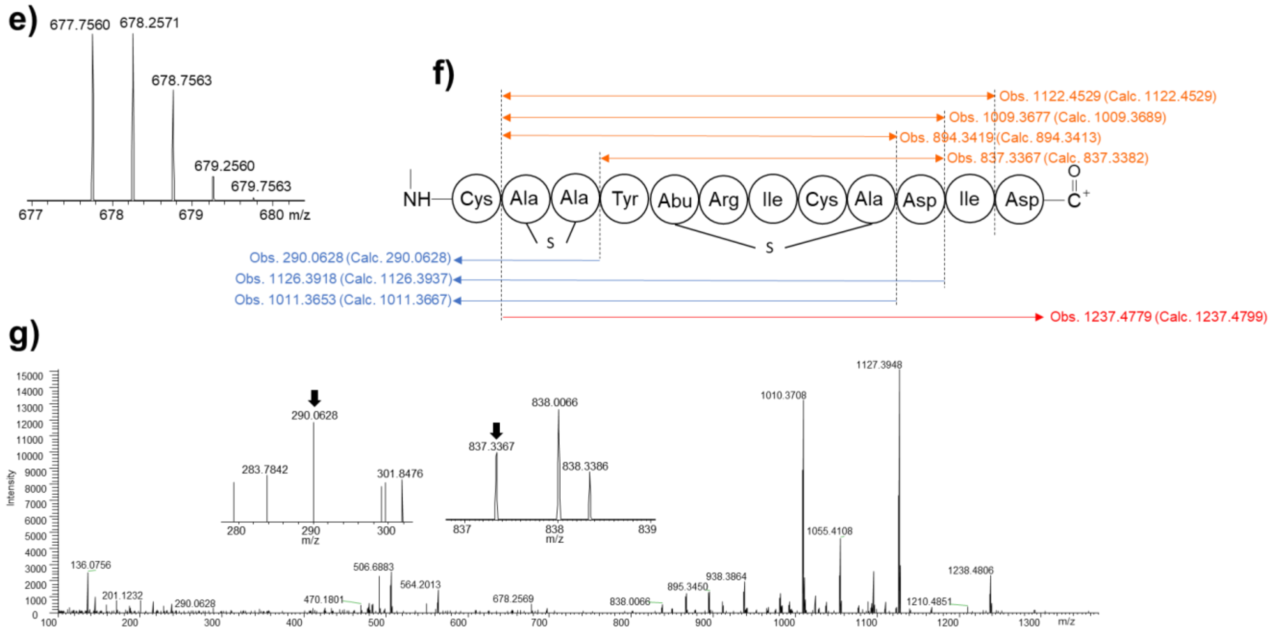

**Supplementary Figure 17. MS$^n$ spectrum of DTT reduction product of archalan γ (6)**

a) The amino acid sequence of the DTT reduction product of archalan γ is presented with b and y ions, along with the assumed location of the confirmed reduction of Cys1 and Cys8. Major fragment ions of the doubly charged precursor ion, *m/z* 730.2773 (b), are annotated with their b/y ion identity (c), and the amino acid residues deduced from fragment ions are labeled in blue/red, respectively. d) The observed and calculated b/y ions are listed in the table. The key MS$^3$ fragments of MS$^2$ ion, *m/z* 677.7560 (e), are presented with observed and calculated mass (f) which existed in the MS$^3$ spectrum (g). All the clues indicated that the S-S bond occurred between Cys1 and Cys8, and the crosslink between Cys3 and Ser2.

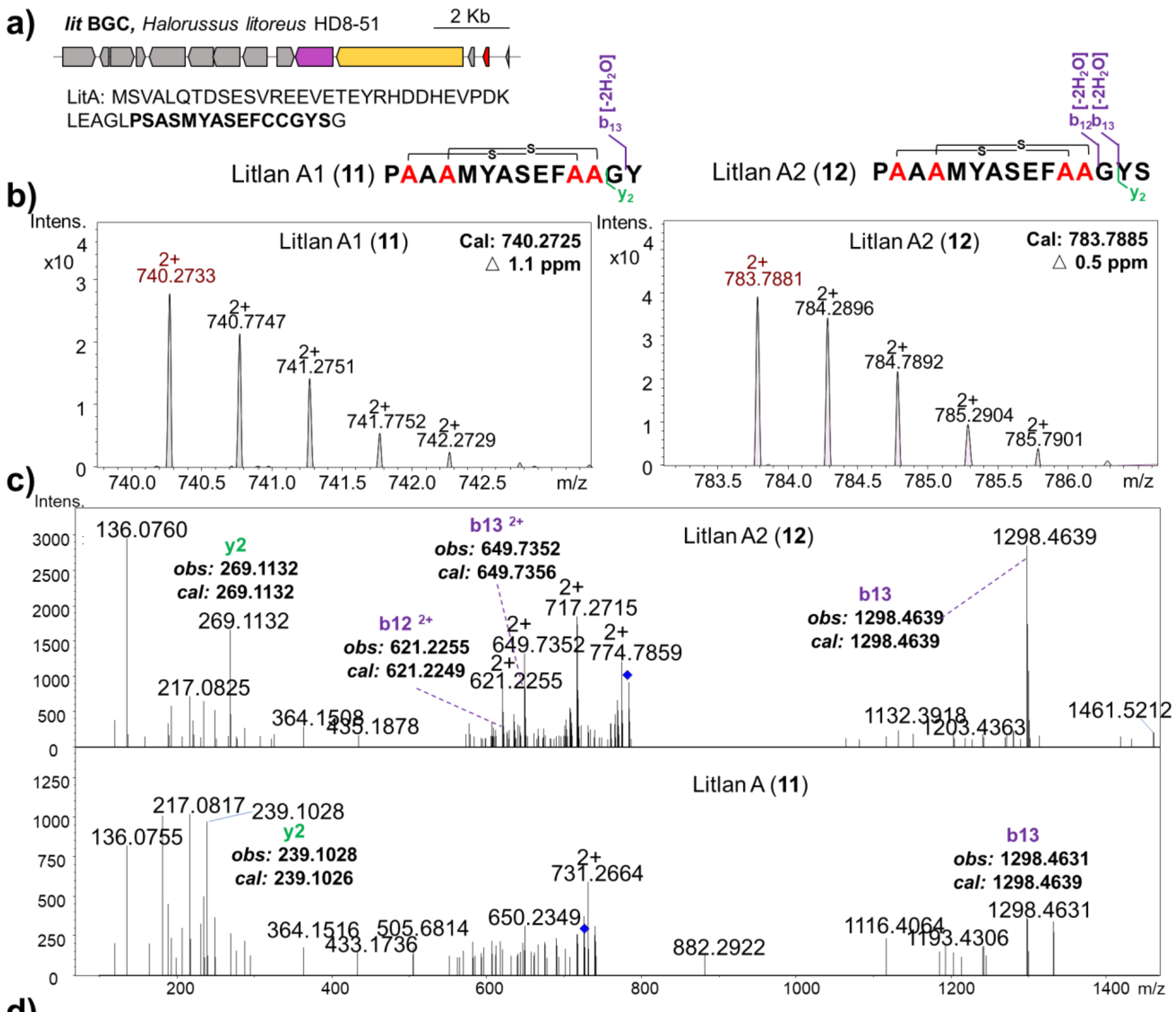

| Ion | Litlan A1 (11) Obs m/z | Calc m/z | △ppm | Litlan A2 (12) Obs m/z | Calc m/z | △ppm |
|---|---|---|---|---|---|---|
| b12 | | | | 621.2255 | 621.2249 | 1.0 |
| b13 | 1298.4631 | 1298.4639 | 0.6 | 1298.4639 | 1298.4639 | 0.0 |
| y2 | 239.1028 | 239.1026 | 0.8 | 269.1132 | 269.1132 | 0.0 |

**Supplementary Figure 18. LC-MS/MS analysis of heterologously expressed classical lanthipeptide litlans (11-12)**

The MS1 analysis suggested a putative modification of litlan A1 (**11**) and A2 (**12**), which involved the loss of two molecules of $H_2O$, pointing to the presence of two thioether rings, akin to archalan β. Examination of the cyclization of litlans revealed that only the C-terminal overhang amino acids outside the thioether rings could be identified, consistent with the LC-MS/MS analysis results of archalan β. Litlan A2 features an additional Ser in the C-terminus compared to A1, as evidenced by key MS/MS fragments of $b_{12}$, $b_{13}$, and $y_2$. This variation may be attributed to the influence of extracellular proteases from the host genome, as no discernible protease is encoded within the BGC. In the SSN analysis, LitA from BGC *lit* and Alnβ from BGC *alnβ* were in the same cluster. Therefore, we assigned the thioether rings of litlans to be in Ser2-Cys11 and Ser4-Cys12, based on the verified structure of archalan β.

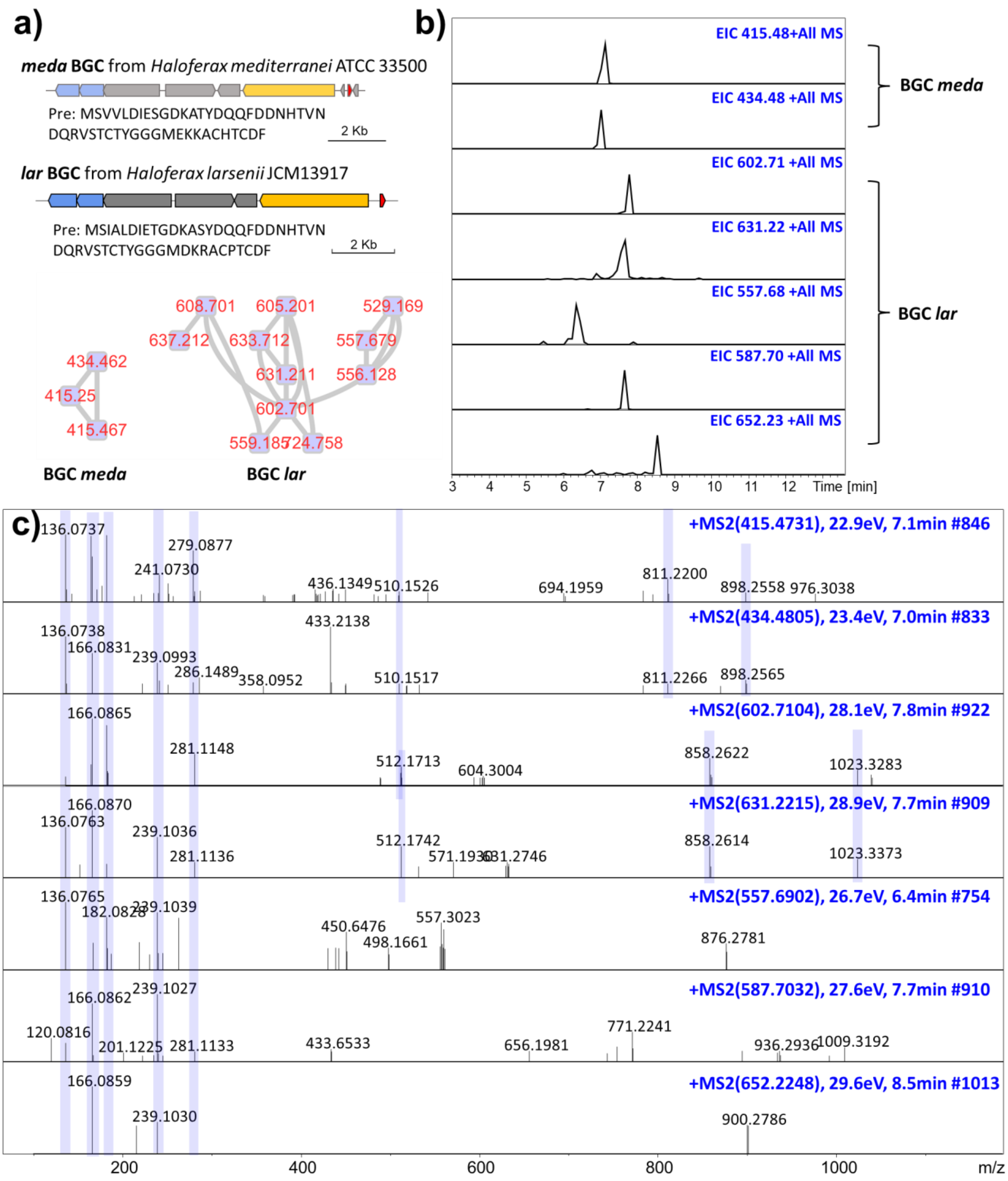

**Supplementary Figure 19. LC-MS/MS analysis of noncanonical diamino dicarboxylic (DADC) lanthipeptides from the metabolites of BGCs *meda* and *lar***

Two sets of unusual peptide mass signals were detected in the metabolites of BGCs *meda* and *lar*(a-c), present in both wild-type and heterologous expression extracts, thereby confirming their association with the respective BGCs (a). Global Natural Products Social Molecular Networking (GNPS)[6] analysis of the crude extracts revealed analogs (a) with many similar MS/MS fragments (c) that cannot be linked as a linear amino acid sequence and aligned with any known modifications (such as methylation, dehydration, desulfurization, and hydroxylation) of the presumed lanthipeptide precursors as classical lanthipeptides (b). In addition, the detection of a mass difference of 56 Da ($[M+3H]^{3+}$ = 415.48 and 434.48 Da, $[M+2H]^{2+}$ = 602.71 and 631.22 Da) and several common MS/MS fragments (e.g., 136.08, 166.09, 239.10, 811.22 or 858.26), indicated the hypothesis that these analogs were derived from the same BGCs, differing by a Gly residue. Ultimately, we isolated and characterized a representative compound from each group using NMR to elucidate their structures. They exhibited diamino-dicarboxylic termini and were named DADC lanthipeptides, a previously unrecognized subfamily within the lanthipeptide class. Furthermore, other analogs were elucidated through LC-MS analysis based on their confirmed structures.

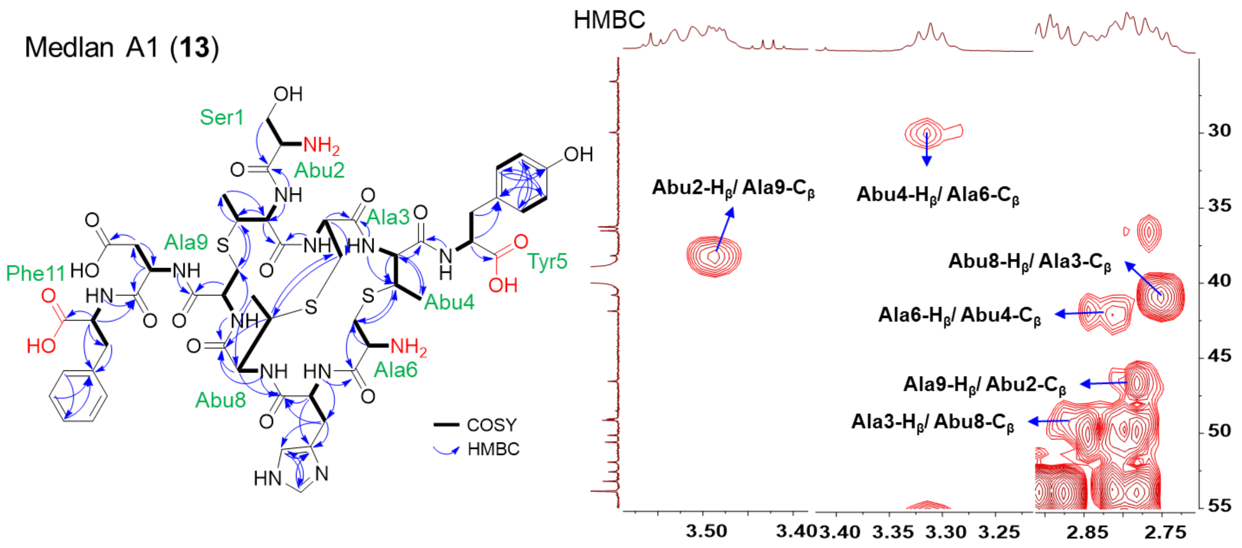

**Supplementary Figure 20. Structure elucidation of purified archaeal lanthipeptide medlan A1 (13)**

The mass signal of medlan A1 (**13**), observed as $[M+3H]^{3+}$ = 415.4769, was predicted with a molecular formula of $C_{52}H_{69}N_{13}O_{17}S_3$ (*cal*: $[M+3H]^{3+}$ = 415.4772, Δ = 0.7 ppm). The purified compound **13**, obtained from a 20 L culture broth of the wild-type strain with a yield of 3.2 mg, was elucidated using extensive NMR analysis in DMSO-$d_6$. The observation of many exchangeable amide NH protons ($\delta_H$ 7.6-9.2 ppm) occurred in the $^1$H-NMR and carbonyl carbons ($\delta_C$ 167.0-173.0 ppm) in $^{13}$C-NMR spectra supported the peptidic nature of **13** (Supplementary Fig. 42 and Supplementary Table 3). Three pairs of HMBC signals of Abu2-H$_\beta$/Ala9-C$_\beta$ ($\delta_H$ = 3.50 ppm to $\delta_C$ = 38.3 ppm) and Ala9-H$_\beta$/Abu2-C$_\beta$ ($\delta_H$ = 2.78 ppm to $\delta_C$ = 46.5 ppm), Ala6-H$_\beta$/Abu4-C$_\beta$ ($\delta_H$ = 2.82 ppm to $\delta_C$ = 41.9 ppm) and Abu4-H$_\beta$/Ala6-C$_\beta$ ($\delta_H$ = 3.32 ppm to $\delta_C$ = 30.1 ppm), and Ala3-H$_\beta$/Abu8-C$_\beta$ ($\delta_H$ = 2.86, 2.81 ppm to $\delta_C$ = 49.1 ppm) and Abu8-H$_\beta$/Ala3-C$_\beta$ ($\delta_H$ = 2.75 ppm to $\delta_C$ = 40.9 ppm) suggested three methyllanthionine rings in Thr2-Cys9, Thr4-Cys6 and Cys3-Thr8. The detection of eleven NH protons and carbonyl carbons while only with nine pairwise HMBC correlations indicated the occurrence of two free amino ($\delta_H$ = 8.23 ppm for Ser1 and $\delta_H$ = 8.28 ppm for Ala6) and two free carboxylic groups ($\delta_C$ = 172.8 ppm for Tyr5 and $\delta_C$ = 172.6 ppm for Phe11), which is the reason why we named this kind of unforeseen lanthipeptide as diamino-dicarboxylic (DADC) lanthipeptides. Combining HSQC, COSY, and HMBC data, compound **13** was thereby deduced to contain 1 × Ser, 1 × Tyr, 1 × His, 1 × Asp, 1 × Phe, and 3 ×MeLan (Supplementary Fig. 42). The stereochemistry of **13** was confirmed with L configuration in all unmodified amino acids, along with two LL-MeLan (2*R*,3*R*,6*R*) configurations, and one in a DL-MeLan (2*S*,3*S*,6*R*) configuration by comparing the standard samples in advanced Marfey's analysis (Supplementary Fig. 9 and Supplementary Table 4). However, the precise configuration assignment could only be deduced by the configuration of two mutants of larlan A2 (**14**) (Supplementary Fig. 9 and Supplementary Table 4). Given that MedA and LarA belong to the same SSN family, the findings above enable us to propose the conformation of thioether rings in **13**: Thr2-Cys9, Thr8-Cys3 are in the LL-MeLan (2*R*,3*R*,6*R*) configuration and Thr4-Cys6 in DL-MeLan (2*S*,3*S*,6*R*) configuration.

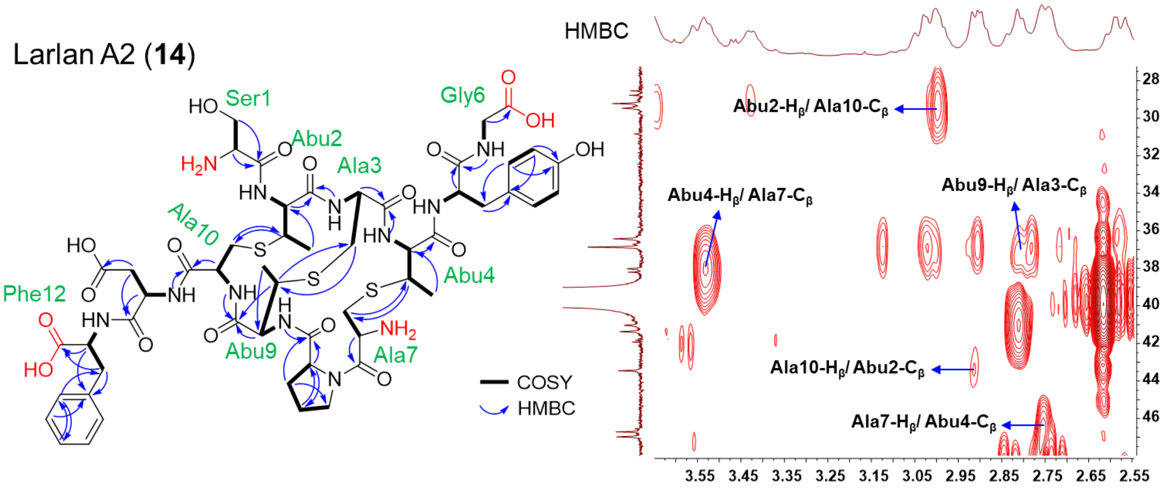

**Supplementary Figure 21. Structure elucidation of purified archaeal lanthipeptide larlan A2 (14)**

The mass signal of larlan A2 (**14**), observed as $[M+2H]^{2+}$ = 631.2194, was predicted with a molecular formula of $C_{53}H_{72}N_{12}O_{18}S_3$ (*cal*: $[M+2H]^{2+}$ = 631.2198, $\Delta$ = 0.6 ppm). Compound **14**, purified from the culture broth of heterologously expressed genes *larAMTBCDEFHGIJ* in recombinant *H. volcanii* H1424 with a yield of 3.5 mg, was elucidated using extensive NMR analysis in DMSO-$d_6$. The observation of many exchangeable amide NH protons ($\delta_H$ 7.6-9.1 ppm) occurred in the $^1$H-NMR and carbonyl carbons ($\delta_C$ 165.0-173.0 ppm) in $^{13}$C-NMR spectra supported the peptidic nature of **14** (Supplementary Fig. 43 and Supplementary Table 3). Three pairs of HMBC signals of Abu2-$H_\beta$/Ala10-$C_\beta$ ($\delta_H$ = 2.98 ppm to $\delta_C$ = 29.4 ppm), Ala10-$H_\beta$/Abu2-$C_\beta$ ($\delta_H$ = 2.90 ppm to $\delta_C$ = 43.5 ppm), Ala7-$H_\beta$/Abu4-$C_\beta$ ($\delta_H$ = 2.74 ppm to $\delta_C$ = 46.7 ppm), Abu4-$H_\beta$/Ala7-$C_\beta$ ($\delta_H$ = 3.52 ppm to $\delta_C$ = 38.1 ppm), and Abu9-$H_\beta$/Ala3-$C_\beta$ ($\delta_H$ = 2.80 ppm to $\delta_C$ = 36.9 ppm) suggested three methyllanthionine rings in Thr2-Cys10, Thr4-Cys7 and Cys3-Thr9. The identification of ten NH protons and twelve carbonyl carbons, along with only ten pairwise HMBC correlations, suggested the presence of two free carboxylic groups ($\delta_C$ = 171.5 ppm for Gly6 and $\delta_C$ = 173.0 ppm for Phe12) and two free $NH_2$ protons without signals, confirming its classification of DADC lanthipeptides. Combining HSQC, COSY, and HMBC data, compound **14** was thereby deduced to contain 1 × Ser, 1 × Tyr, 1 × Pro, 1 × Asp, 1 × Phe, 1 × Gly and 3 ×MeLan. The stereochemistry of **14** was confirmed with L configuration in all unmodified amino acids by advanced Marfey's analysis (Supplementary Table 4). As for three MeLan residues in larlan A2, two additional single mutants of larlar A2 (T2S and T4S) were purified to give a clear stereo structure annotation of each MeLan residue. By comparing with four MeLan standards[4, 5], the thioether rings between Thr2 and Cys10, Thr9 and Cys3 are characterized in the LL-MeLan (2*R*,3*R*,6*R*) configuration, while another ring between Thr4 and Cys7 in DL-MeLan (2*S*,3*S*,6*R*) configuration in larlan A2 (Supplementary Fig. 9 and Supplementary Table 4).

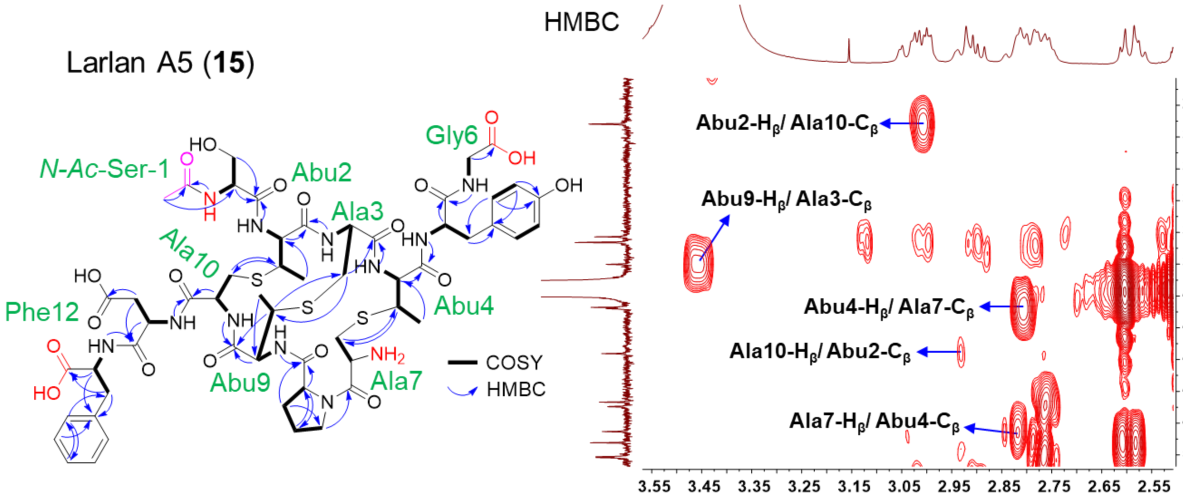

**Supplementary Figure 22. Structure elucidation of purified archaeal lanthipeptide larlan A5 (15)**

The mass signal of larlan A5 (**15**), observed as $[M+2H]^{2+}$ = 652.2301, was predicted with a molecular formula of $C_{55}H_{74}N_{12}O_{19}S_3$ (*cal*: $[M+2H]^{2+}$ = 652.2251, Δ = 7.7 ppm). The purified compound **15**, obtained from the same extracts as **14** with a yield of 2.0 mg, was elucidated using extensive NMR analysis in DMSO-$d_6$. The observation of many exchangeable NH protons ($\delta_H$ 7.6-9.1 ppm) occurred in the $^1$H-NMR and carbonyl carbons ($\delta_C$ 166.0-173.0 ppm) in $^{13}$C-NMR spectra supported the peptidic nature of **15** (Supplementary Fig. 44 and Supplementary Table 3). The HMBC signals of Abu2-$H_\beta$/Ala10-$C_\beta$ ($\delta_H$ = 3.01 ppm to $\delta_C$ = 29.2 ppm) and Ala10-$H_\beta$/Abu2-$C_\beta$ ($\delta_H$ = 2.93 ppm to $\delta_C$ = 43.6 ppm), Ala7-$H_\beta$/Abu4-$C_\beta$ ($\delta_H$ = 2.83 ppm to $\delta_C$ = 48.5 ppm) and Abu4-$H_\beta$/Ala7-$C_\beta$ ($\delta_H$ = 2.80 ppm to $\delta_C$ = 40.7/40.8 ppm), and Abu9-$H_\beta$/Ala3-$C_\beta$ ($\delta_H$ = 3.45 ppm to $\delta_C$ = 38.0 ppm) suggested three methyllanthionine rings in Thr2-Cys10, Thr4-Cys7 and Cys3-Thr9. As an analog of **14**, **15** exhibited two free carboxylic groups ($\delta_C$= 171.1 ppm for Gly6 and $\delta_C$= 172.5 ppm for Phe12), along with one more acetylation ($\delta_C$= 169.9 ppm for CO and $\delta_H$= 1.86 ppm, $\delta_C$= 22.5 ppm for $CH_3$) on the Ser1 and one free $NH_2$ without signal, suggesting its classification of DADC lanthipeptides. Combining HSQC, COSY, and HMBC data, compound **15** was thereby deduced to contain 1 × *N-Ac*-Ser, 1 × Tyr, 1 × Pro, 1 × Asp, 1 × Phe, 1 × Gly and 3 ×MeLan. The stereochemistry of **15** was confirmed with L configuration in all unmodified amino acids by advanced Marfey's analysis (Supplementary Table 4). The remaining three MeLan residues are annotated and proposed based on the confirmed structure of **14**: the thioether rings between Thr2 and Cys10, Thr9 and Cys3 are characterized in LL-MeLan (2*R*,3*R*,6*R*) configurations, while another ring between Thr4 and Cys7 in DL-MeLan (2*S*,3*S*,6*R*) configuration (Supplementary Fig. 9 and Supplementary Table 4).

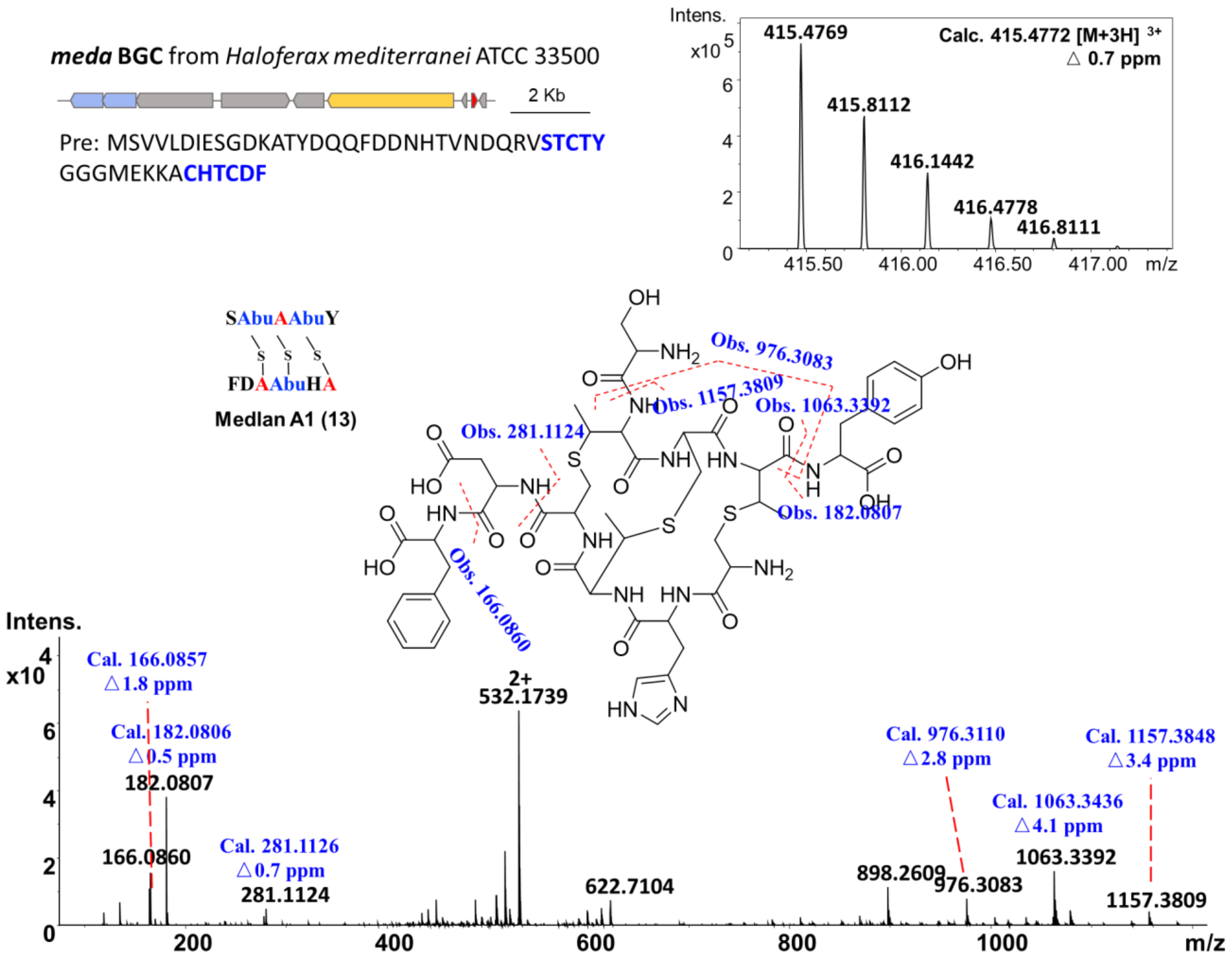

#### Supplementary Figure 23. LC-MS/MS analysis of medlan A1 (13)

Peptide signal $[M+3H]^{3+}$ = 415.4769 was putative lanthipeptide products linked to BGC *meda* identified from *H. mediterranei* ATCC35500. After confirmation by NMR data, medlan A1 (**13**) matched the putative precursor of *meda* BGC with the absence of partial amino acids within the thioether rings which haven't been discovered in the classical lanthipeptides. Its structure and LC-MS fragments are annotated based on the NMR results. It was impossible to elucidate their structure barely based on the MS data. At first, for the MS1 prediction, the core peptide of DADC lanthipeptides couldn't be predicted based on classical lanthipeptide biosynthetic rules which contain intact core peptides with modifications. Secondly, it contains diamino and dicarboxylic termini which could be fragmented by ESI causing short amino acid fragments from different termini (e.g. 'FD' from the second C-terminus, 'Y' from the first C-terminus, and 'S' from the first N-terminus). However, the MS/MS data still provided a confidential clue to link the final product with the putative precursors. Therefore, the NMR spectrum and LC-MS data of medlan A1 (**13**) served as a foundation for the predicted structure of the subsequent analog, medlan A2 (**16**).

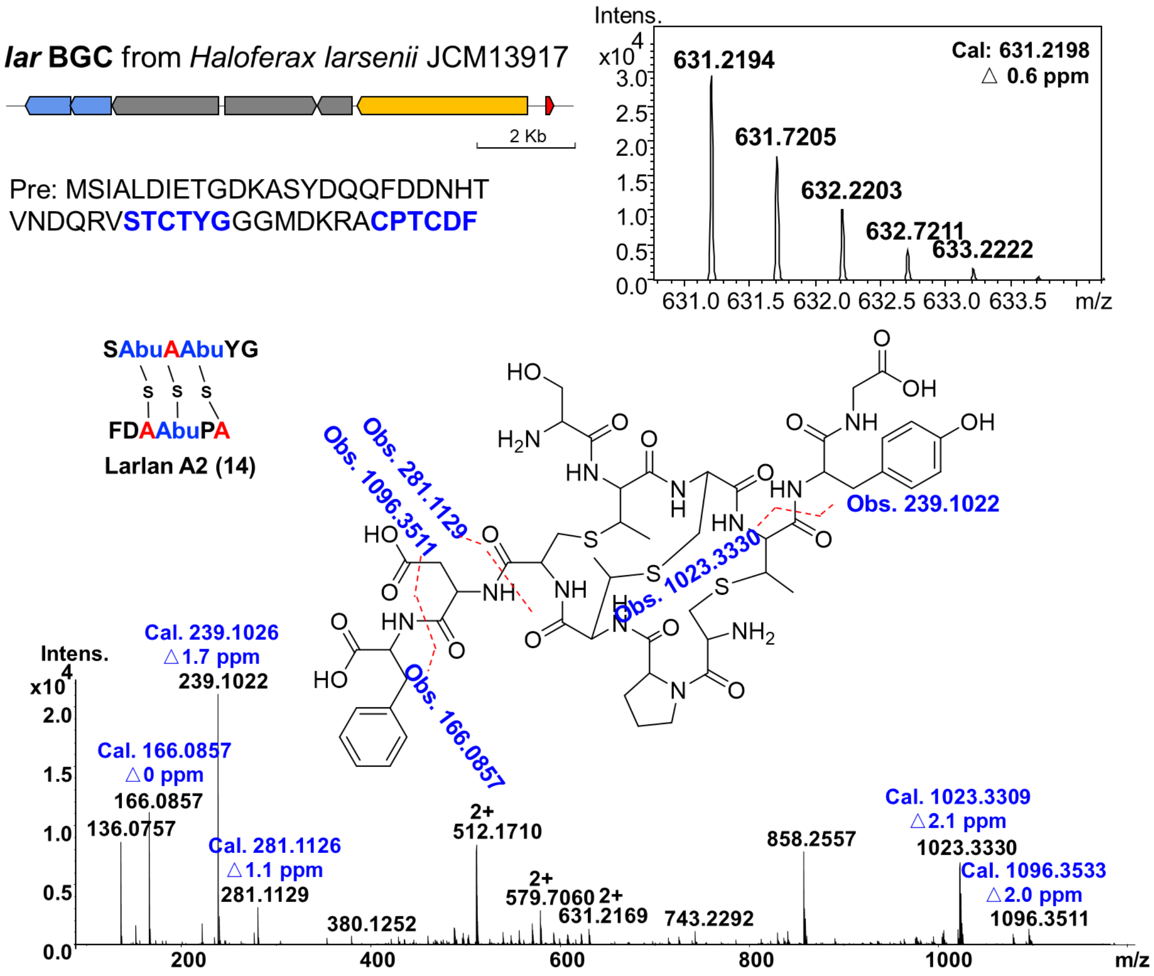

#### Supplementary Figure 24. LC-MS/MS analysis of larlan A2 (14)

BGC *lar* was identified from strain *H. larsenii* JCM 13917. The precursor peptides of both BGCs *lar* and *medb* belong to the same SSN cluster. The peptide signal $[M+2H]^{2+}$ = 631.2194, corresponding to larlan A2 (**14**), was identified as putative DADC lanthipeptide products linking to BGC *lar*. Through NMR analysis, the structure of **14** revealed diamino-dicarboxylic termini and three methyllanthionine rings, characteristics shared with medlans. Notably, the structure of larlan A2 supports the proposed structure of medlan A2, both showing an additional Gly residue following Tyr at the first C-terminus. In addition, larlan A2 serves as the foundation for predicting the structure of other following analogs, including larlans A1, A3 and A4.

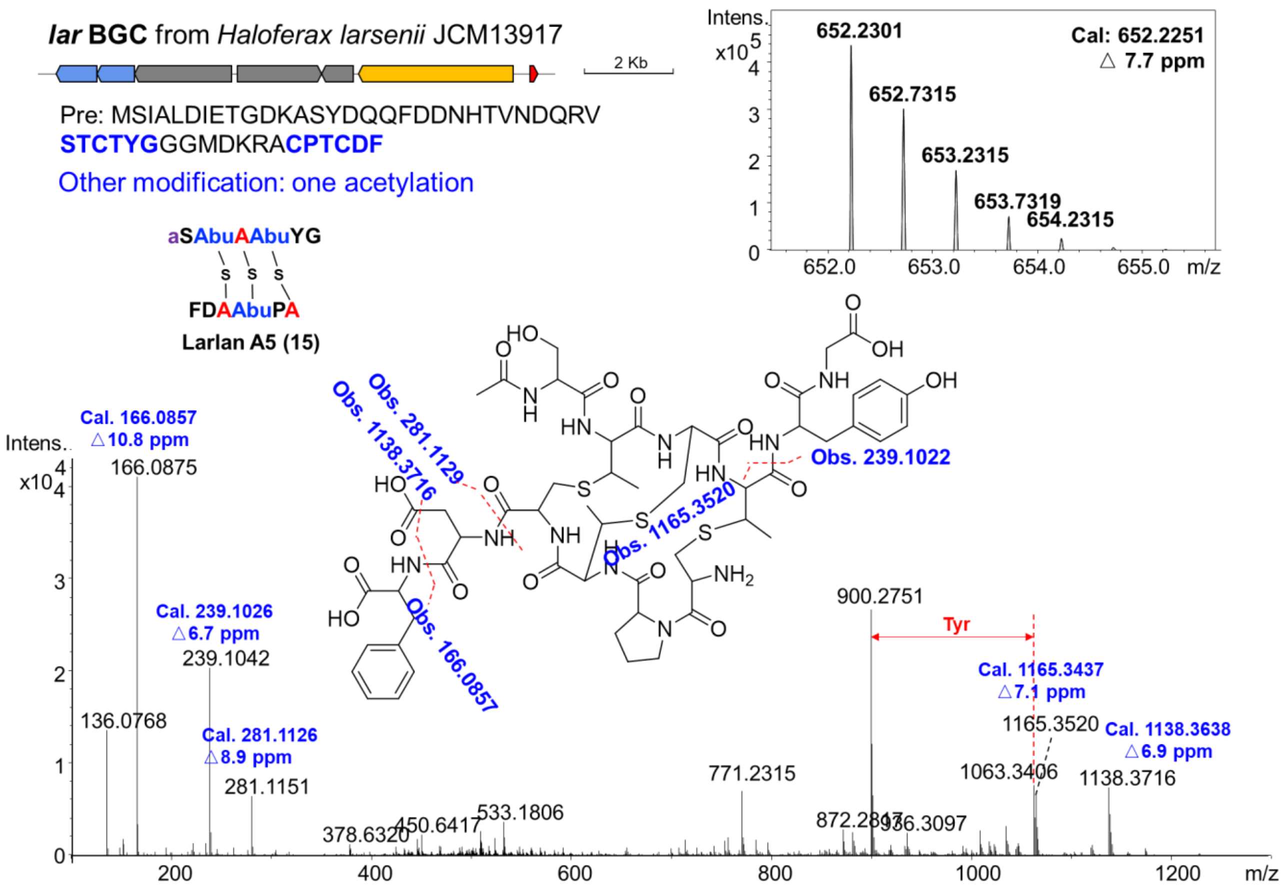

**Supplementary Figure 25. LC-MS/MS analysis of larlan A5 (15)**

Larlan A5 (**15**), $[M+2H]^{2+}$ = 652.2301, had a mass increase of 42 Da than larlan A2 (**14**), implying the addition of an acetylation modification. The MS/MS data indicated that while no modifications were observed at the C-termini, there was a variance at the N-termini, indicating acetylation at one of the two N-terminus. It was challenging to definitively determine the location of the acetylation solely based on MS/MS analysis. Consequently, the compound was purified to facilitate structural elucidation. The NMR spectrum determined the precise location of the acetylation on the first Ser residue at the first N-terminus.

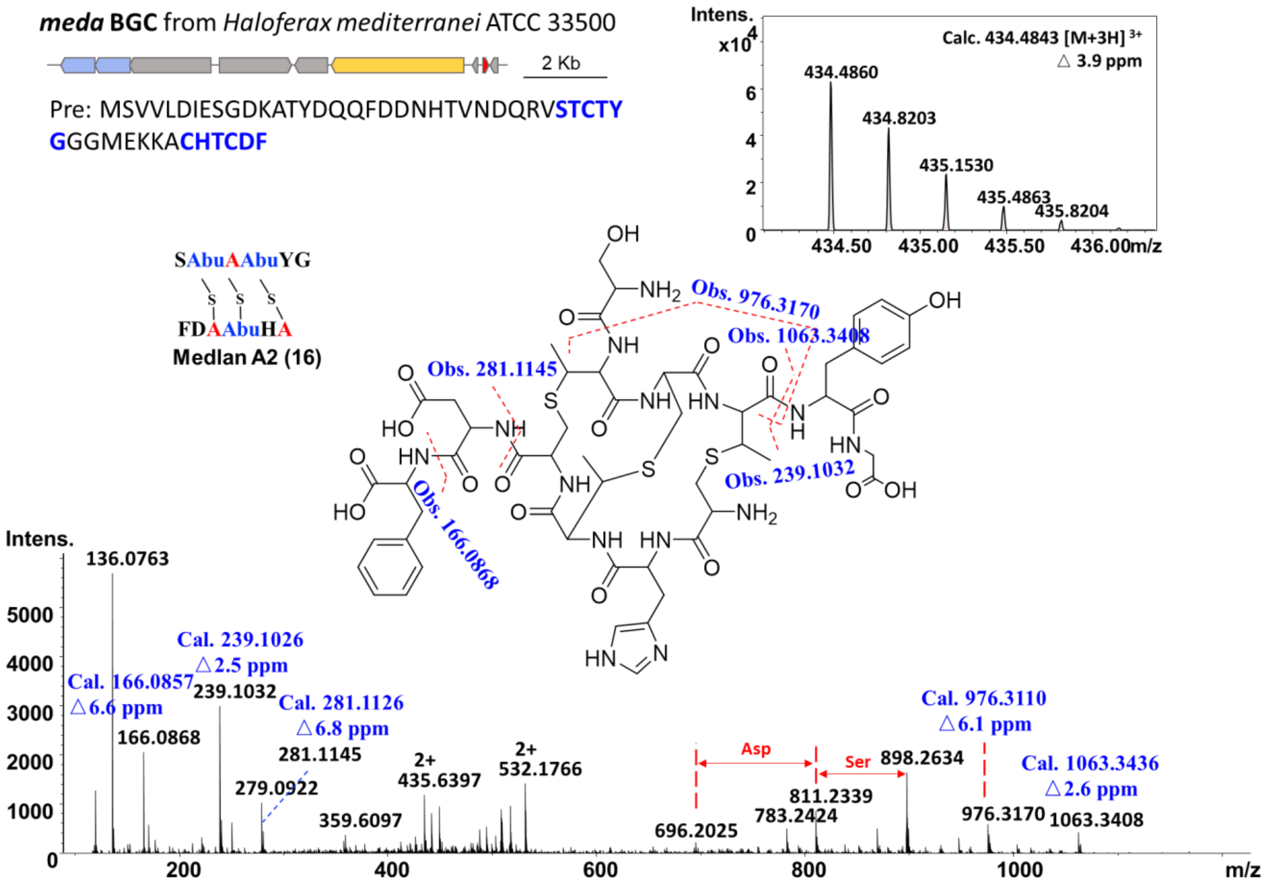

#### Supplementary Figure 26. LC-MS/MS analysis of medlan A2 (16)

Peptide signal $[M+3H]^{3+}$ = 434.4860 was putative DADC lanthipeptide product linked to BGC *meda*, serving as an analog of medlan A1 (**13**). Notably, this signal exhibited a mass increase of 57.0213 Da, which is believed to be a Gly residue. In the MS/MS analysis, fragments $[M+H]^{+}$ = 239.1032 in **16** and $[M+H]^{+}$ = 182.0807 in **13** displayed a mass shift of 57.0225 Da, supporting the presence of an additional Gly at the first C-terminus. This result aligned with the precursor amino acid sequence where Gly follows Tyr in the first peptide chain (SCTCTYG). Furthermore, other MS/MS fragments observed in **16** were identical to those in **13**, indicating no other differences in these regions. Based on the confirmed structure of **13**, **16** was proposed to include an additional Gly following Tyr as drawn above. This modification could potentially be attributed to the action of an unidentified aminopeptidase enzyme.

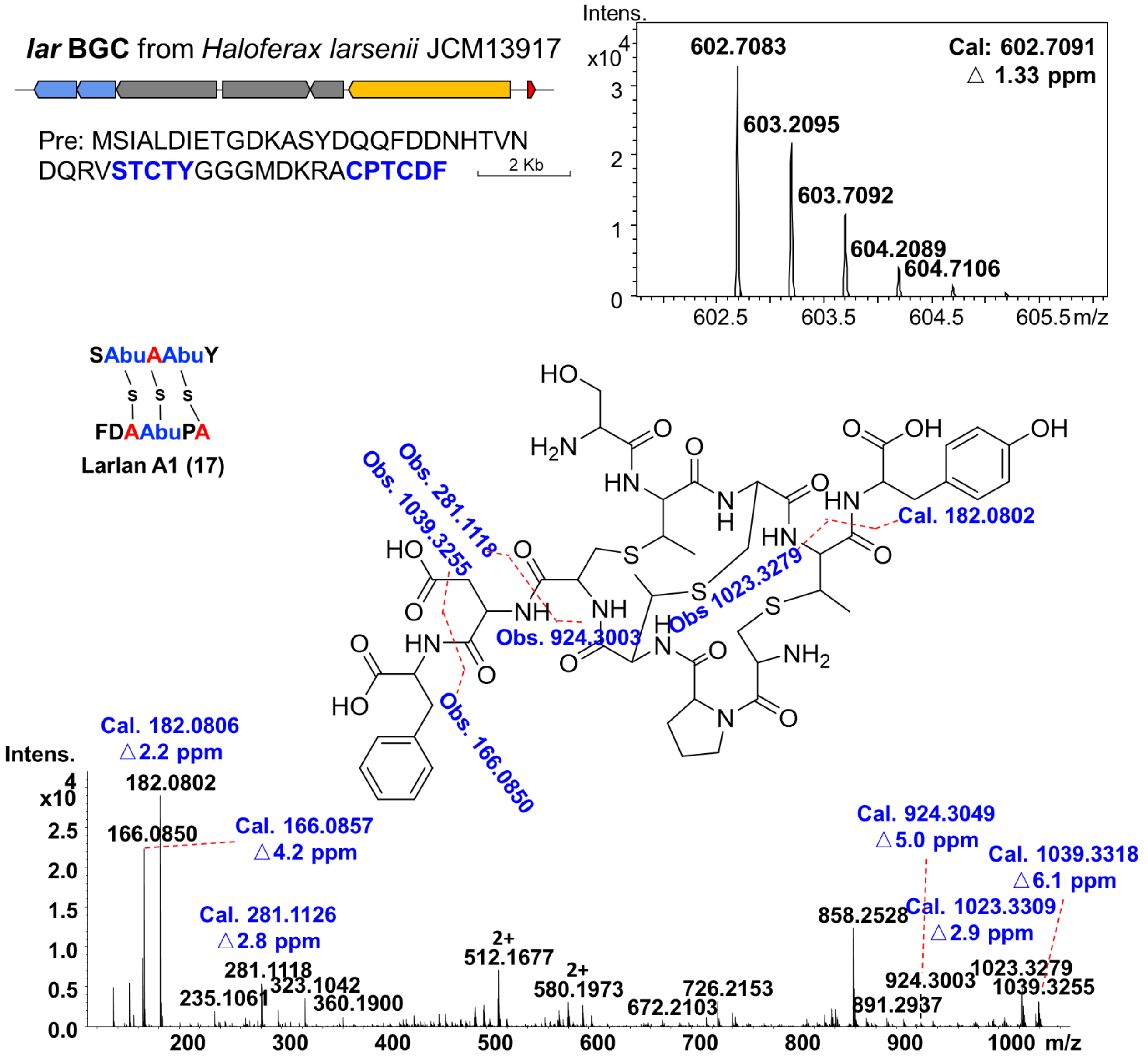

**Supplementary Figure 27. LC-MS/MS analysis of larlan A1 (17)**

Peptide signal $[M+2H]^{2+}$ = 602.7083 (larlan A1, **17**) was identified as an analog of larlan A2. The structure and LC-MS fragments of **17** were characterized based on the confirmed structure of larlan A2 (**14**). Similar to medlans, the key MS/MS fragment of $[M+H]^{+}$ = 182.0802 in **17** and $[M+H]^{+}$ = 239.1022 in **14** had the mass shift of 57.0220 Da, supporting a loss of Gly residue following Tyr in the first peptide chain. Other MS/MS fragments labeled in larlan A1 were consistent with those in larlan A2, suggesting no other differences in these regions.

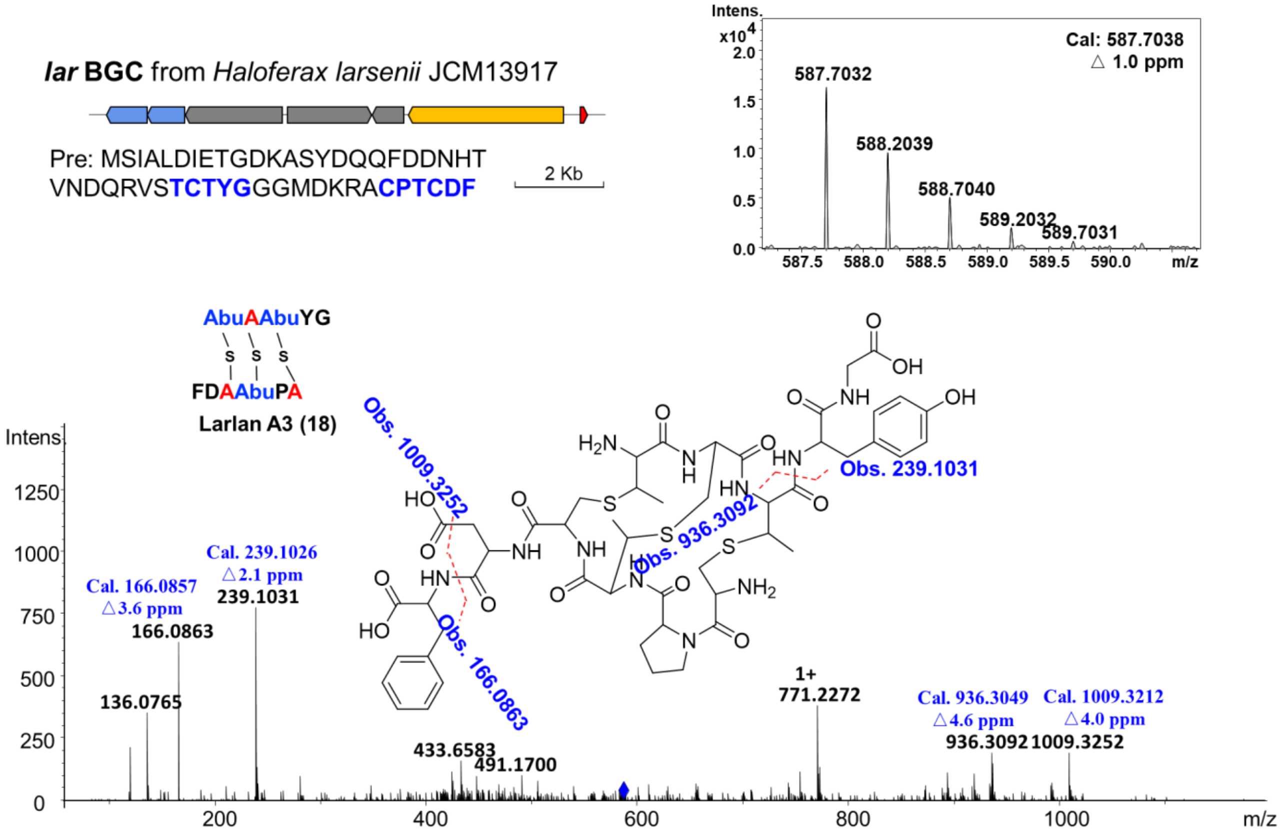

#### Supplementary Figure 28. LC-MS/MS analysis of larlan A3 (18)

Except for the distinction of a Gly residue between larlan A1 (**16**) and A2 (**14**), the analogs of larlans have exhibited variations in other modifications (such as methylation, dehydration, desulfurization, and hydroxylation) and amino acid compositions, as revealed by LC-MS/MS analysis. Three out of the four termini have different amino acid overhangs beyond the thioether ring, creating a series of analogs that could potentially be cleaved by unidentified aminopeptidases. Larlan A3 (**18**), a peptide signal of $[M+2H]^{2+}$ = 587.7032, showed a mass decrease of 87.0324 Da compared to larlan A2 ($[M+2H]^{2+}$ = 631.2194) based on the MS1 data, suggesting a variance attributable to a Ser residue (87.0320 Da). Additionally, the MS/MS fragments of C-termini were identical to those of larlan A2 and a mass shift occurred in the central ring portion, consistent with a Ser residue at the N-terminus of the core peptide. Thus, we propose the structure of larlan A3 to be similar to the confirmed structure of larlan A2, but lacking the Ser residue at the first N-terminus.

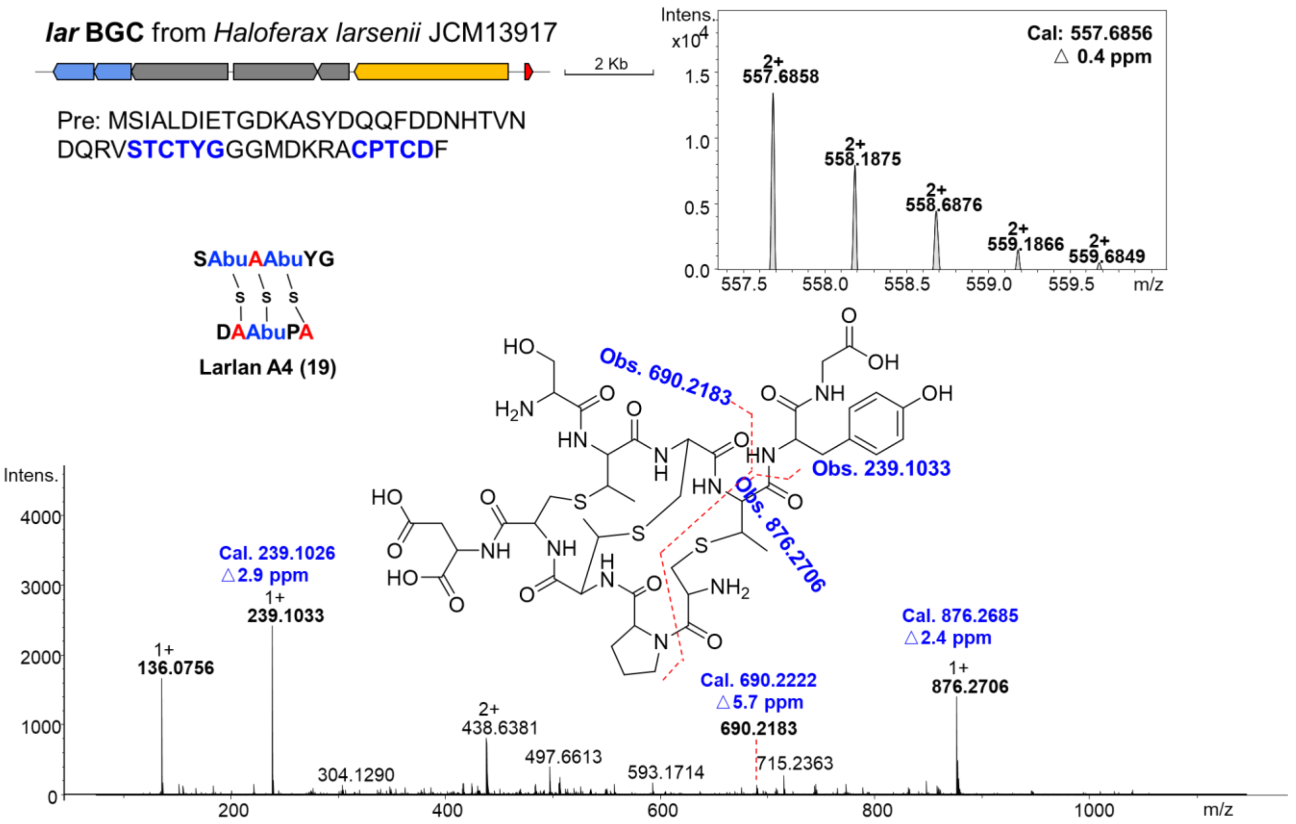

#### Supplementary Figure 29. LC-MS/MS analysis of larlan A4 (19)

Larlan A4 (**19**), $[M+2H]^{2+}$ = 557.6858, had a mass decrease of 147.0672 Da in comparison to larlan A2 ($[M+2H]^{2+}$ = 631.2194) as determined by the MS1 analysis, suggesting a difference corresponding to the Phe residue (147.0684). The MS/MS data revealed no mass shift at the C-terminus of the first chain, with differences observed in the region encompassing the second C-terminus and the first N-terminus, featuring a crucial fragment of $[M+H]^{+}$ = 690.2183. Moreover, there is only one Phe located at the second C-terminus and no other possible rearrangement to be a Phe residue for this part. Therefore, the structure of larlan A4 was deduced by combining MS/MS analysis, the biosynthetic rule, and the confirmed structure of larlan A2.

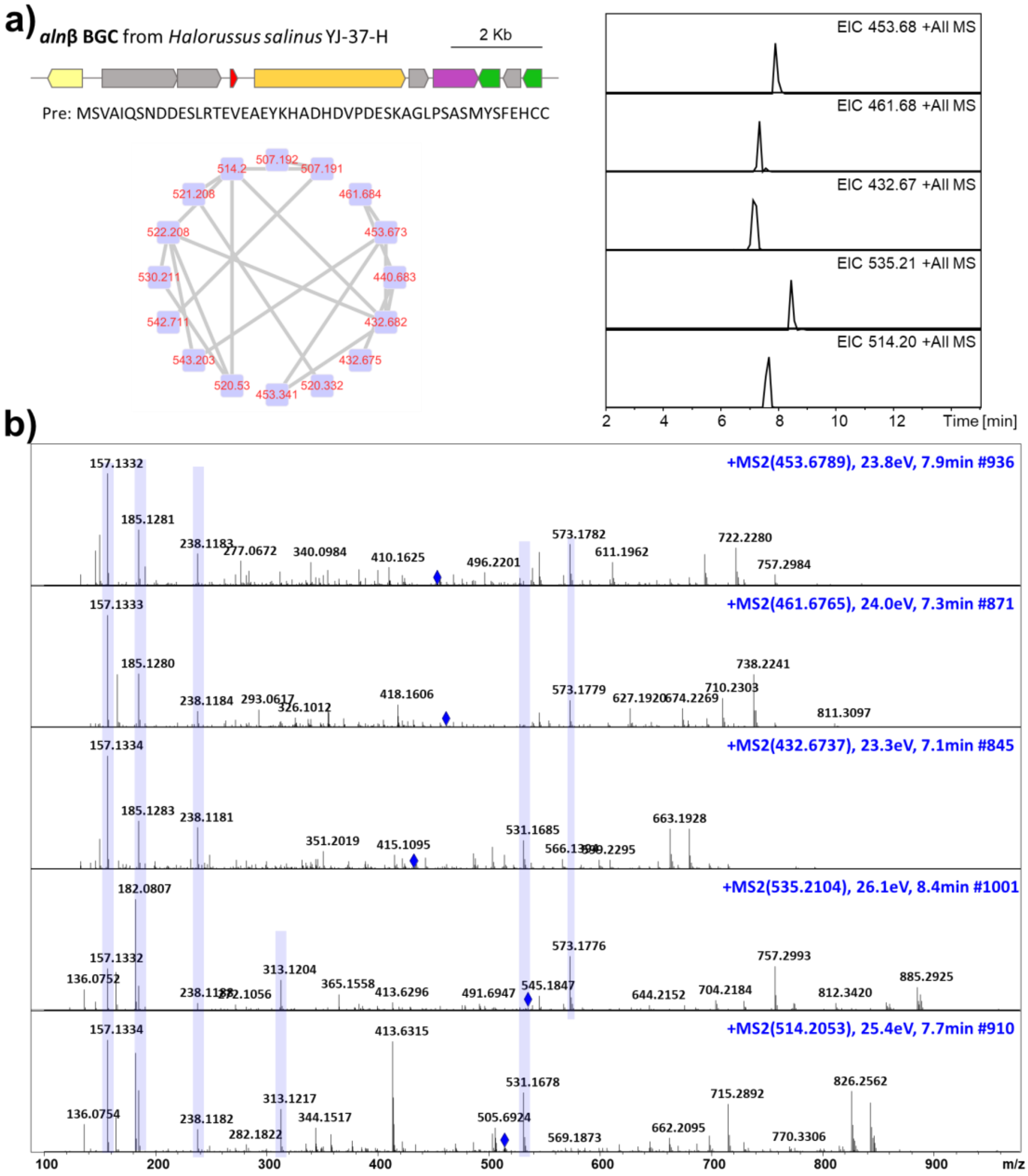

**Supplementary Figure 30. LC-MS/MS analysis of noncanonical DADC lanthipeptides produced by *H. salinus* YJ-37-H**

To find whether DADC lanthipeptides have appeared in other archaea strains, we analyzed the LC-MS data of lanthipeptide high-production strain *H. salinus* YJ-37-H which contained six lanthipeptide BGCs and produced archalans $\alpha$-$\gamma$[1]. The MS results also contained a set of mass signals resembling larlans and medlans, characterized by similar MS/MS fragments (a,b) that couldn't match the core region like classical lanthipeptides. Notably, we observed an incremental mass shift of 163.06 Da between m/z 514.20 $[M+2H]^{2+}$ and m/z 432.67 $[M+2H]^{2+}$, as well as between m/z 535.21 $[M+2H]^{2+}$ and m/z 453.68 $[M+2H]^{2+}$ (b), indicating the presence of a Tyr residue (163.06 Da). The GNPS analysis of crude extracts revealed structural similarities among these analogs (a) while their MS/MS fragments (b) couldn't be linearly linked as classical lanthipeptides. These analogs, akin to previous instances, exhibited additional modifications such as methylation, dehydration, desulfurization, and hydroxylation, along with variations in amino acid composition. Ultimately, we isolated and characterized a representative compound, archalan β3, which featured diamino-dicarboxylic termini as DADC lanthipeptides. Furthermore, other analogs archalan β4-7 were elucidated through LC-MS analysis based on the confirmed structures of archalan β3.

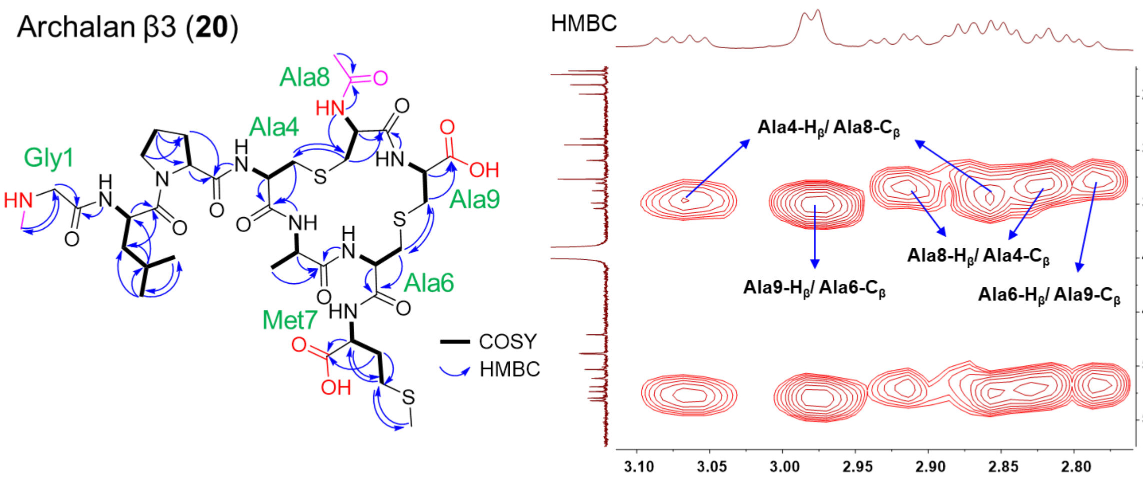

**Supplementary Figure 31. Structure elucidation of purified archaeal lanthipeptide archalan β3 (20)**

The mass signal of archalan β3 (**20**), observed as $[M+2H]^{2+}$ = 453.6800, was predicted with a molecular formula of $C_{36}H_{59}N_9O_{12}S_3$ (*cal*: $[M+2H]^{2+}$ = 453.6796, △0.9 ppm). The purified compound **20**, obtained from the fermentation broth of wild-type strain with a yield of 1.0 mg, was elucidated using extensive NMR analysis in DMSO-$d_6$. The observation of many exchangeable amide protons ($\delta_H$ 7.5-8.7 ppm) occurred in the $^1$H-NMR and carbonyl carbons ($\delta_C$ 165.0-173.0 ppm) in $^{13}$C-NMR spectra supported the peptidic nature of **20** (Supplementary Fig. 45 and Supplementary Table 3). The observation of two pairs of HMBC signals: Ala4-$H_\beta$/Ala8-$C_\beta$ ($\delta_H$ = 2.87, 3.07 ppm to $\delta_C$ = 34.9 ppm) and Ala8-$H_\beta$/Ala4-$C_\beta$ ($\delta_H$ = 2.83, 2.92 ppm to $\delta_C$ = 33.8 ppm), as well as Ala6-$H_\beta$/Ala9-$C_\beta$ ($\delta_H$ = 2.80, 2.87 ppm to $\delta_C$ = 33.1 ppm) and Ala9-$H_\beta$/Ala6-$C_\beta$ ($\delta_H$ = 2.98 ppm to $\delta_C$ = 35.4 ppm), suggested two lanthionine rings in Ser4-Cys8 and Ser6-Cys9. Two free carboxylic groups ($\delta_C$= 173.0 ppm for Met7 and $\delta_C$= 171.2 ppm for Ala9), along with one methylation ($\delta_H$= 2.54 ppm, $\delta_C$= 32.7 ppm for $CH_3$) on Gly1 and one acetylation ($\delta_C$= 169.3 ppm for CO and $\delta_H$= 1.90 ppm, $\delta_C$= 22.7 ppm for $CH_3$) on Cys8, suggesting its classification of DADC lanthipeptides. Combining HSQC, COSY, and HMBC data, compound **20** was thereby deduced to contain 1 × *N-Me*-Gly, 1 × Leu, 1 × Pro, 1 × Ala, 1 × Met, 1 ×Lan and 1 ×*N-Ac*-Lan. The stereochemistry of **20** was confirmed with L configuration in all unmodified amino acids by advanced Marfey's analysis (Supplementary Table 4). The remaining two Lan residues are LL (2*R*,6*R*) configurations compared with the standard samples (Supplementary Fig. 9 and Supplementary Table 4).

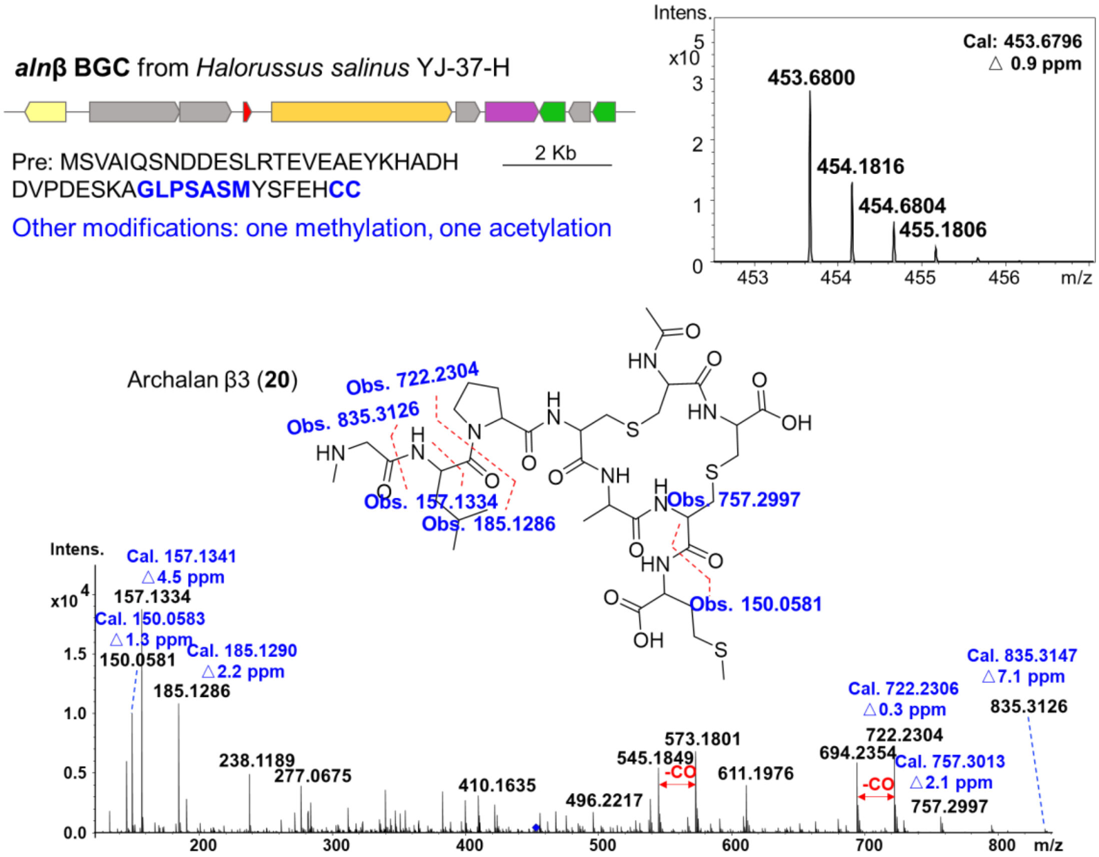

#### Supplementary Figure 32. LC-MS/MS analysis of archalan β3 (20)

The MS1 data of archalan β3 (**20**), m/z 453.6800 $[M+2H]^{2+}$, couldn't match the core peptide as classical lanthipeptide indicating its distinct features. The MS/MS data provided evidence of a Met residue at the C-terminus (m/z 150.0581 $[M+H]^{+}$) and methylated 'GL' fragments at the N-terminus (m/z 185.1286 $[M+H]^{+}$), which were aligned with partial fragments of the core peptide (<u>GL</u>PSAS<u>M</u>YSFEHCC). To investigate the complex core peptide region and associated modifications, **20** was isolated and characterized using NMR spectroscopy. Ultimately, the characterized structure of **20** exhibited the same feature as DADC lanthipeptides, where partial unmodified amino acids (YSFEH) within the thioether rings were cleaved to expose the diamino and dicarboxylic groups. In addition, one methylation and one acetylation were located on the first and the last amino group respectively, supporting the engineering possibility in these four termini.

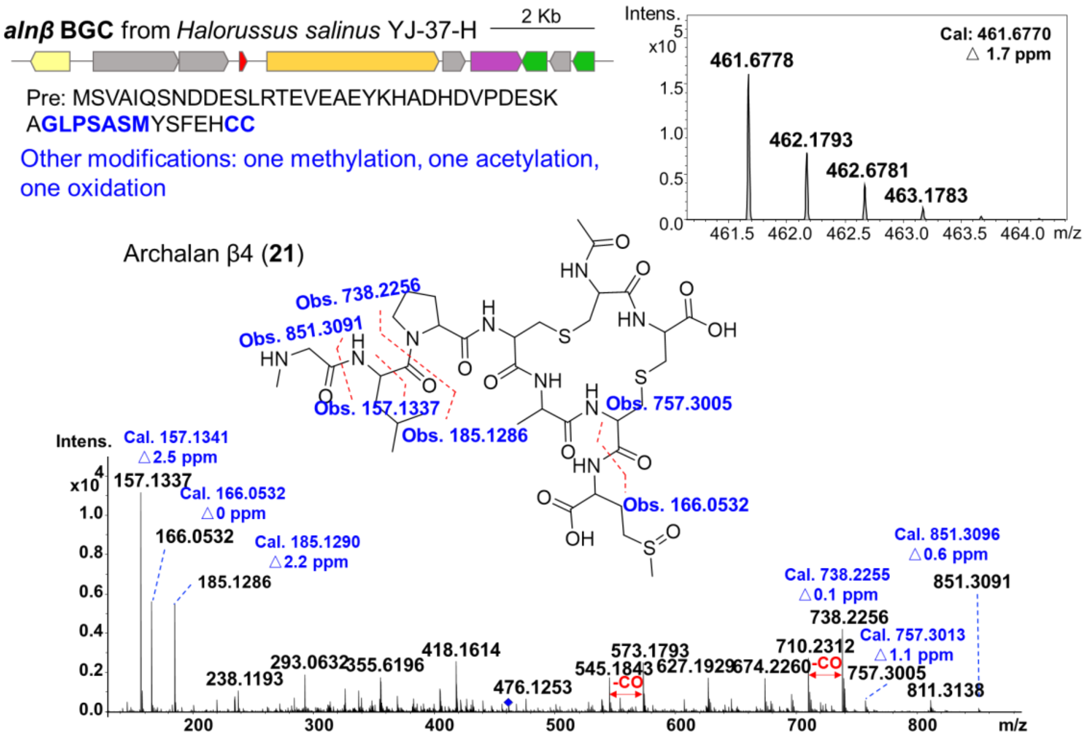

#### Supplementary Figure 33. LC-MS/MS analysis of archalan β4 (21)

Based on the confirmed structure of archalan β3 (**20**, m/z 453.6800 $[M+2H]^{2+}$), archalan β4 (**21**, m/z 461.6778 $[M+2H]^{2+}$) was proposed with one more oxidation (O, 15.9949 Da) deduced from the MS1 mass shift (15.5596 Da). Furthermore, in the MS/MS analysis, no disparities were observed at the N-terminus. However, the presence of the fragment m/z 166.0532 $[M+H]^+$ in **21** instead of m/z 150.0581 $[M+H]^+$ in **20** supported the occurrence of a methionine sulfoxide at the first C-terminal Met residue. Therefore, the structure of archalan β4 (**21**) was proposed as an analog of the validated archalan β3 (**20**), featuring an additional oxidation on the Met residue.

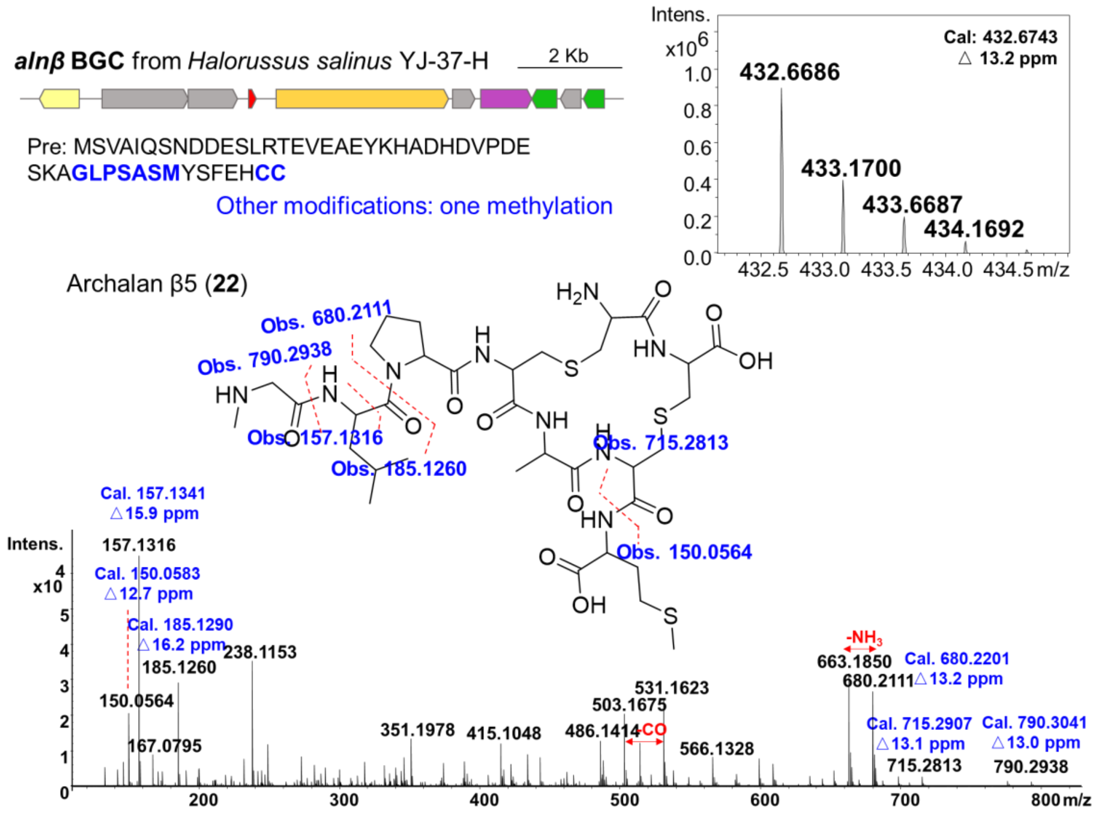

#### Supplementary Figure 34. LC-MS/MS analysis of archalan β5 (22)

In the MS1 profile, a reduction of 42 Da was observed in archalan β5 (**22**, m/z 432.6686 $[M+2H]^{2+}$) when compared with archalan β3 (**20**, m/z 453.6800 $[M+2H]^{2+}$), implying the potential removal of an acetylation residue (42.0106 Da). Furthermore, the MS/MS data confirmed that the first N-terminus and C-terminus of **22** were consistent with those of **20**, indicating the preservation of N-methylation. Distinct variations were observed in the fragments containing the terminal Cys residues, where the acetylation was located in **20**, implying no acetylation in **22**. Therefore, the structure of archalan β5 (**22**) was proposed without the acetylation at the second N-terminus based on its MS/MS analysis, the biosynthetic rule and the confirmed structure of archalan β3 (**20**). The structure of archalan β3 and β5, with or without the acetylation, suggested the involvement of an acetylation enzyme in catalyzing the transformation from archalan β5 to archalan β3, following hydrolysis and transport processes.

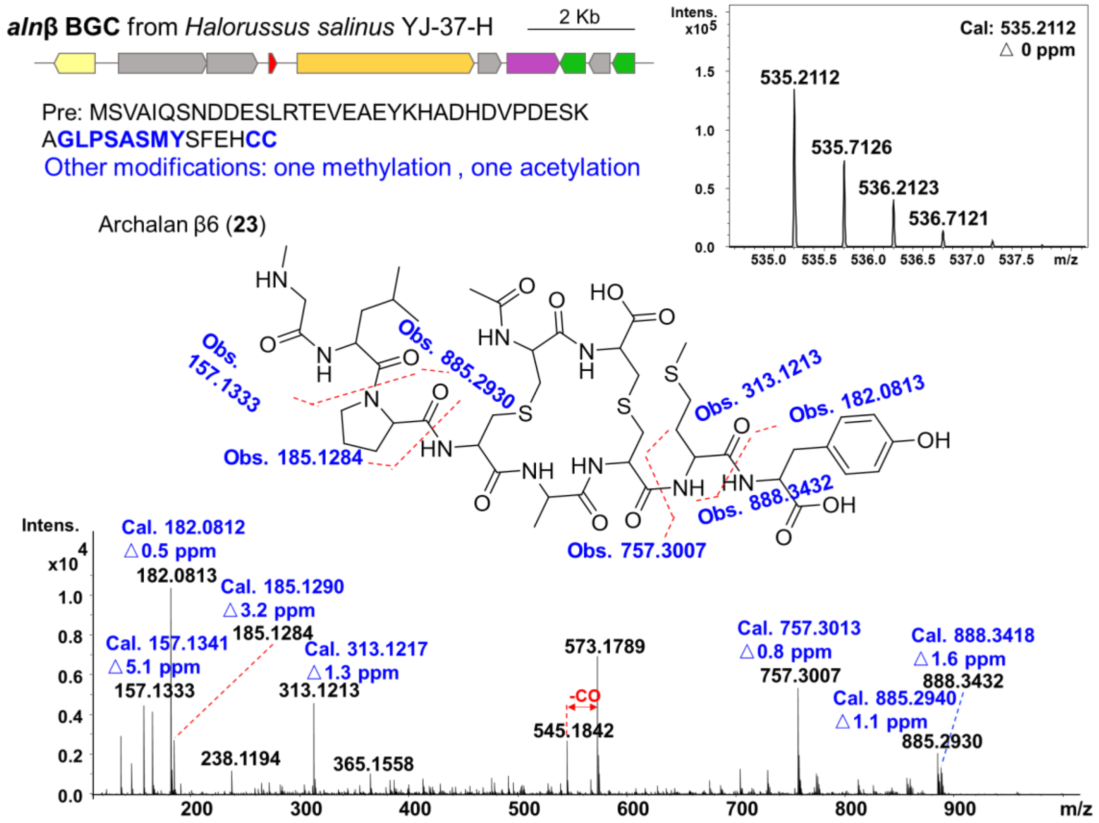

#### Supplementary Figure 35. LC-MS/MS analysis of archalan β6 (23)

The observed mass shift of 163.0624 Da between archalan β6 (**23**, m/z 535.2112 $[M+2H]^{2+}$) and archalan β3 (**20**, m/z 453.6800 $[M+2H]^{2+}$) is proposed to be a Tyr residue (163.0633 Da). The MS/MS data substantiated that the N-terminal fragments of archalan β6 (**23**) were identical to those of β3 (**20**), whereas the C-terminus exhibited differences characterized by mass shifts. Key fragments at m/z 182.0813 $[M+H]^{+}$ and m/z 313.1213 $[M+H]^{+}$ in **23** confirmed the presence of a Tyr following the Met residue in the first C-terminus, which is consistent with the precursor sequence. Varied amino acid extensions of archalan β6 and archalan β3 could potentially be attributed to extracellular proteases from the wild-type strain.

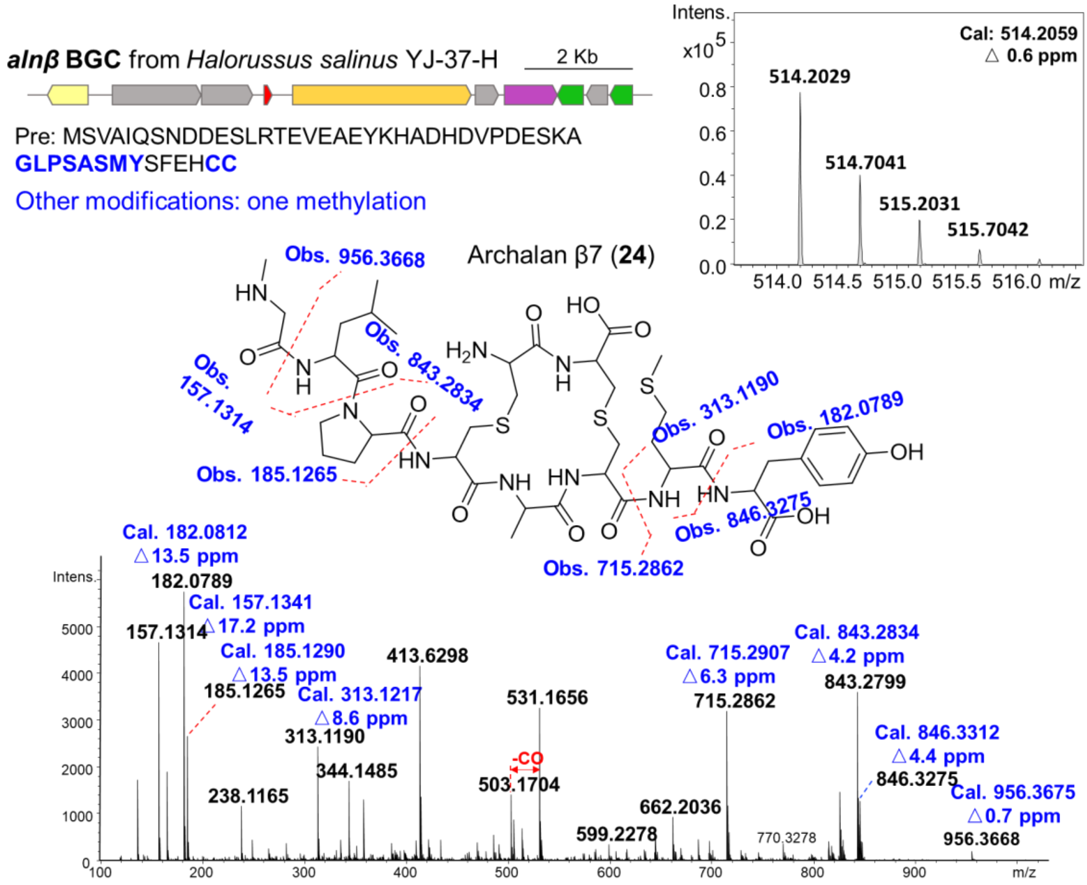

#### Supplementary Figure 36. LC-MS/MS analysis of archalan β7 (24)

The MS1 profile of archalan β7 (**24**, m/z 514.2029 $[M+2H]^{2+}$) with a decrease of 42 Da than archalan β6 (**23**, m/z 535.2112 $[M+2H]^{2+}$), implying the loss of acetylation residue (42.0106 Da). Analysis of the MS/MS fragmentation patterns confirmed that the initial N-terminal fragments of archalan β7 exhibited identical to those of archalan β6 and archalan β3. Moreover, akin to archalan β6, the C-terminal fragments of archalan β7 featured an additional Tyr residue following the Met residue, as evidenced by m/z 182.0789 $[M+H]^+$ and m/z 313.1190 $[M+H]^+$. Hence, the structure of archalan β7 was proposed without acetylation but with an additional Tyr in comparison to archalan β3, based on its MS/MS analysis and biosynthetic rule. These structures suggest that the putative acetylase may utilize archalan β7 as a substrate to produce archalan β6, a process probably following transport and leader peptide hydrolysis stages.

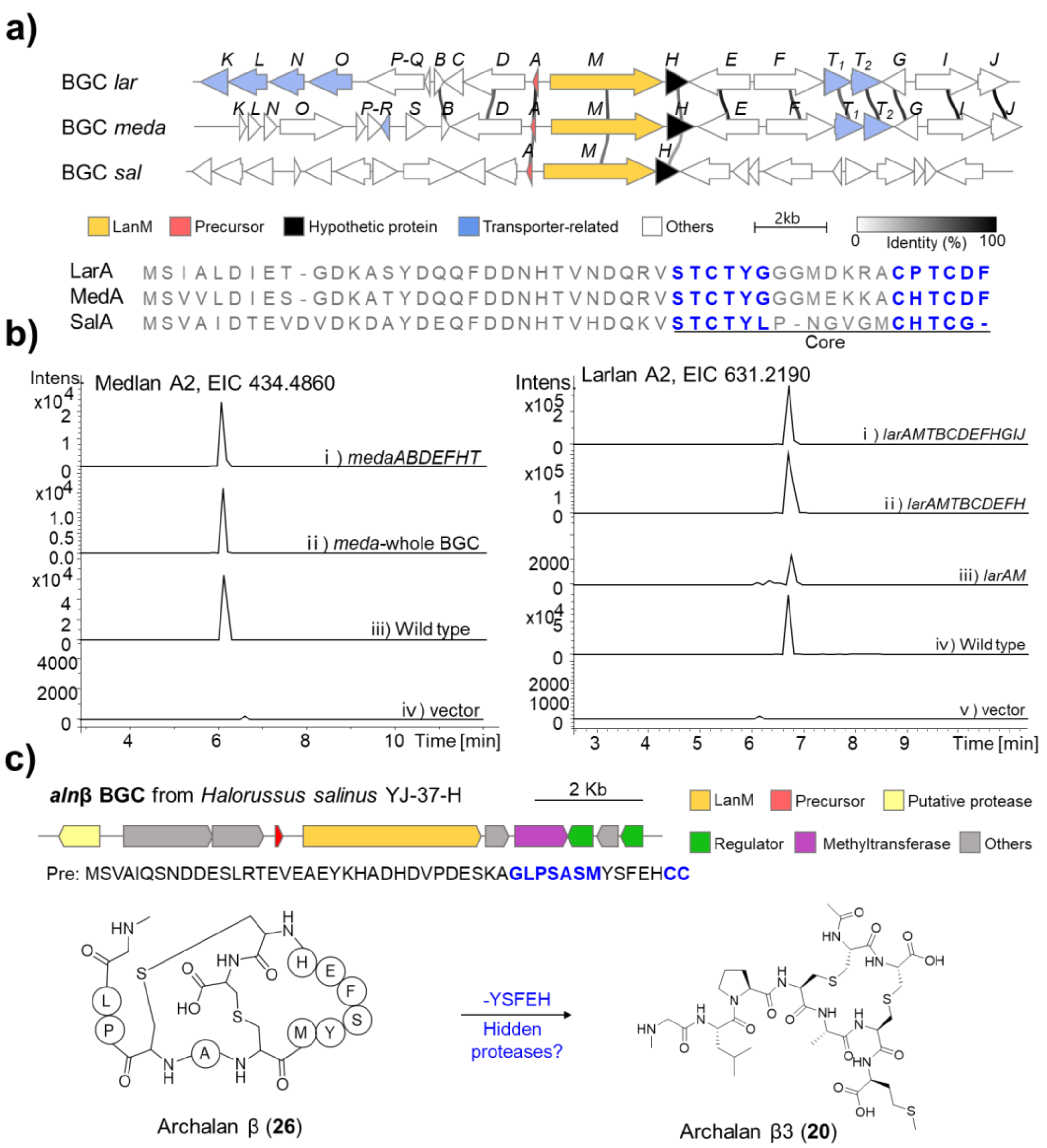

#### Supplementary Figure 37. Reconstitution of DADC lanthipeptide BGCs in *H. volcanii* H1424

a, Synteny analysis of BGCs from DacA family. BGCs *lar*, *meda* and *sal* are identified from strain *H. larsenii* JCM 13917, *H. mediterranei* ATCC35500, and *Halocatena salina* AD-1, respectively. The presence of three conserved genes: *A*, *M*, and *H*, might serve as the boundary of DADC lanthipeptide BGCs. Precursor peptide LanA, modification enzyme LanM, transporter, and hypothetic protein H are in red, orange, blue, and black respectively. Lines link the identity threshold of genes higher than 30%. b, LC-MS analysis of in vivo heterologous expression of BGCs *meda* and *lar* with different gene combinations. In vivo reconstitution study of *lar* BGC with only *larA* and *larM* resulted in the production of a trace amount of larlan A2, suggesting the possibility of putative hidden peptidases from the host genome. c. The construction of BGC *alnβ* and its classical and non-canonical products. Heterologous expression of the complete BGC yielded solely the classical lanthipeptide, lacking the DADC lanthipeptide, whereas the wild-type strain produced both types of lanthipeptides. This outcome provides evidence for the involvement of putative proteases outside the BGC within the genome of the wild-type strain.

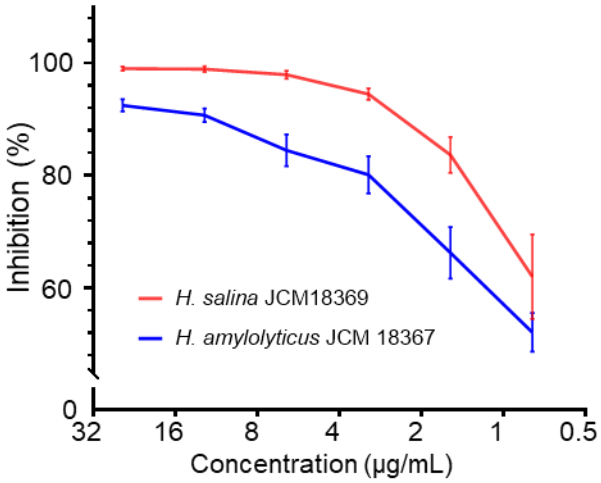

**Supplementary Figure 38. The antagonistic activity of archalan α (25) against *H. amylolyticus* JCM 18367 and *H. salina* JCM 18369.**

Bars show the mean ± standard error of the mean (SEM), n = 6. Each group has three biological replicates with two technical repeats each time. Archalan α exhibited a strong inhibition against *H. amylolyticus* JCM 18367 and *H. salina* JCM 18369 even at the concentration of 0.78 μg $mL^{-1}$.

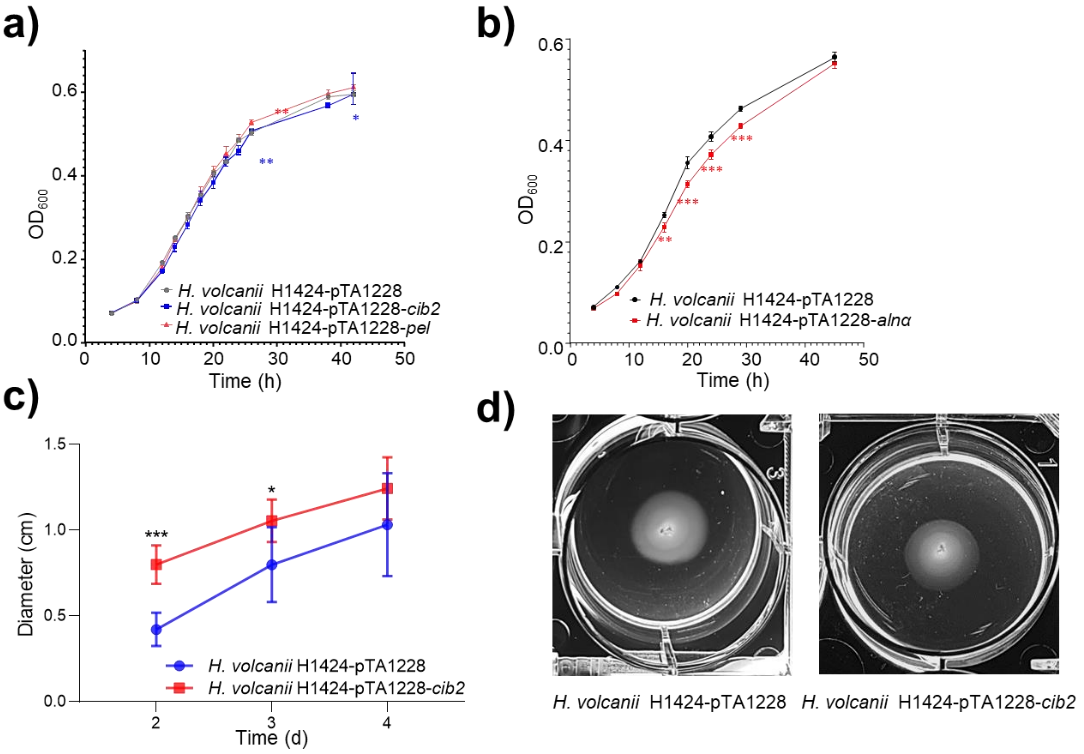

#### Supplementary Figure 39. Motility regulation assay of recombinant strains

a-b, The growth curve of recombinant strains with empty plasmid pTA1228, BGCs *cib2*, *pel* and *alnα* respectively. Each group has three biological replicates. The error bar represents mean ± SD, n = 3. *** $P < 0.001$, ** $P < 0.01$, * $P < 0.05$. c, Time-course motility assays of recombinant strains with empty plasmid pTA1228 or BGC *cib2.* Each group has three biological replicates with three technical repeats each time. The error bar represents mean ± SD, n=6. *** $P < 0.001$, * $P < 0.05$. d, The Bull's eye pattern of recombinant strains with empty plasmid pTA1228 or BGC *cib2* in 6 well plates.

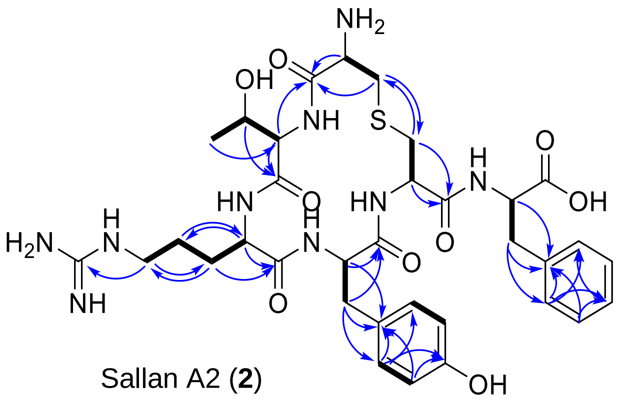

Sallan A2 (**2**)

**a) $^{1}$H NMR spectrum (600 MHz, Methonal-$d_4$) of sallan A2 (2)**

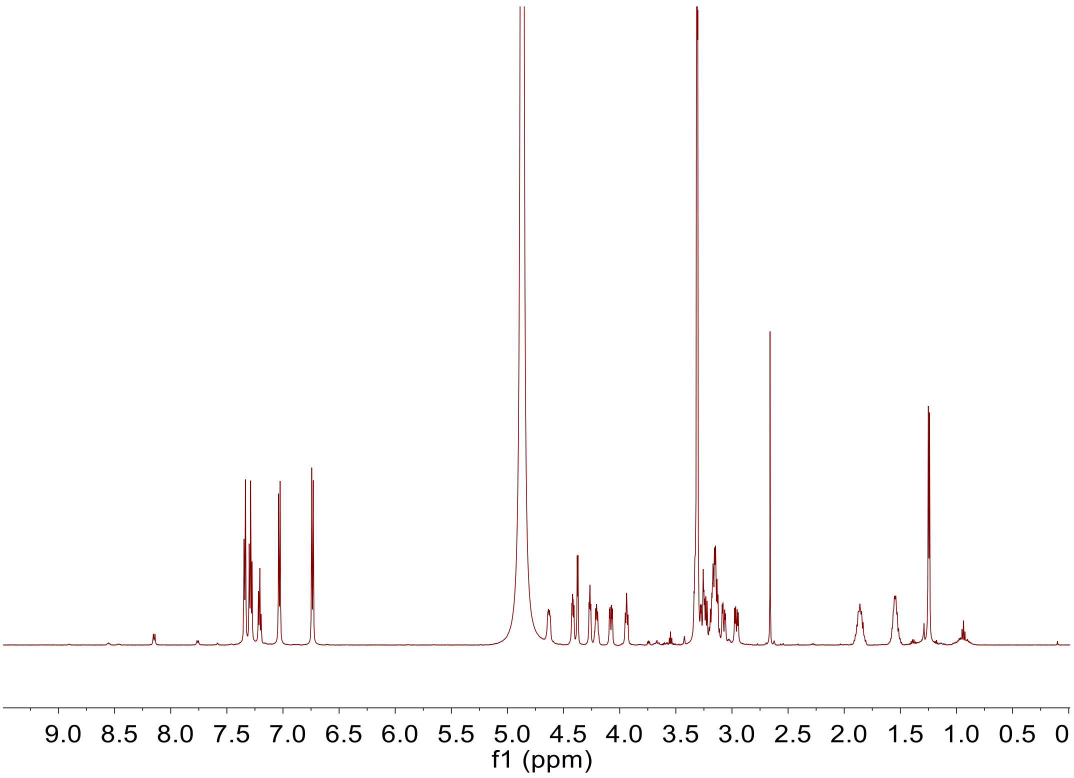

b) $^{13}$C NMR spectrum (150 MHz, Methonal-$d_4$) of sallan A2 (2)

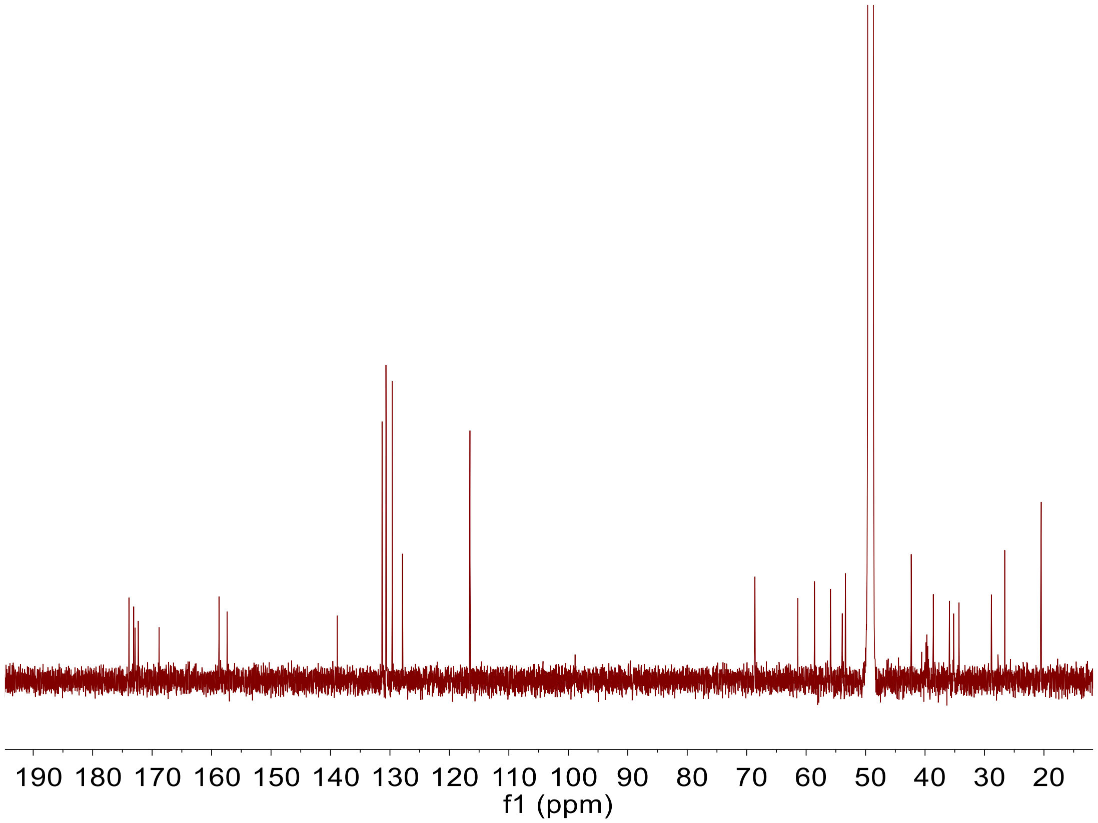

c) ¹H-¹H COSY spectrum (600 MHz, Methonal-$d_4$) of sallan A2 (2)

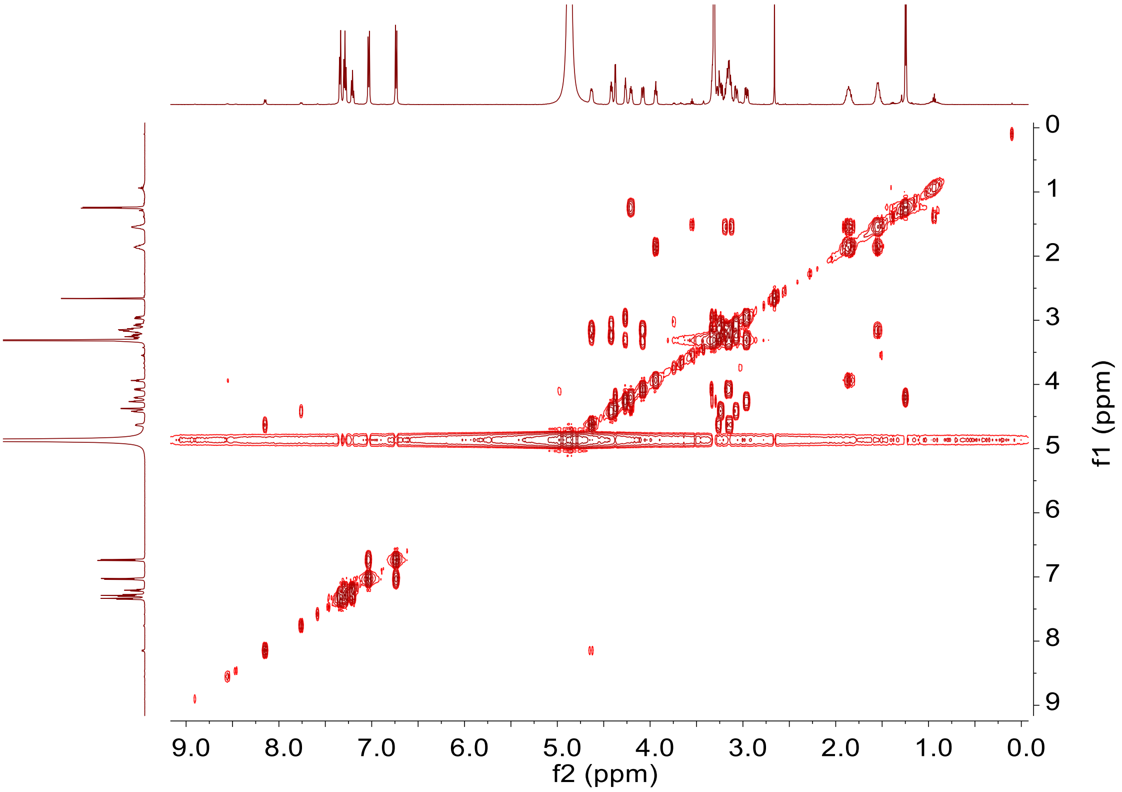

**d) HSQC spectrum (600 MHz, Methonal-$d_4$) of sallan A2 (2)**

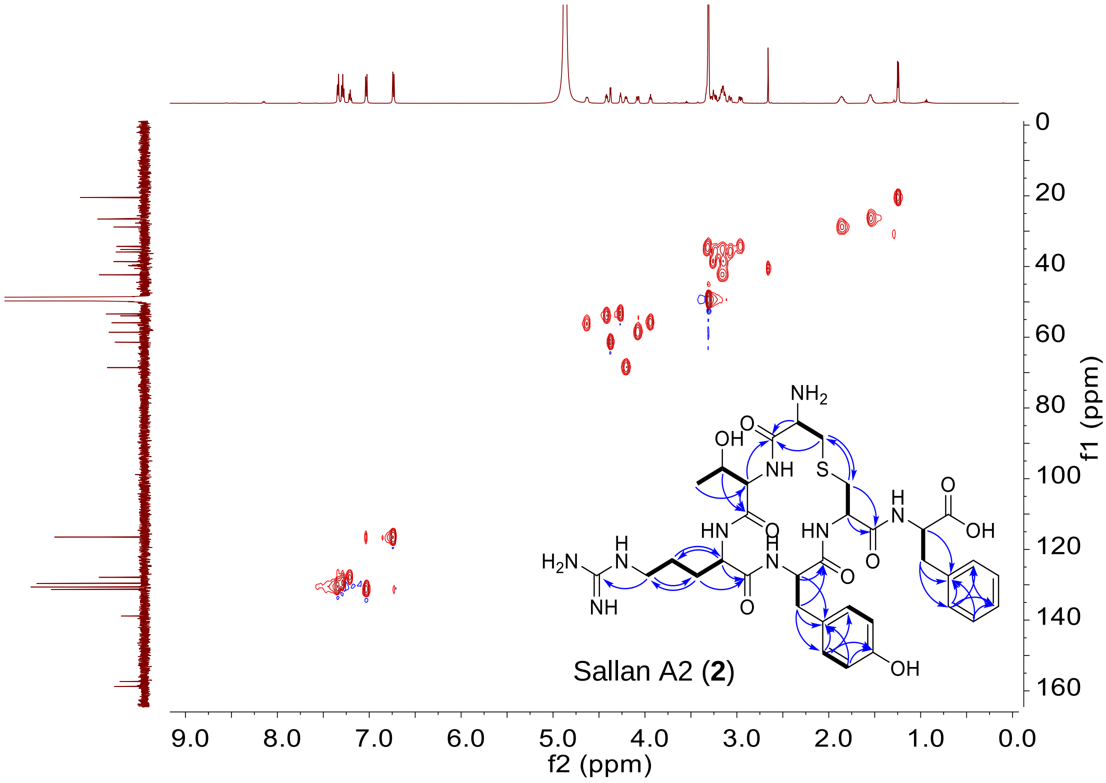

**e) HMBC spectrum (600 MHz, Methonal-$d_4$) of sallan A2 (2)**

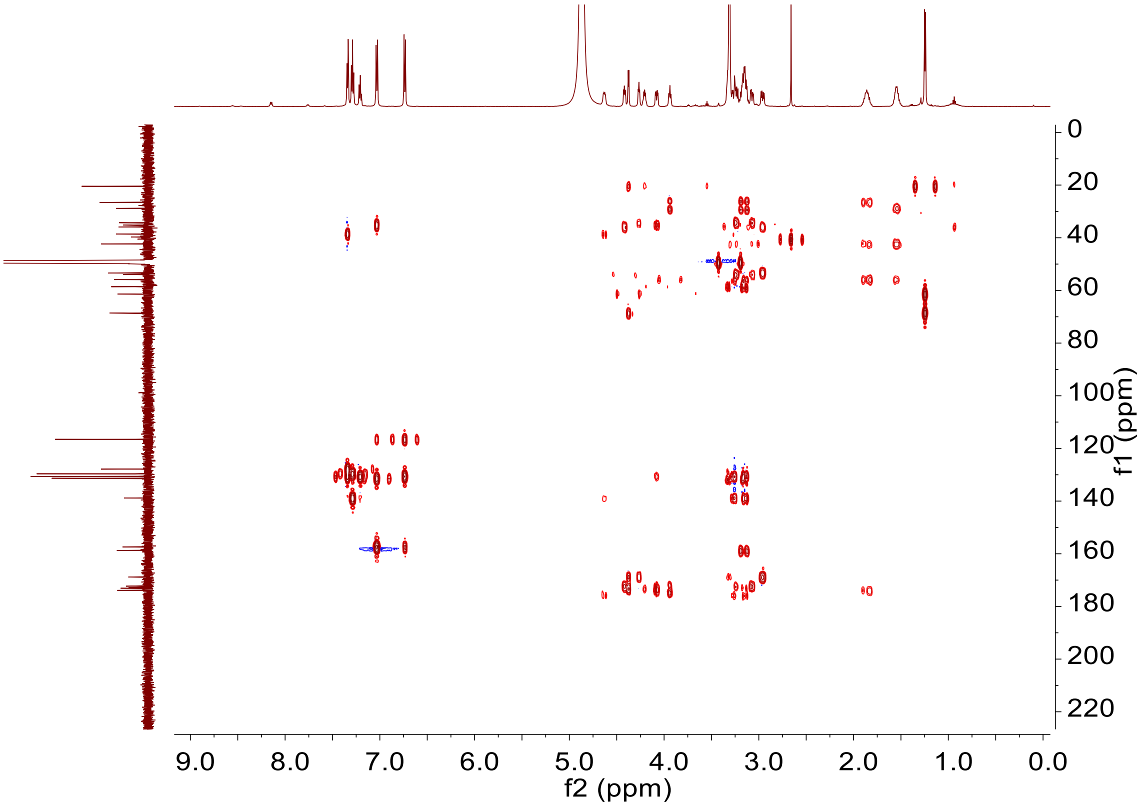

**f) NOESY spectrum (600 MHz, Methonal-$d_4$) of sallan A2 (2)**

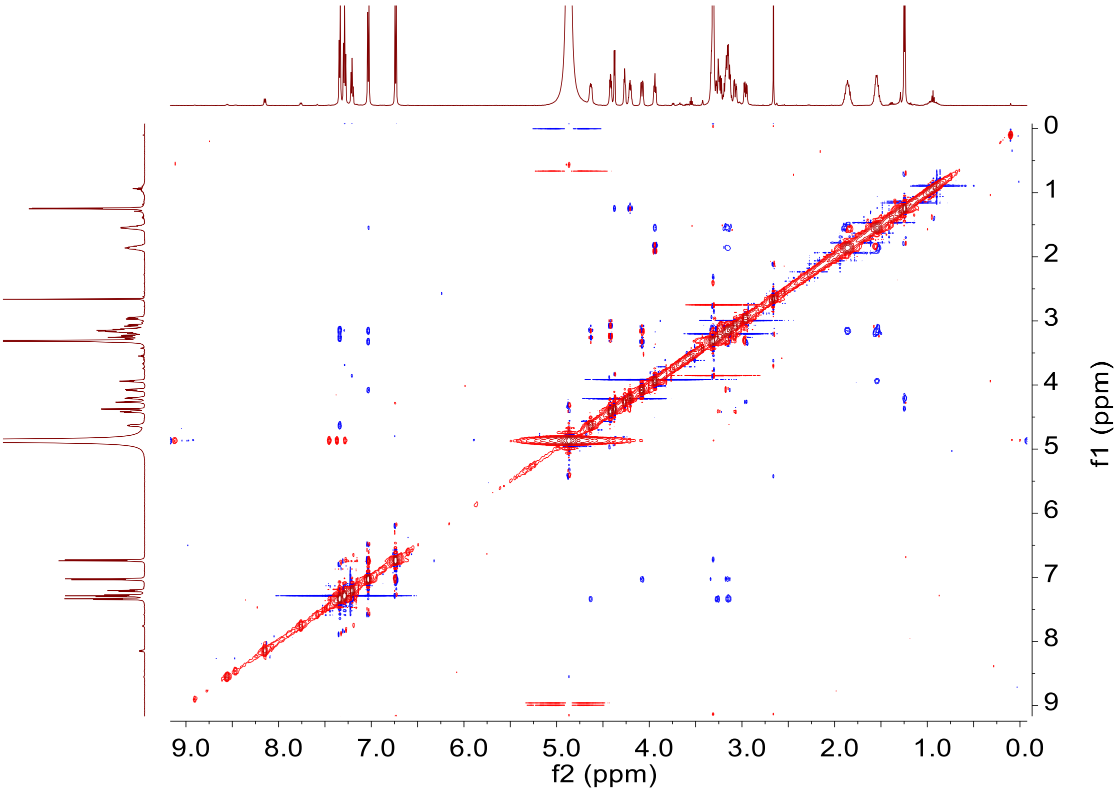

**Supplementary Figure 40. NMR spectra of sallan A2 (2) in Methonal-$d_4$: 600 MHz $^1$H NMR, 150 MHz $^{13}$C NMR, 600 MHz $^1$H-$^1$H COSY, HSQC, HMBC, NOESY spectra**

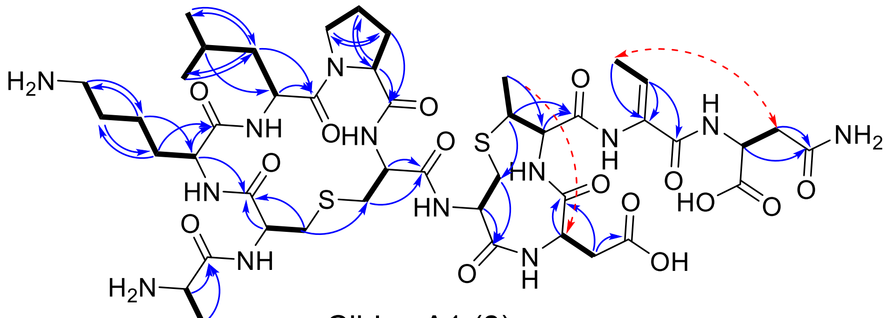

Ciblan A1 (**3**)

**a) [1]H NMR spectrum (600 MHz, Methonal-*d4*) of ciblan A1 (3)**

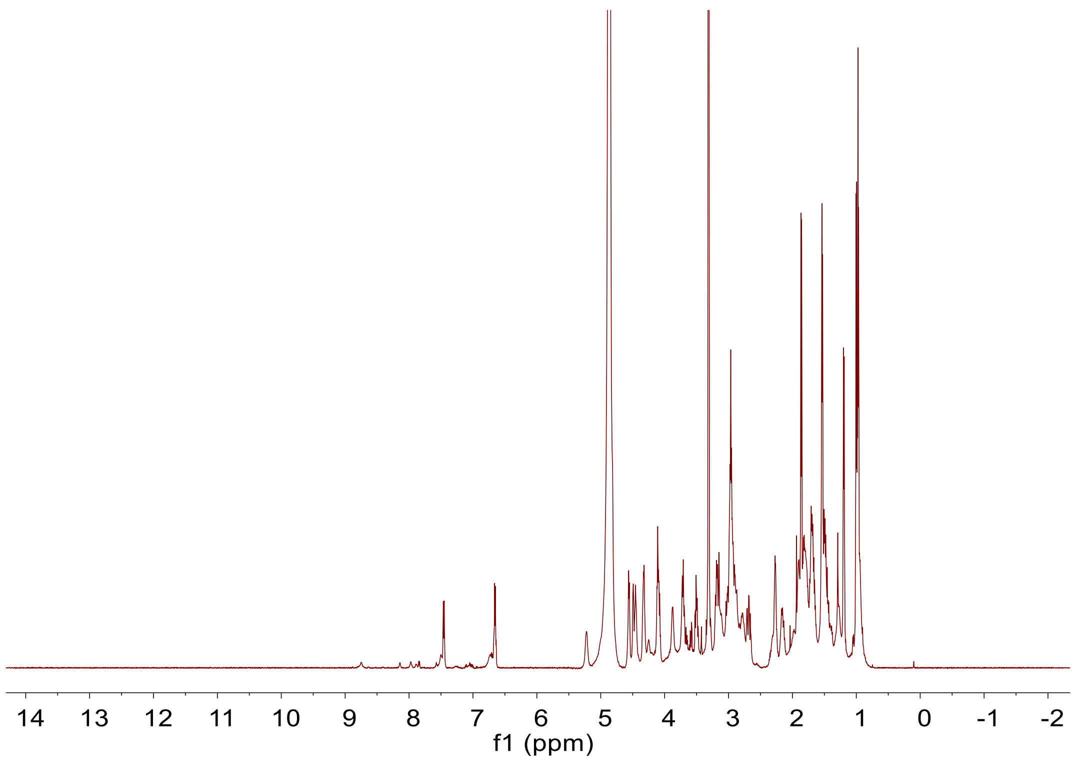

**b) $^{13}$C NMR spectrum (150 MHz, Methonal-$d_4$) of ciblan A1 (3)**

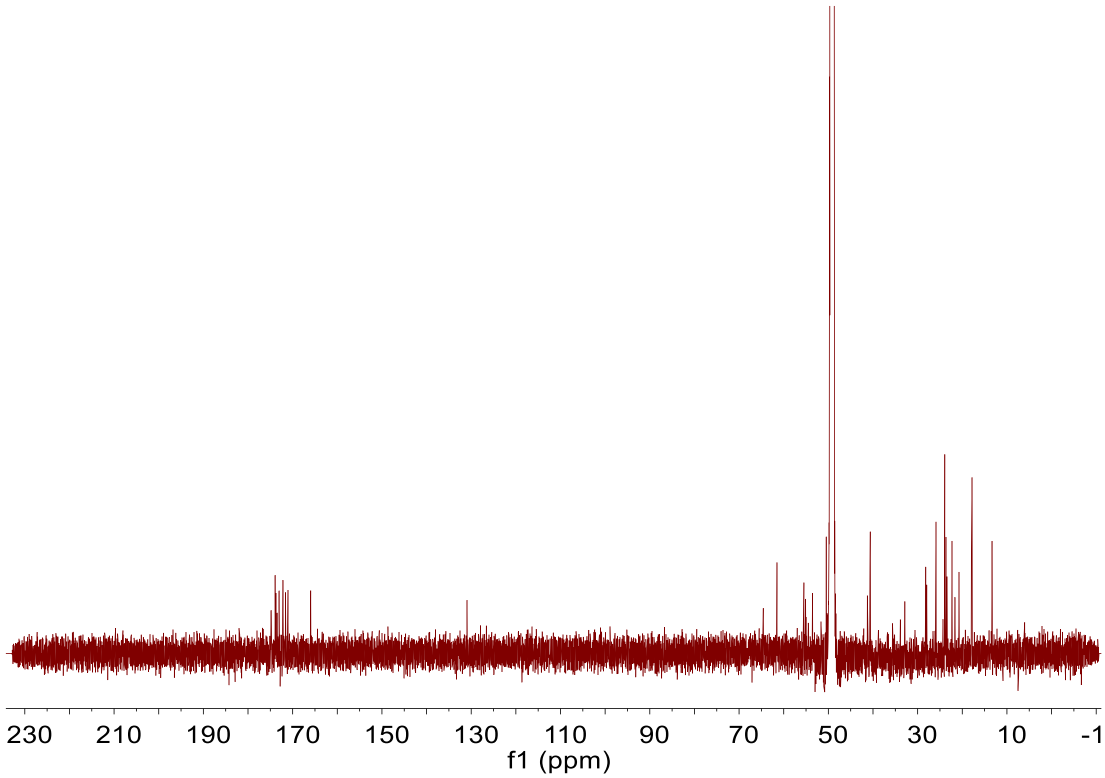

c) [1]H-[1]H COSY NMR spectrum (600 MHz, Methonal-$d_4$) of ciblan A1 (3)

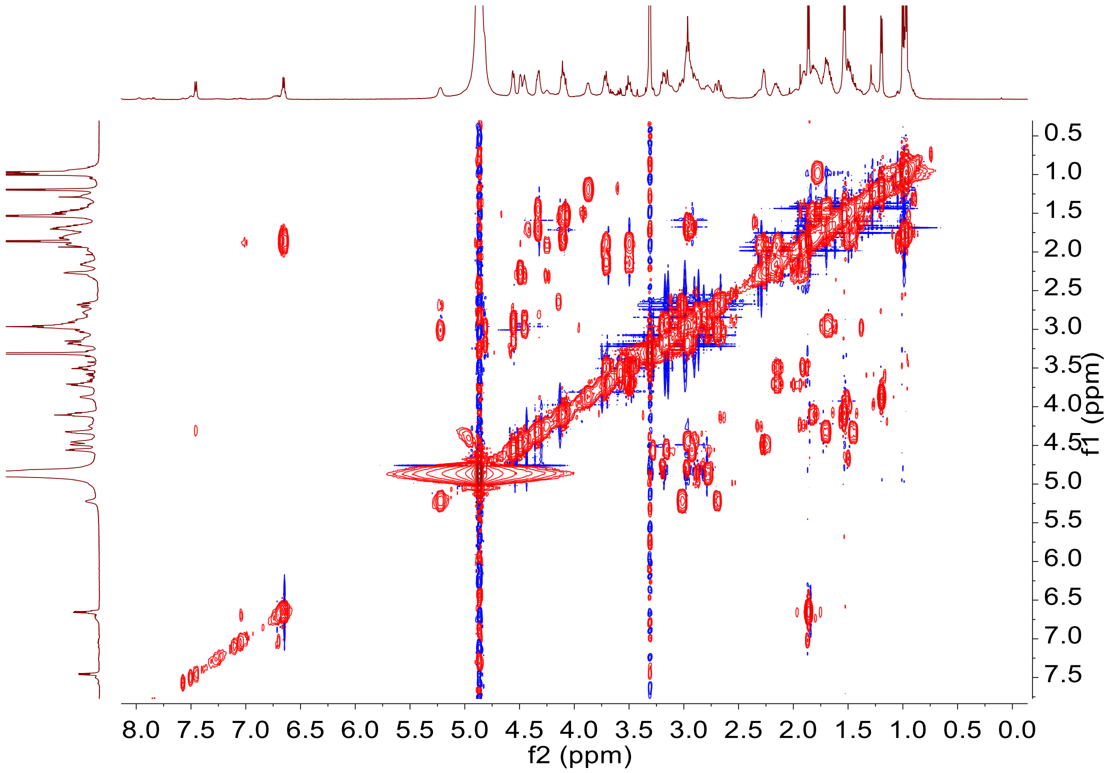

**d) HSQC NMR spectrum (600 MHz, Methonal-$d_4$) of ciblan A1 (3)**

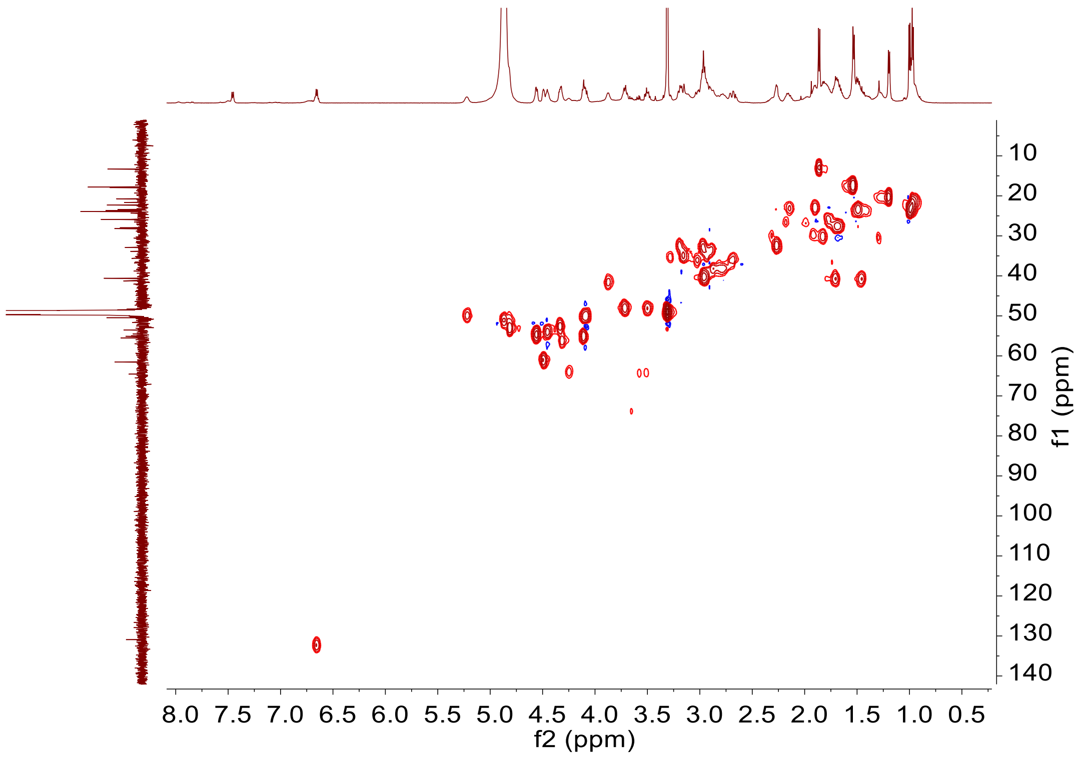

**e) HMBC NMR spectrum (600 MHz, Methonal-$d_4$) of ciblan A1 (3)**

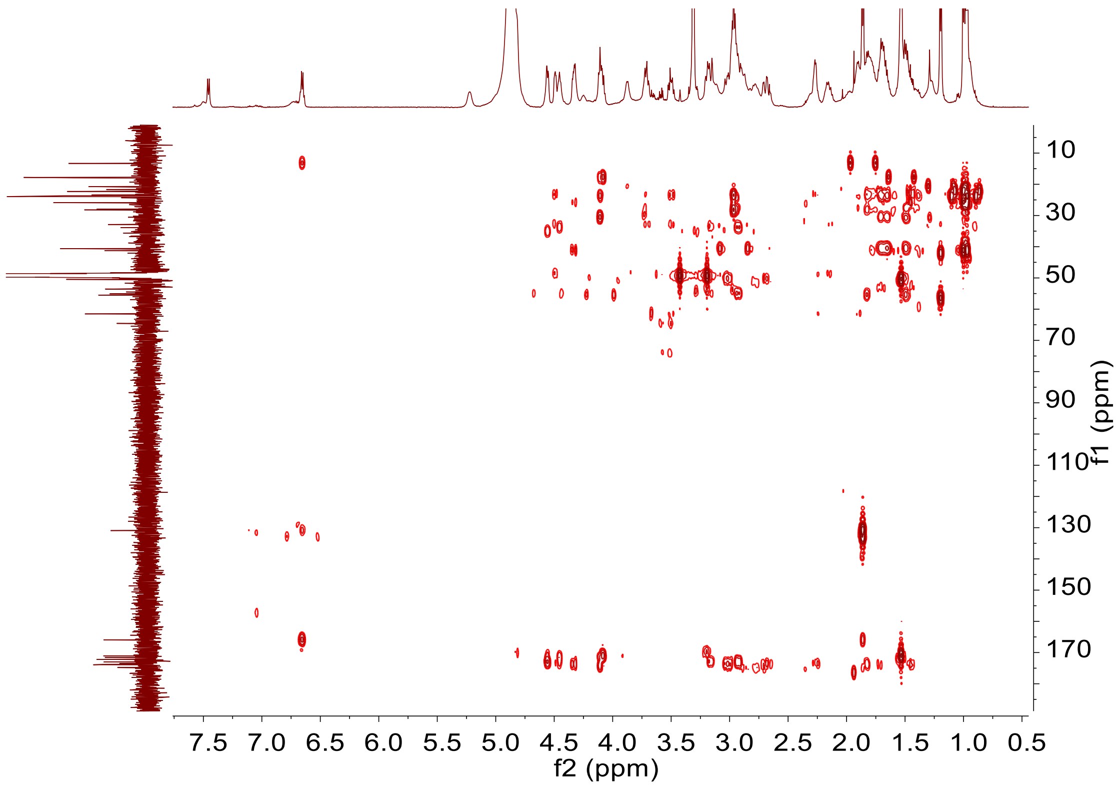

**f) TOCSY NMR spectrum (600 MHz, Methonal-$d_4$) of ciblan A1 (3)**

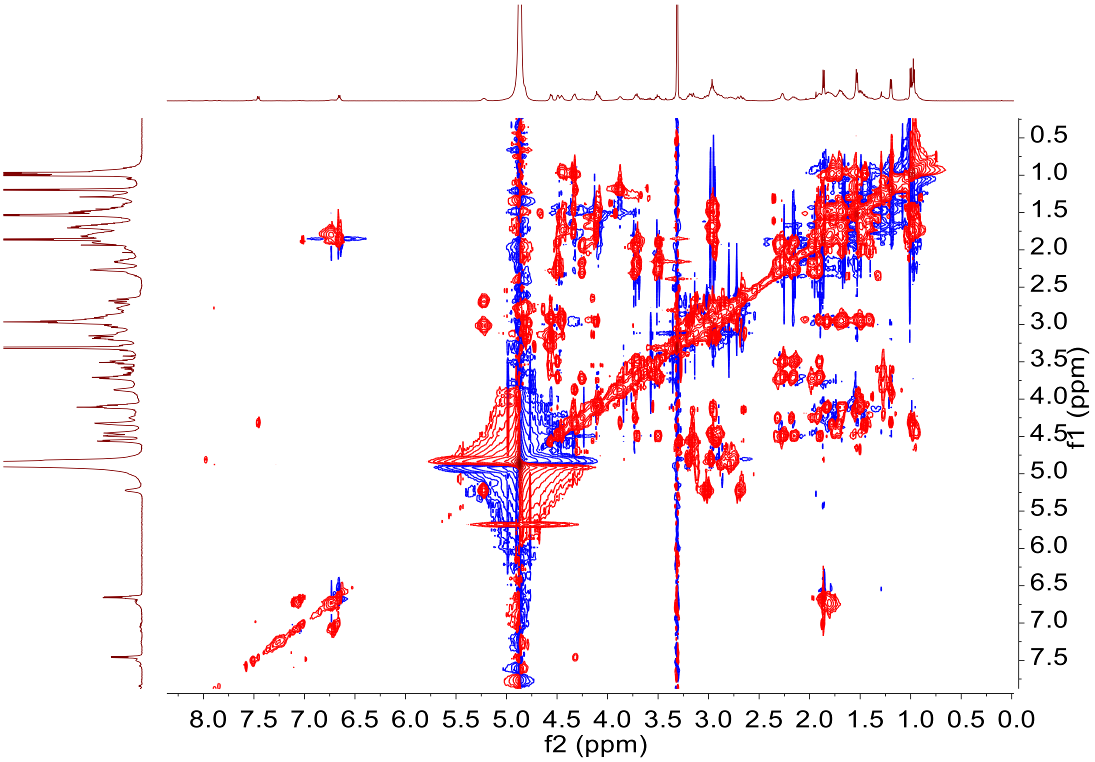

g) NOESY NMR spectrum (600 MHz, Methonal-$d_4$) of ciblan A1 (3)

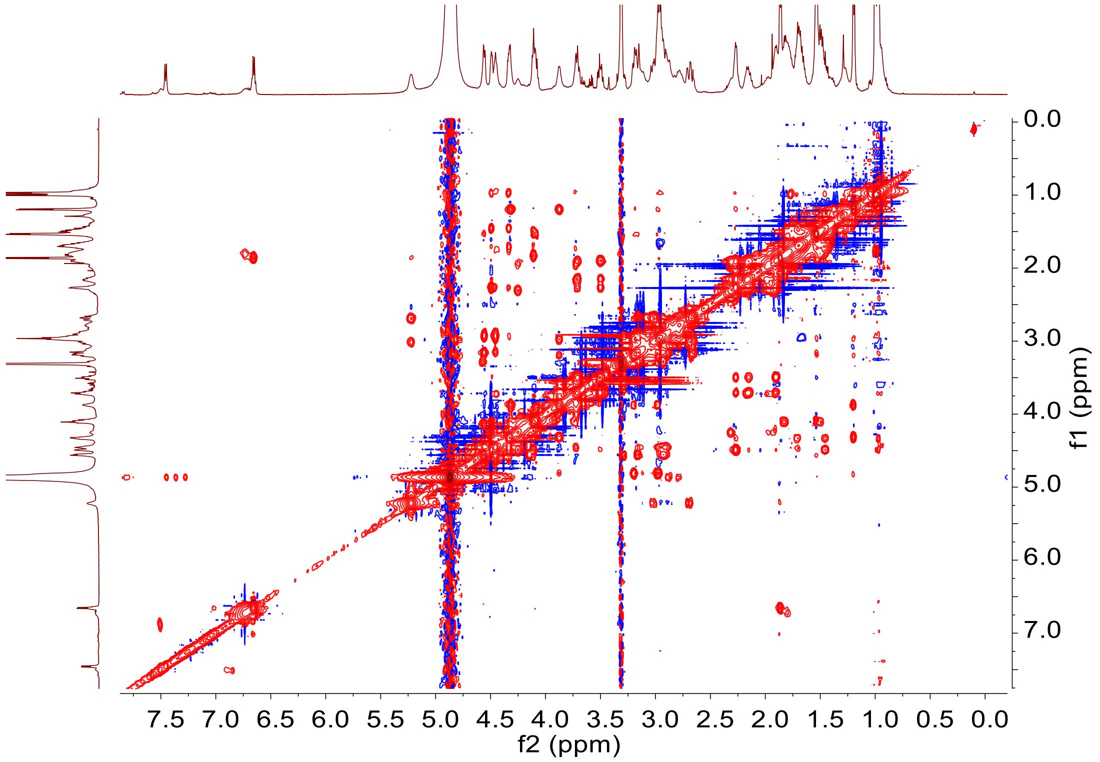

**Supplementary Figure 41. NMR spectra of ciblan A1 (3) in Methonal-$d_4$: 600 MHz $^1$H NMR, 150 MHz $^{13}$C NMR, 600 MHz $^1$H-$^1$H COSY, HSQC, HMBC, TOCSY, NOESY spectra**

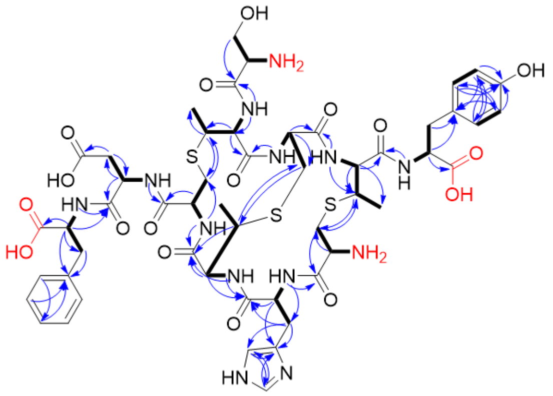

Medlan A1 (**13**)

**a) [1]H NMR spectrum (600 MHz, DMSO-*d*$_6$) of medlan A1 (13)**

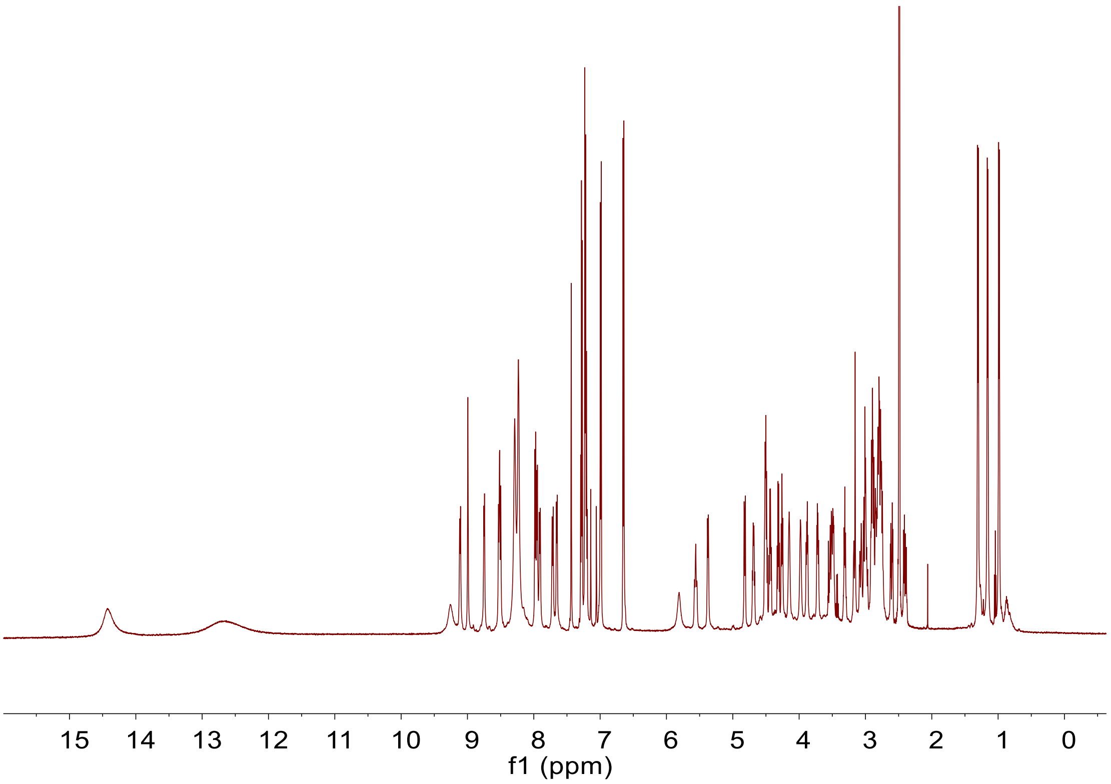

b) $^{13}$C NMR spectrum (150 MHz, DMSO-$d_6$) of medlan A1 (13)

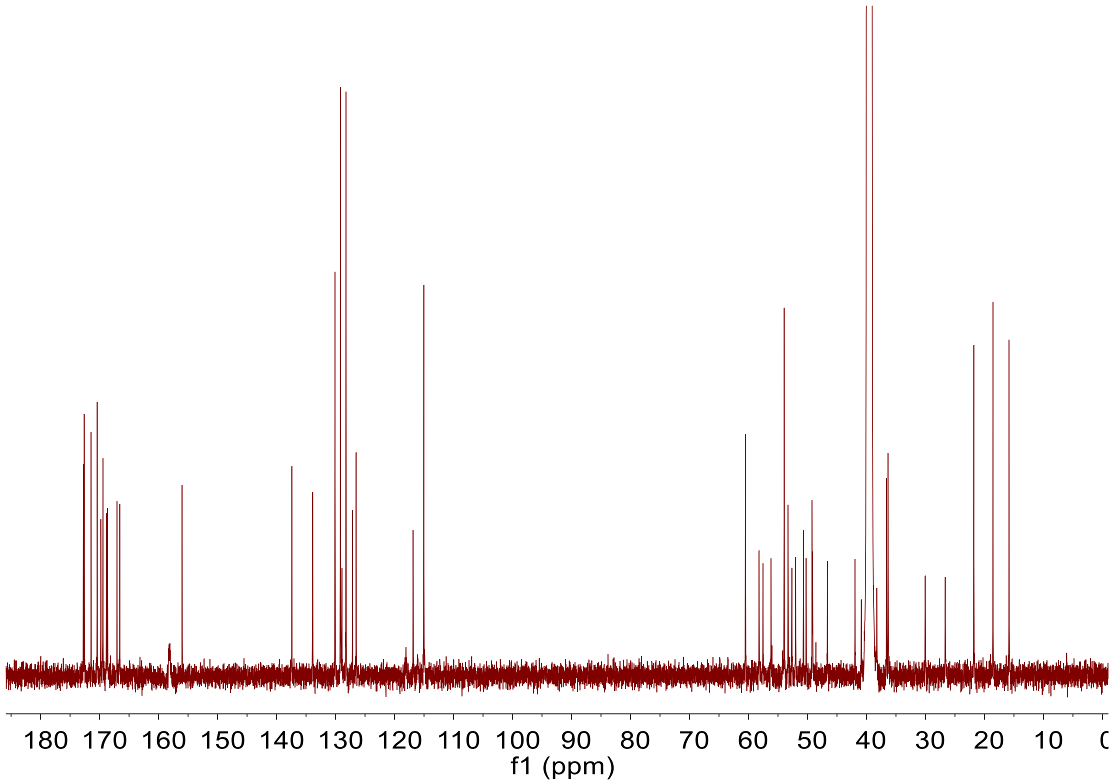

c) [1]H-[1]H COSY spectrum (600 MHz, DMSO-$d_6$) of medlan A1 (13)

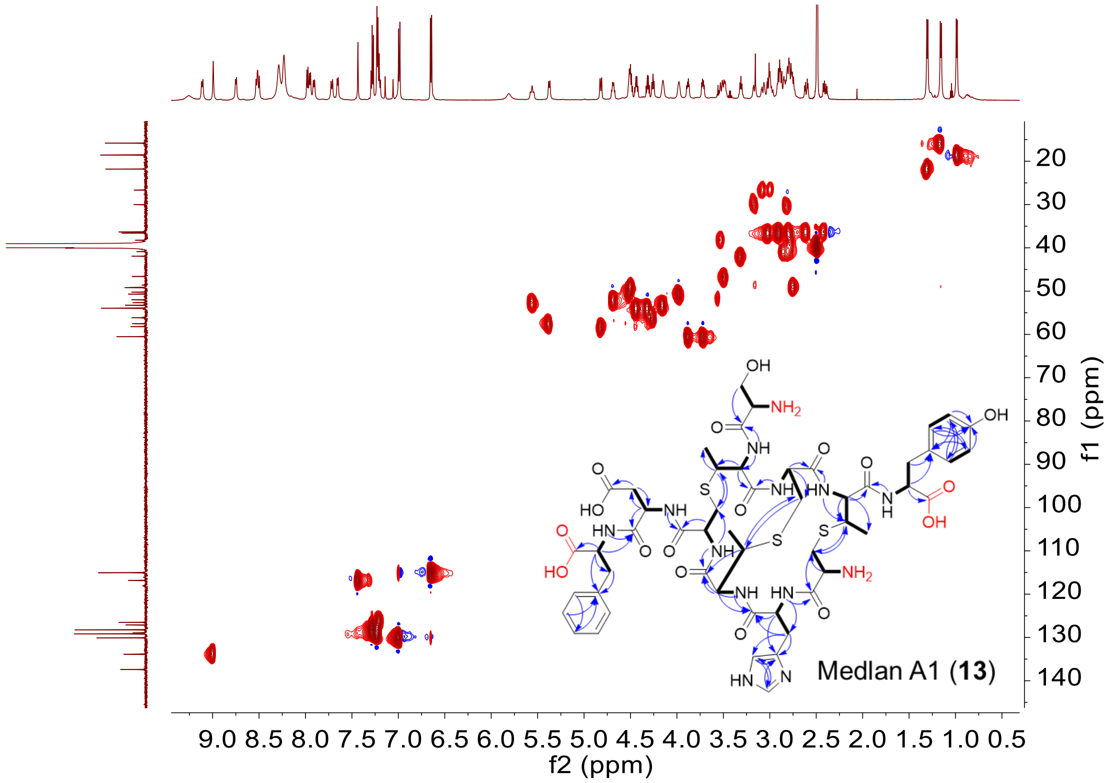

**d) HSQC spectrum (600 MHz, DMSO-$d_6$) of medlan A1 (13)**

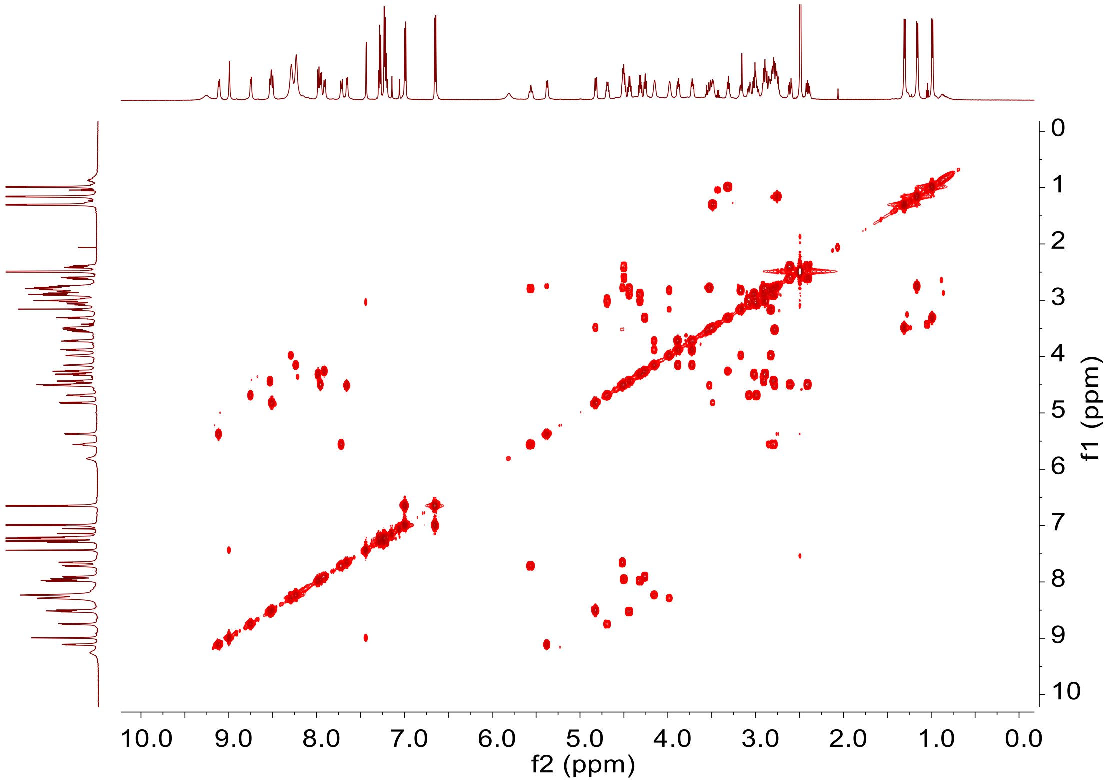

**e) HMBC spectrum (600 MHz, DMSO-$d_6$) of medlan A1 (13)**

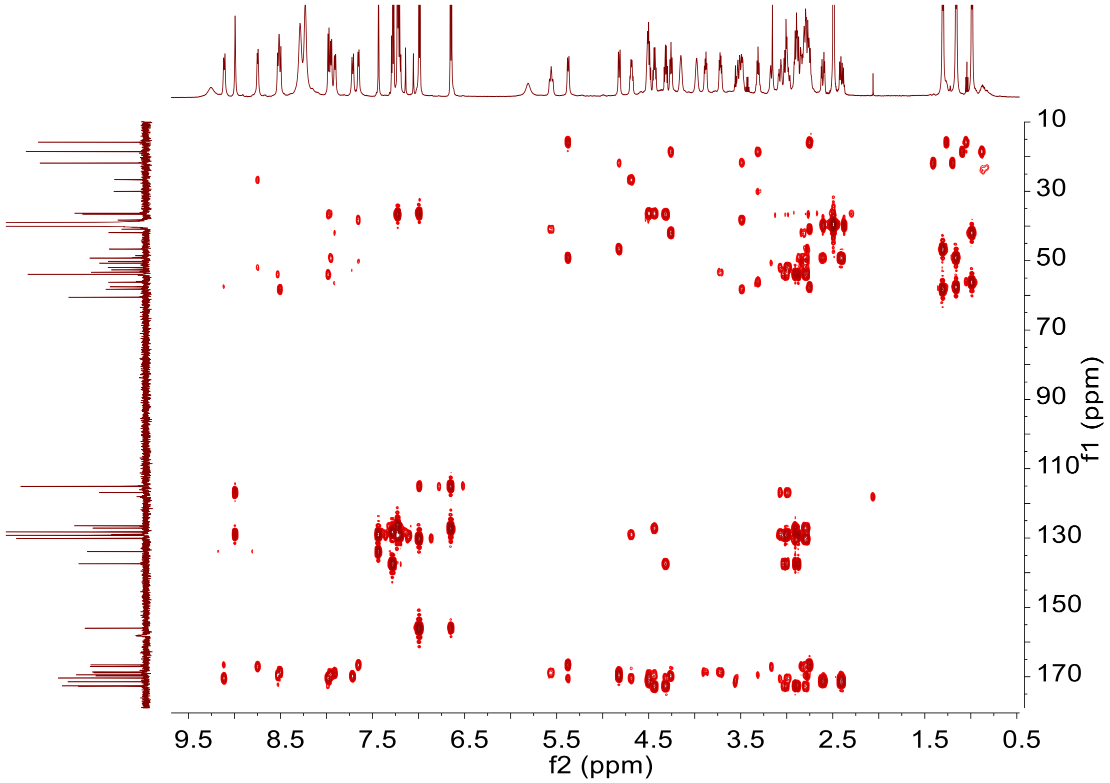

f) NOESY spectrum (600 MHz, DMSO-$d_6$) of medlan A1 (13)

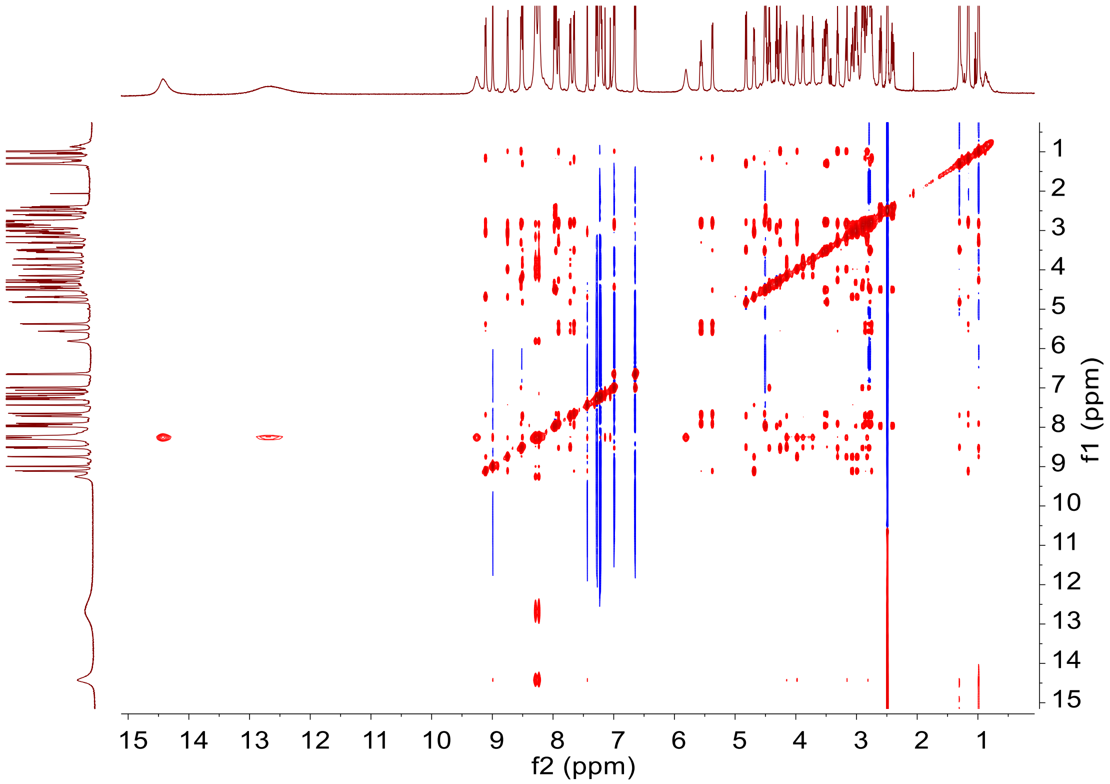

**Supplementary Figure 42. NMR spectra of medlan A1 (13) in DMSO-$d_6$: 600 MHz $^1$H NMR, 150 MHz $^{13}$C NMR, 600 MHz $^1$H-$^1$H COSY, HSQC, HMBC, NOESY spectra**

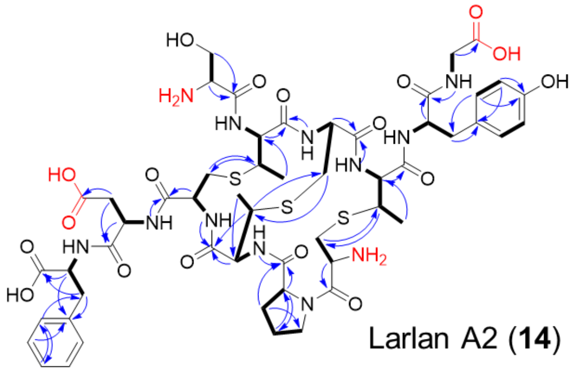

a) **[1]H NMR spectrum (600 MHz, DMSO-*d*$_6$) of larlan A2 (14)**

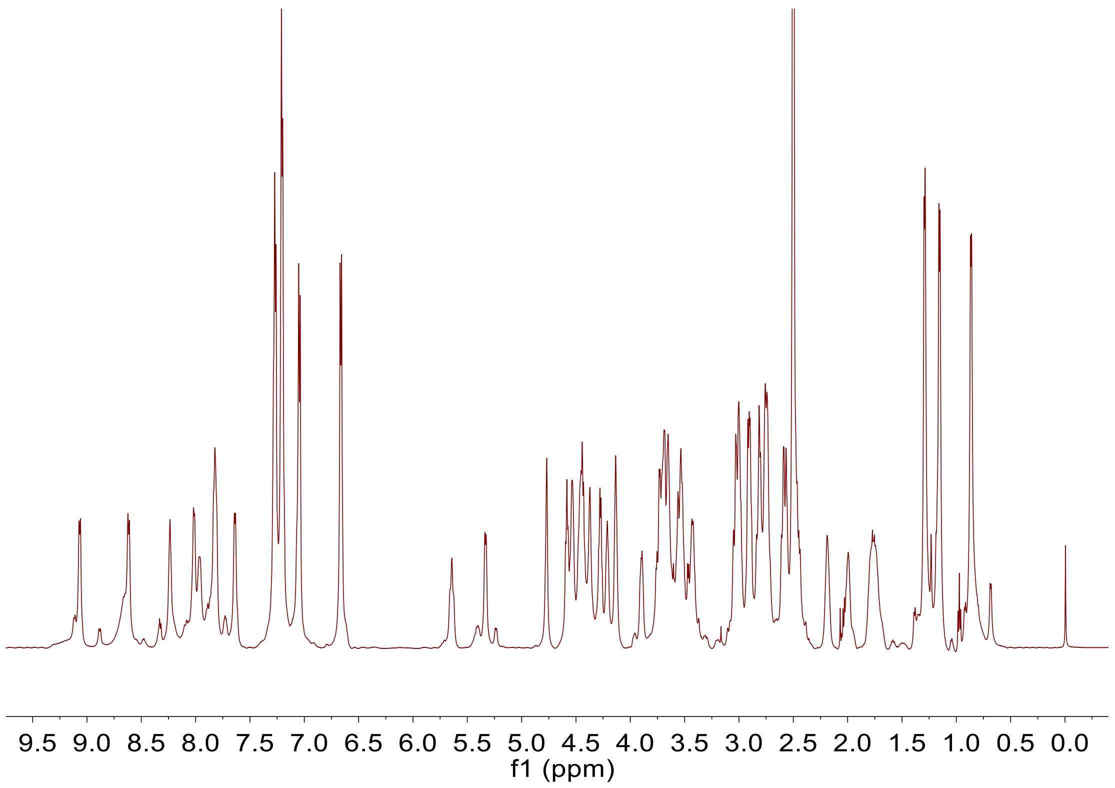

**b) $^{13}$C NMR spectrum (150 MHz, DMSO-$d_6$) of larlan A2 (14)**

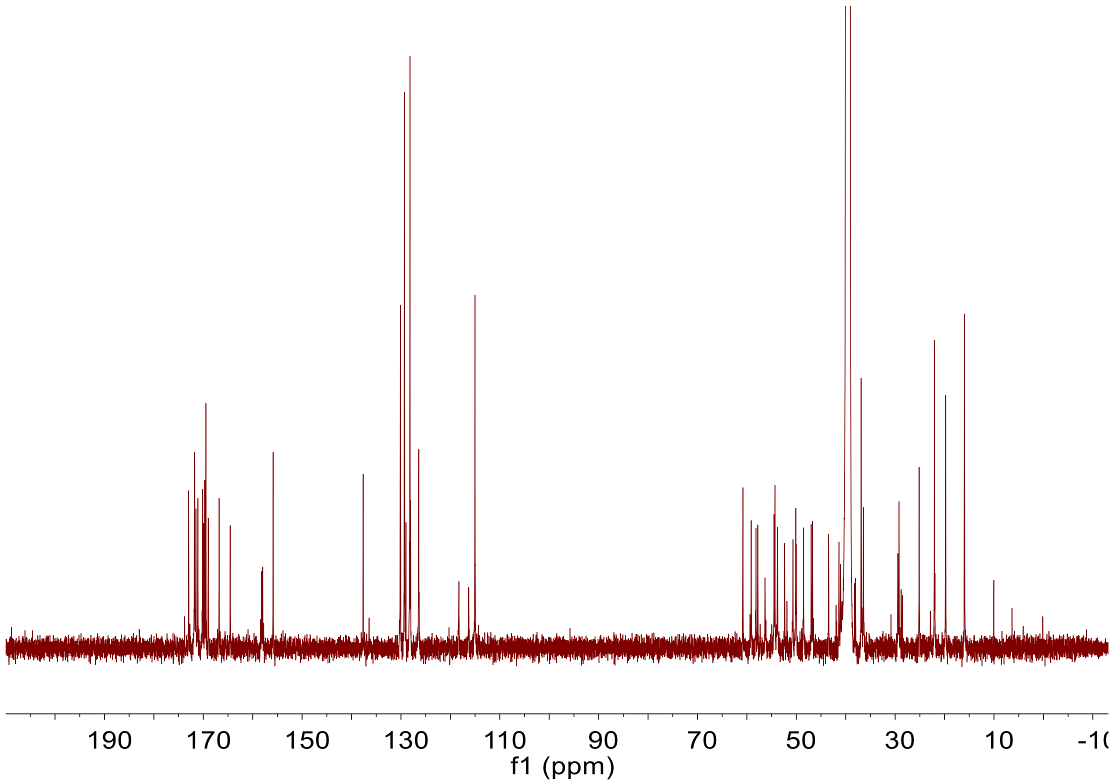

c) $^{1}H$-$^{1}H$ COSY spectrum (600 MHz, DMSO-$d_6$) of larlan A2 (14)

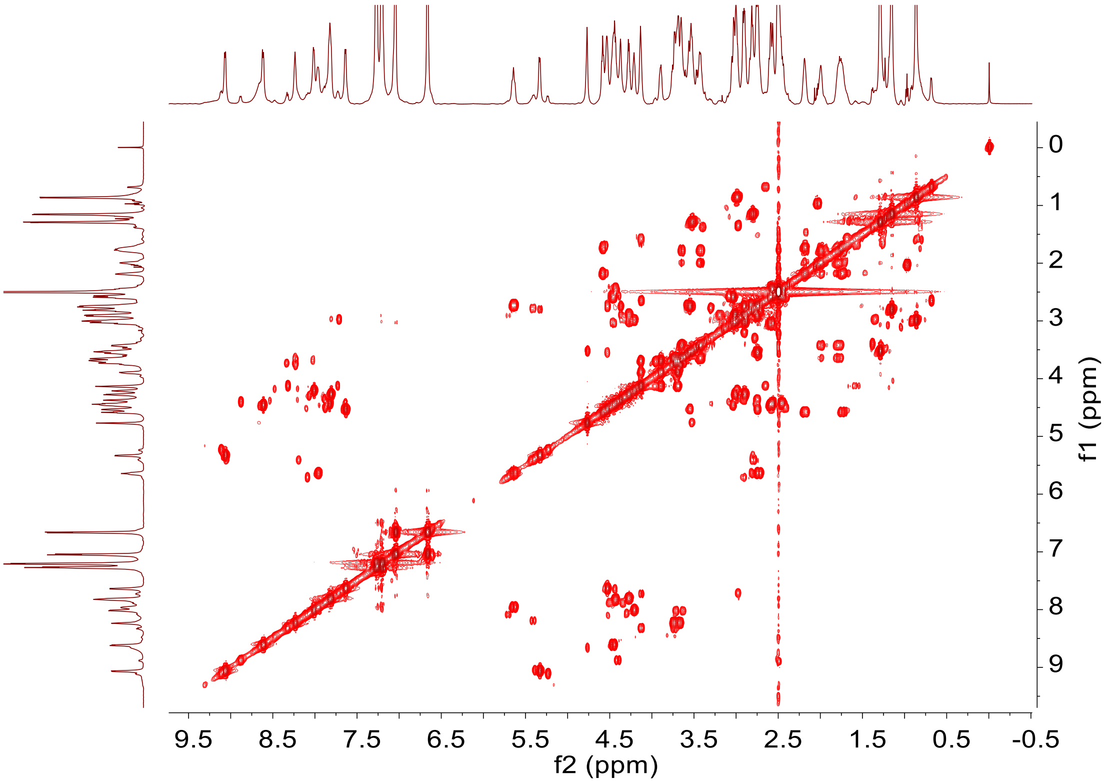

**d) HSQC spectrum (600 MHz, DMSO-$d_6$) of larlan A2 (14)**

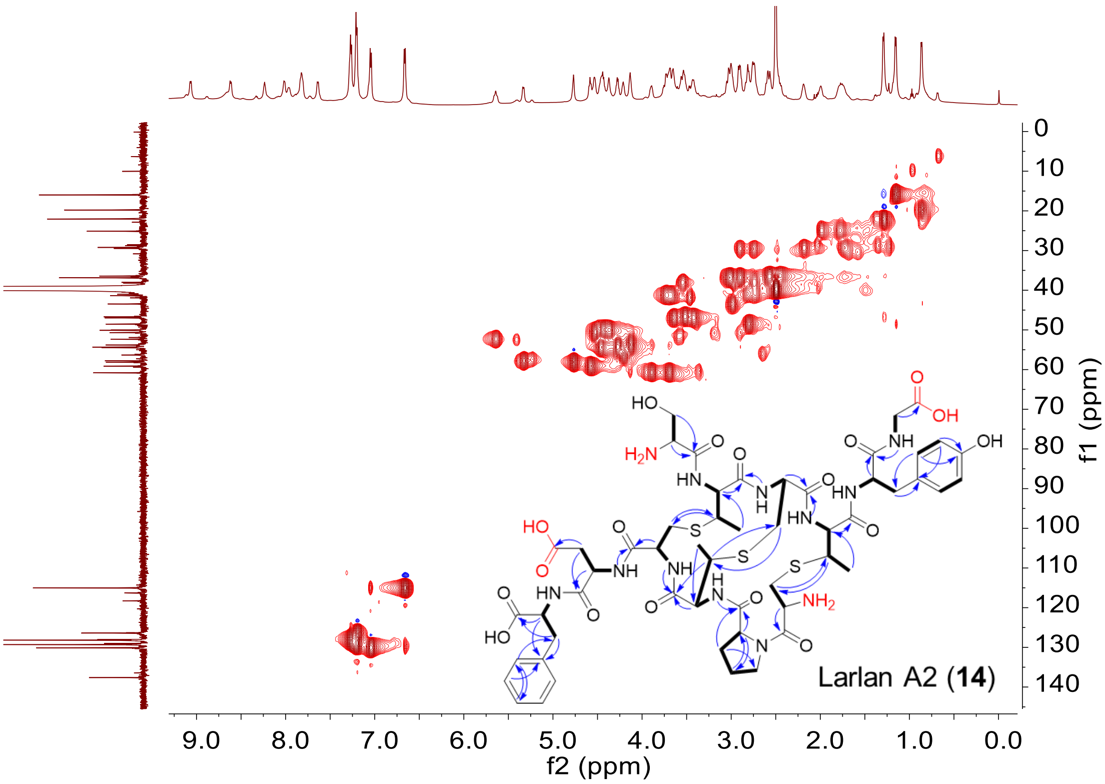

**e) HMBC spectrum (600 MHz, DMSO-$d_6$) of larlan A2 (14)**

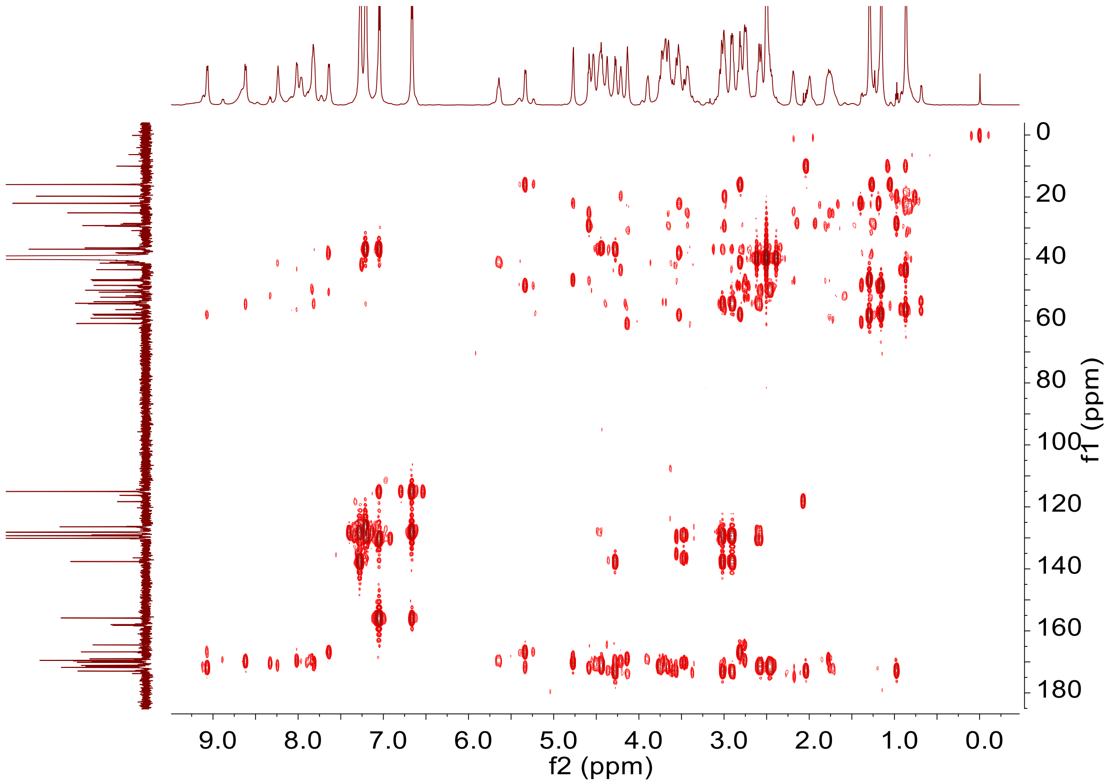

f) NOESY spectrum (600 MHz, DMSO-$d_6$) of larlan A2 (14)

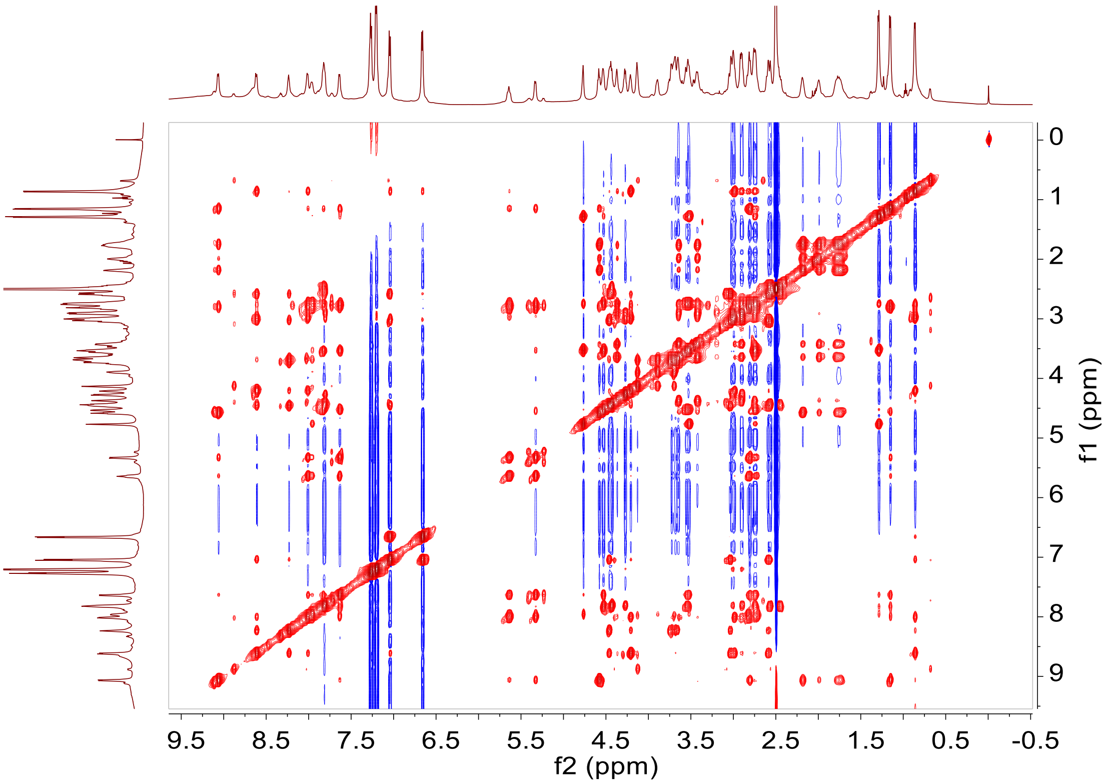

**Supplementary Figure 43. NMR spectra of larlan A2 (14) in DMSO-$d_6$: 600 MHz $^1$H NMR, 150 MHz $^{13}$C NMR, 600 MHz $^1$H-$^1$H COSY, HSQC, HMBC, NOESY spectra**

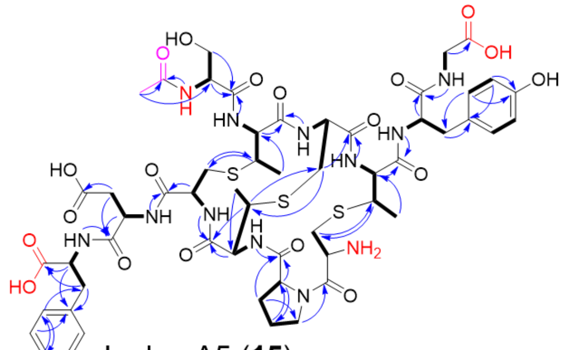

Larlan A5 (**15**)

a) [1]H NMR spectrum (600 MHz, DMSO-$d_6$) of larlan A5 (15)

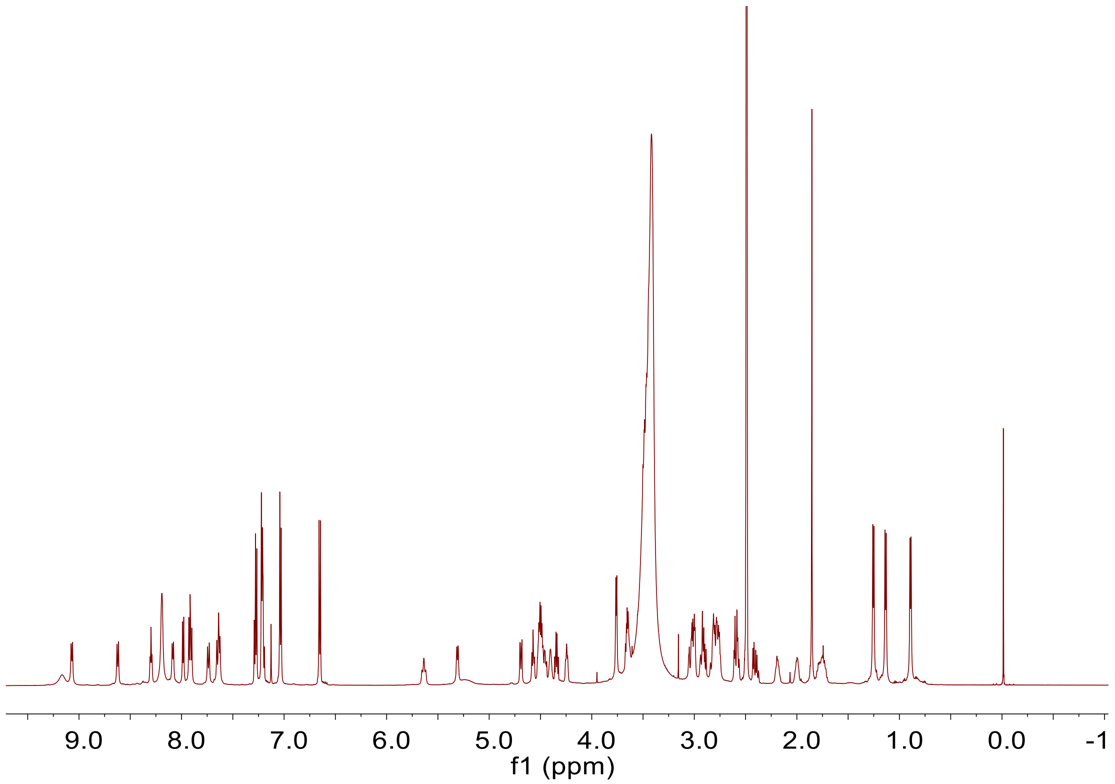

**b) $^{13}C$ NMR spectrum (150 MHz, DMSO-$d_6$) of larlan A5 (15)**

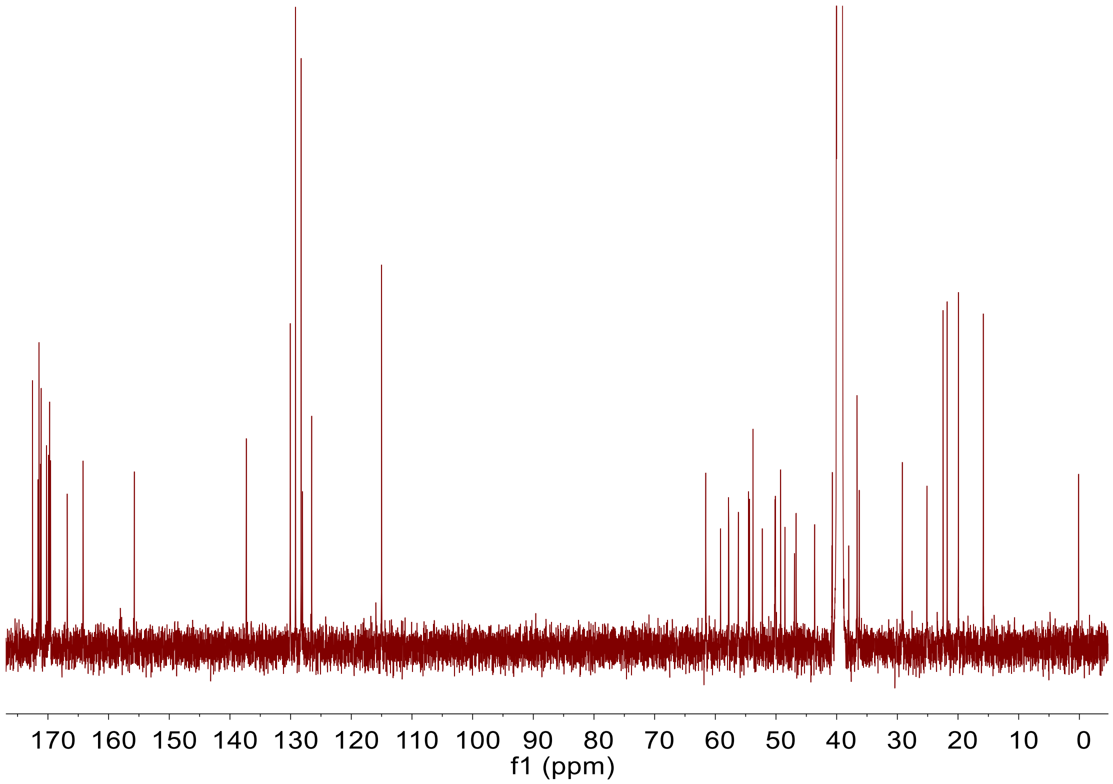

c) [1]H-[1]H COSY spectrum (600 MHz, DMSO-$d_6$) of larlan A5 (15)

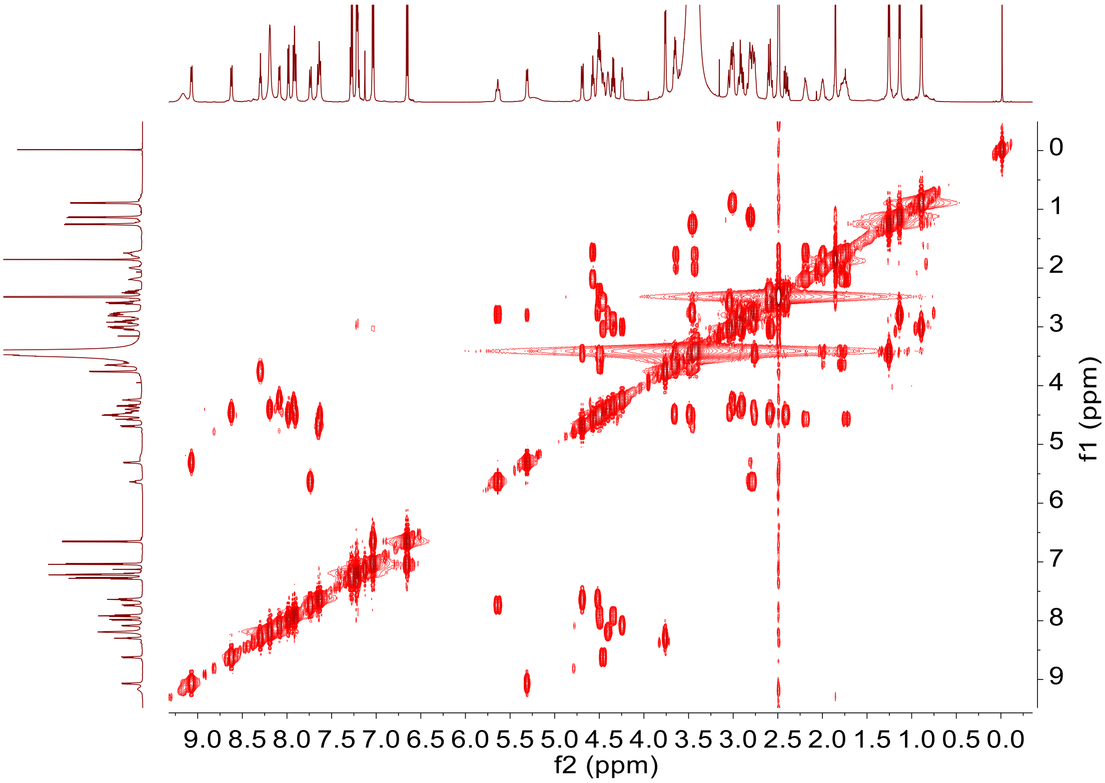

**d) HSQC spectrum (600 MHz, DMSO-$d_6$) of larlan A5 (15)**

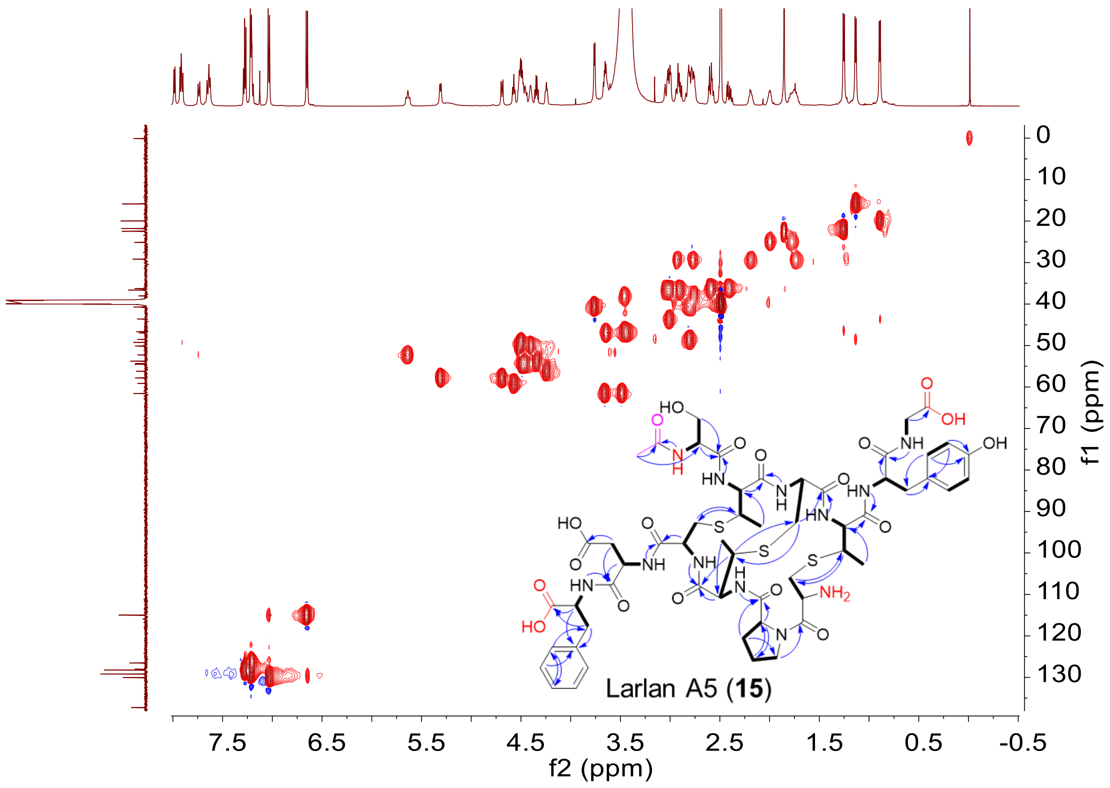

**e) HMBC spectrum (600 MHz, DMSO-$d_6$) of larlan A5 (15)**

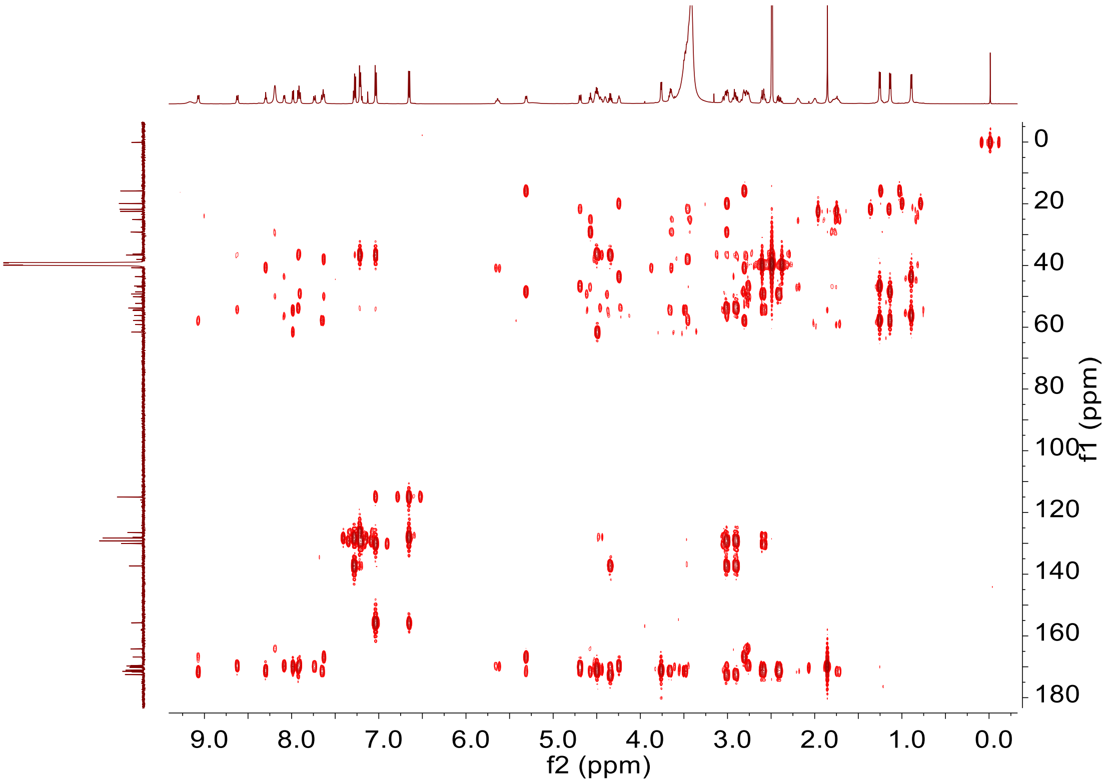

**f) TOCSY spectrum (600 MHz, DMSO-$d_6$) of larlan A5 (15)**

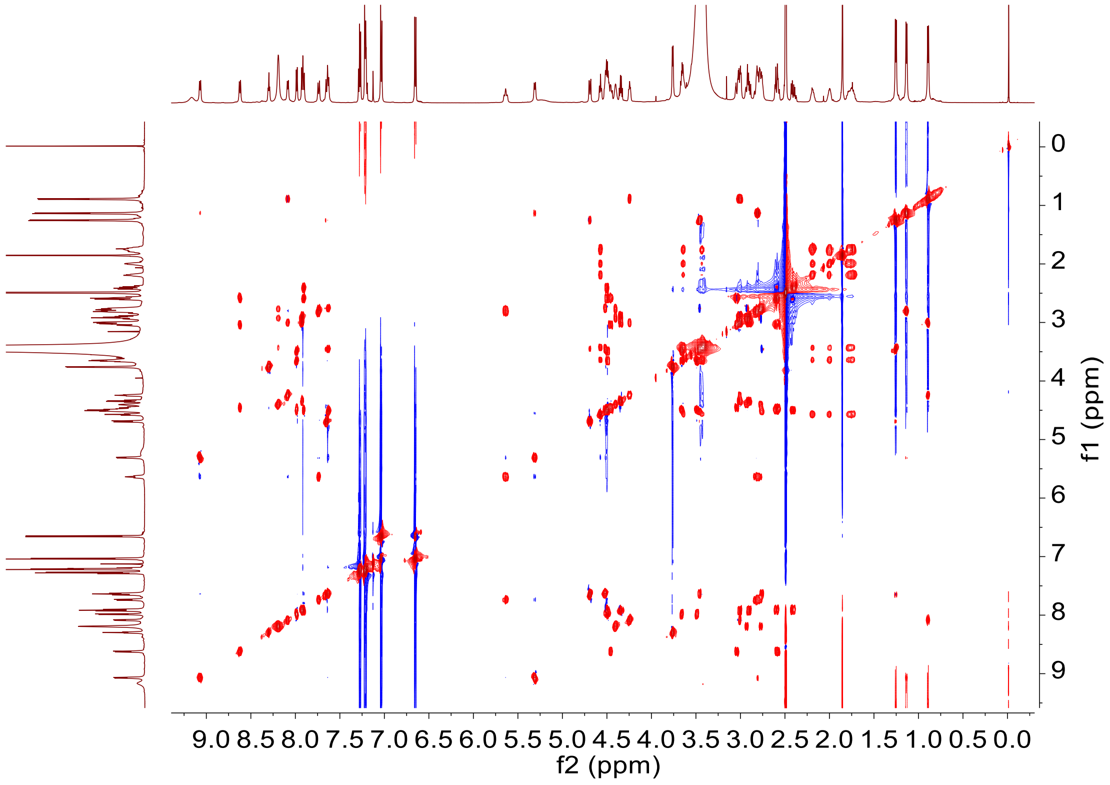

g) NOESY spectrum (600 MHz, DMSO-$d_6$) of larlan A5 (15)

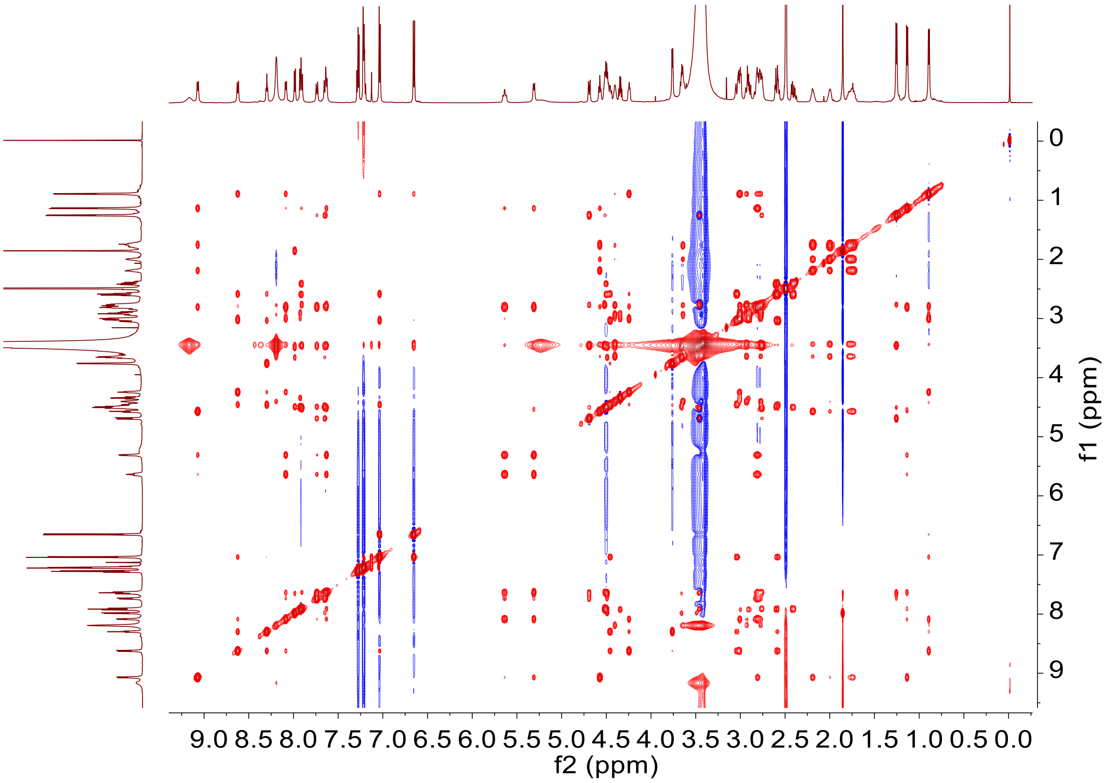

**Supplementary Figure 44. NMR spectra of larlan A5 (15) in DMSO-$d_6$: 600 MHz $^1$H NMR, 150 MHz $^{13}$C NMR, 600 MHz $^1$H-$^1$H COSY, HSQC, HMBC, TOCSY, NOESY spectra**

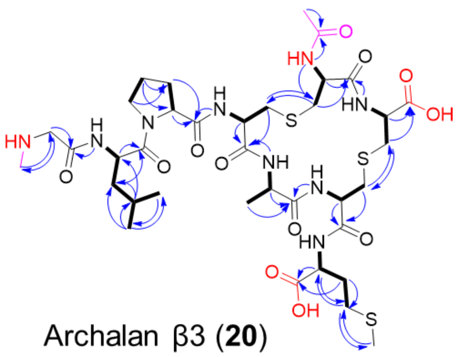

Archalan β3 (**20**)

**a) [1]H NMR spectrum (600 MHz, DMSO-$d_6$) of archalan β3 (20)**

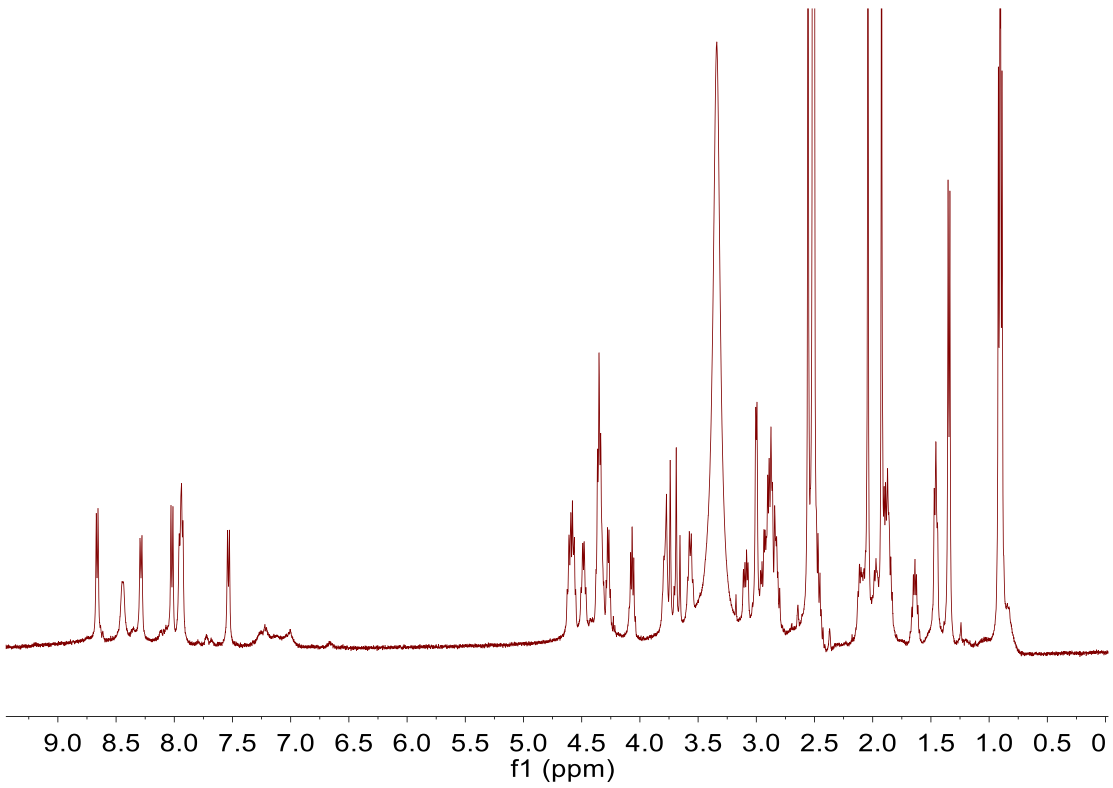

b) $^{13}$C NMR spectrum (150 MHz, DMSO-$d_6$) of archalan β3 (20)

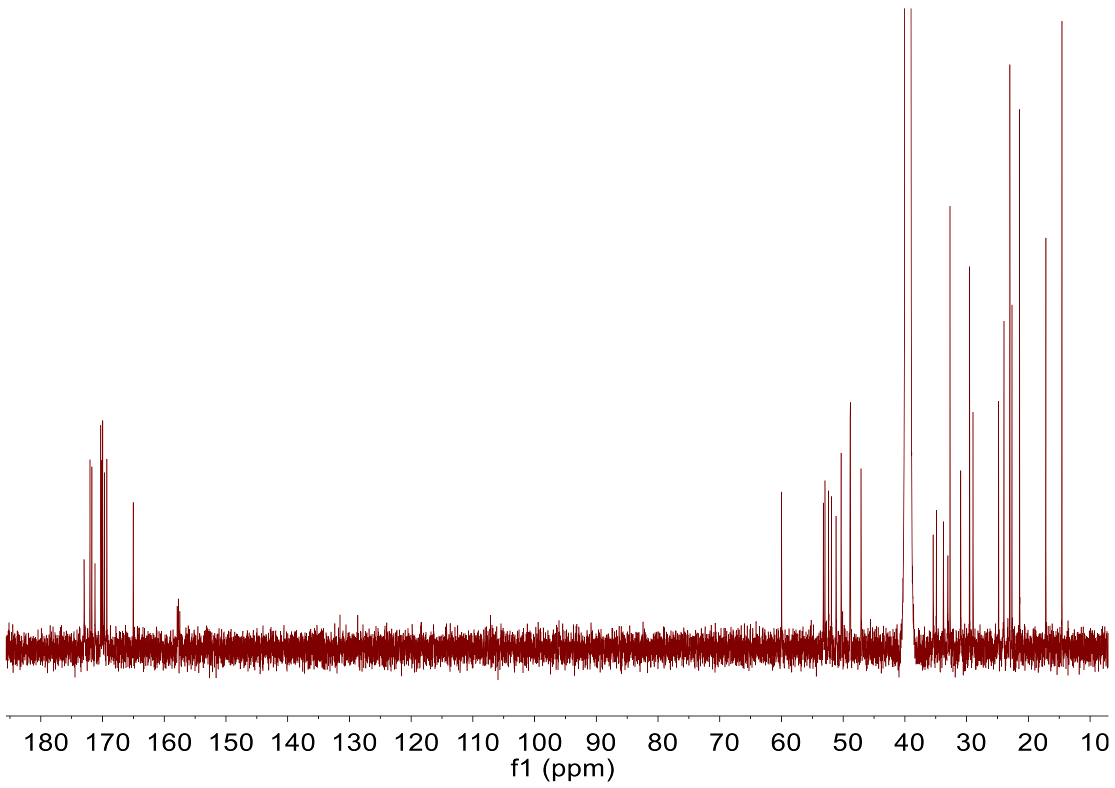

c) [1]H-[1]H COSY spectrum (600 MHz, DMSO-$d_6$) of archalan β3 (20)

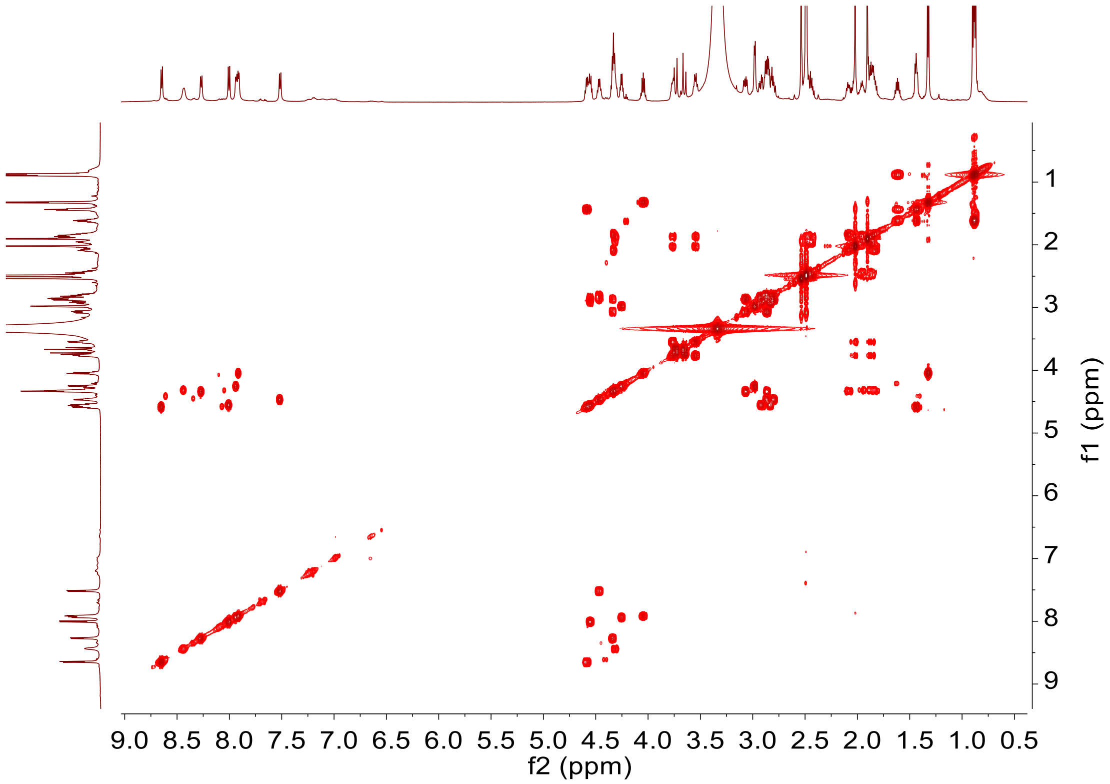

**d) HSQC spectrum (600 MHz, DMSO-$d_6$) of archalan β3 (20)**

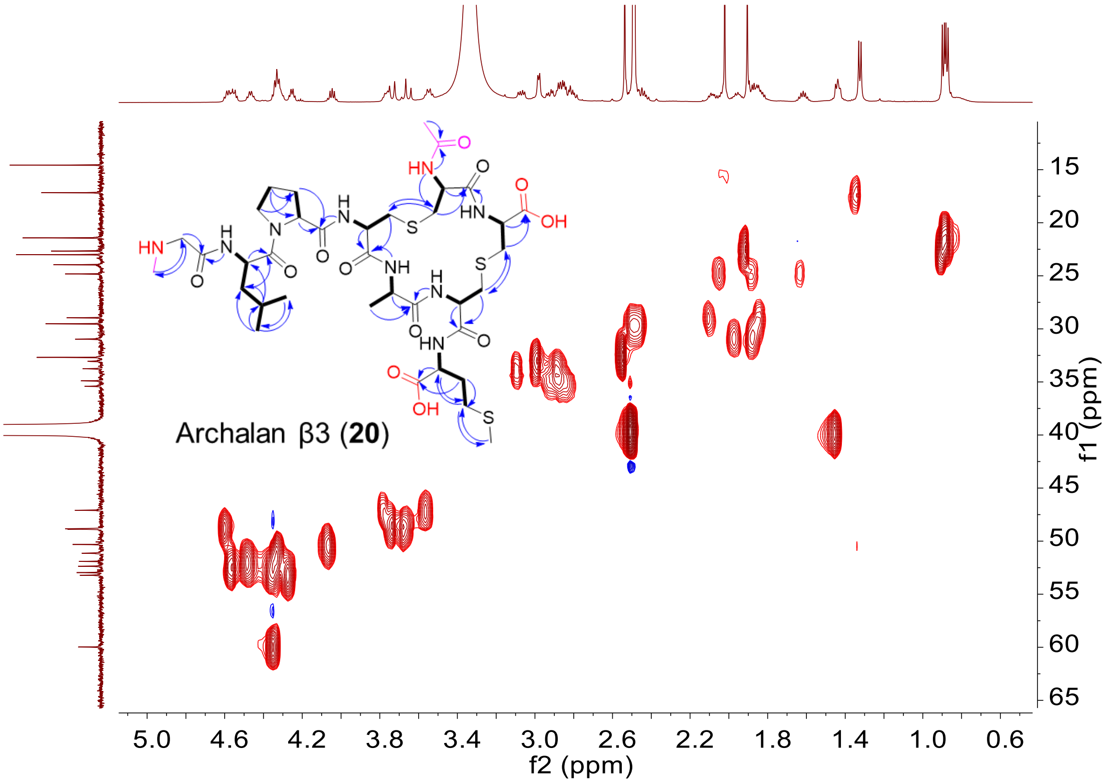

**e) HMBC spectrum (600 MHz, DMSO-$d_6$) of archalan β3 (20)**

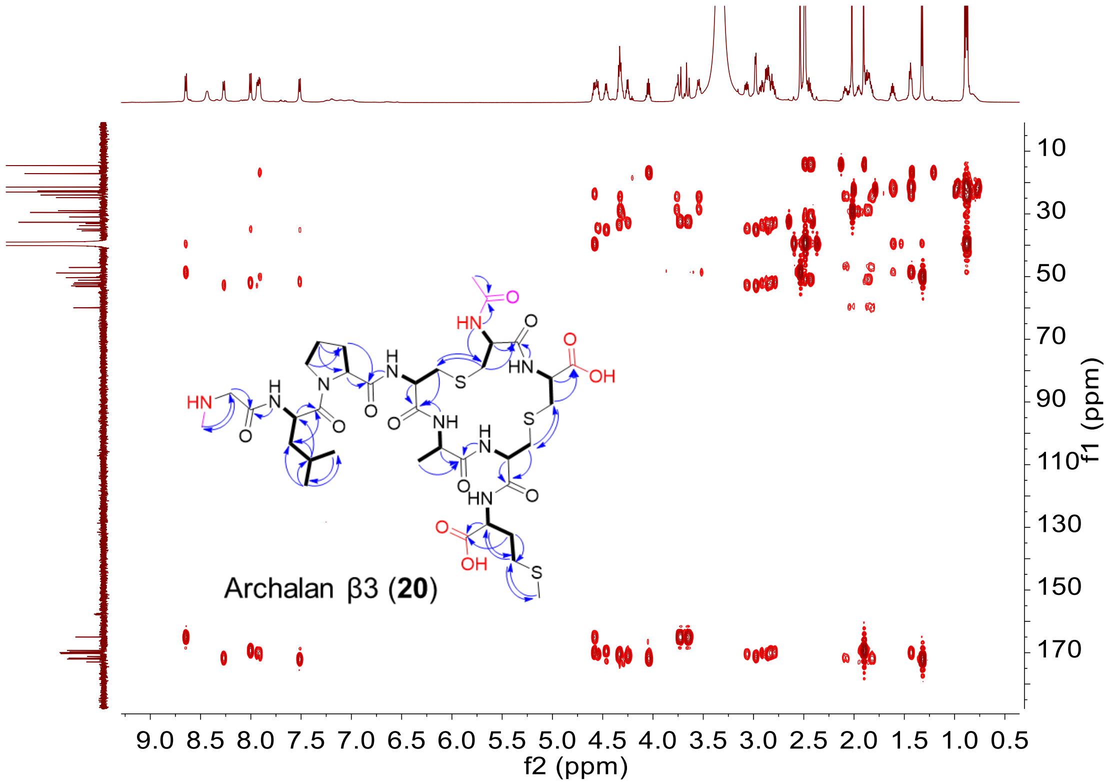

**f) TOCSY spectrum (600 MHz, DMSO-$d_6$) of archalan β3 (20)**

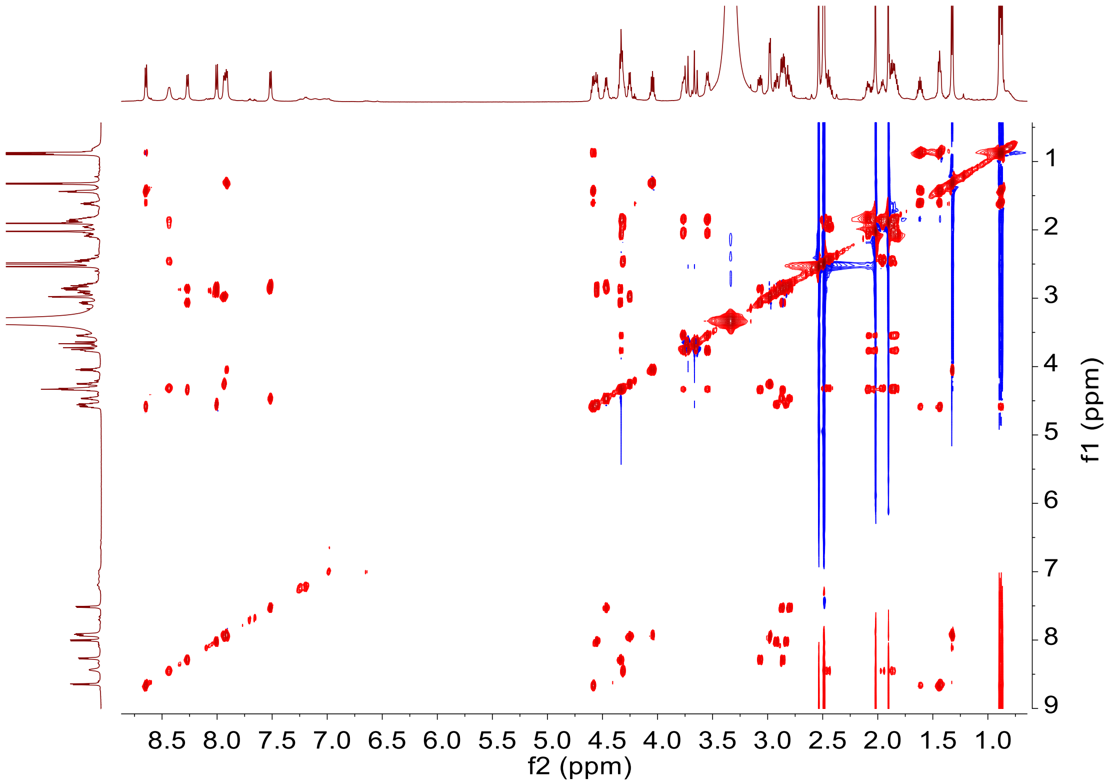

g) NOESY spectrum (600 MHz, DMSO-$d_6$) of archalan β3 (20)

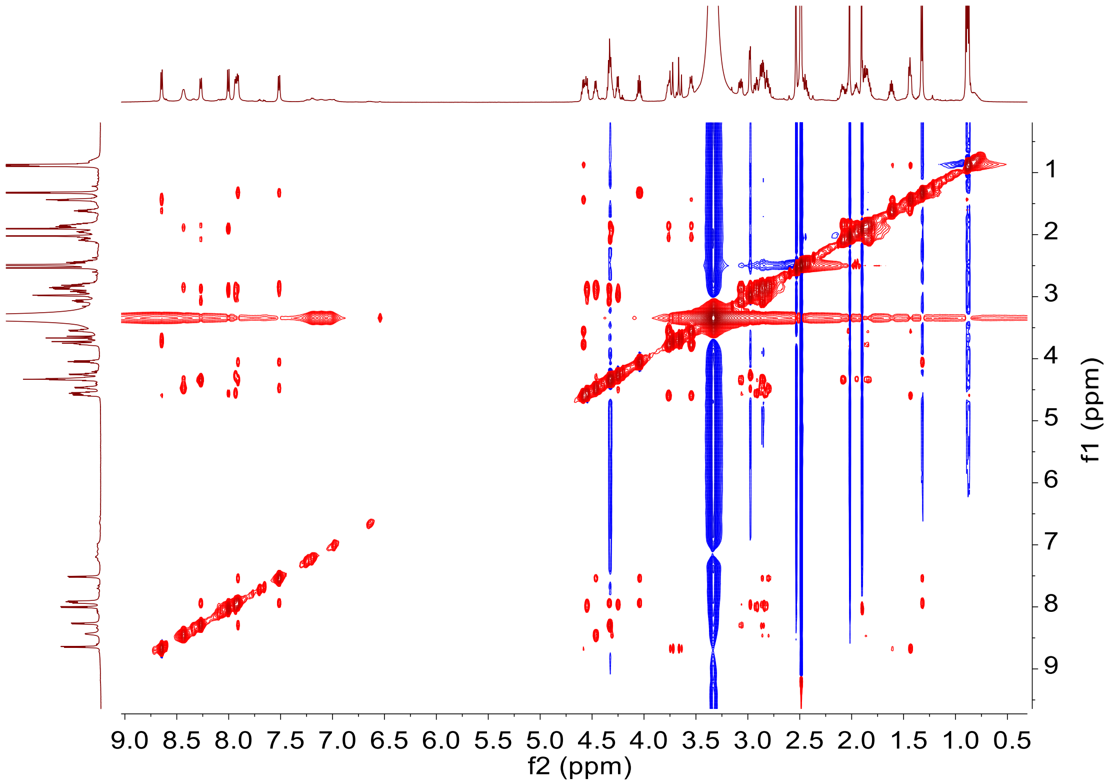

**Supplementary Figure 45. NMR spectra of archalan β3 (20) in DMSO-$d_6$: 600 MHz $^1$H NMR, 150 MHz $^{13}$C NMR, 600 MHz $^1$H-$^1$H COSY, HSQC, HMBC, TOCSY, NOESY spectra**

#### Supplementary Tables

##### Supplementary Table 1. Strains used in this study

| Strain | Genome accession number | Source |
|---|---|---|
| *Haloarcula argentinensis* CGMCC1.6166 | GCF_000336895.1 | CGMCC |
| *Haloferax larsenii* CGMCC1.4296 | GCF_000336955.1 | CGMCC |
| *Haloferax mediterranei* DSM1411 | GCF_000306765.2 | DSMZ |
| *Halomicrobium mukohataei* CGMCC1.6192 | GCF_000023965.1 | CGMCC |
| *Halorussus litoreus* HD8-51 CGMCC1.15333 | GCF_003382685.1 | CGMCC |
| *Haladaptatus cibarius* DSM19505 | GCF_000455345.1 | DSMZ |
| *Haloferax prahovense* JCM13924 | GCF_000336815.1 | JCM |
| *Haloferax denitrificans* DSM4425 | GCF_000337795.1 | DSMZ |
| *Halorussus salinus* YJ-37-H | GCF_004765815.2 | CGMCC |
| *Natrinema gari* JCM14663 | GCF_000337175.1 | JCM |
| *Halomicroarcula salina* JCM18369 | GCF_019061225.1 | JCM |
| *Halorussus amylolyticu* JCM18367 | GCF_004143485.2 | JCM |
| *Halorussus ruber* JCM18363 | GCF_004765785.1 | JCM |
| *Halorussus pelagicus* JCM32953 | GCF_004087835.1 | JCM |
| *Halorussus halobius* JCM31110 | GCF_004765805.1 | JCM |
| *Halobacillus kuroshimensis* JCM14155 | GCF_000425705.1 | JCM |
| *Halobacillus profundi* JCM14154 | / | JCM |
| *Haloferax volcanii* H1424 | / | Thorsten Allers's lab[7] |
| *Methicillin-resistance Staphylococcus aureus* ATCC43300 | / | ATCC |
| *Acinetobacter baumannii* ATCC19606 | | ATCC |
| *Bacillus subtilis* 168 | / | Our lab |
| *Escherichia coli* ATCC25922 | / | Our lab |

#### Supplementary Table 2. Primers used in this study

| Name | Sequence (5'-3') |
|---|---|
| pTA1228_F1 | ATGTCGATAAGCTTGATATCGAATTCCTGCAG |
| pTA1228_R1 | AACGTGAGTTTTCGTTCCACTGAGC |
| pTA1228_F2 | GCTCAGTGGAACGAAAACTCACGTT |
| pTA1228_R2 | CATATGCGCAATAGGTCCGCGAA |
| lar_geneB_bb_F1 | GCGGACCTATTGCGCATATGATGGCTCTGTCGGATTACACCG |
| lar_geneM_R1 | TCAGGACGGCGTCACTCCAT |
| lar_geneH_F2 | ATGGAGTGACGCCGTCCTGAA |
| lar_geneH_bb_R | TTCGATATCAAGCTTATCGACATTCGGCTATCGCGTCTCTGC |
| lar_geneE_F2 | ATGGAGTGACGCCGTCCTGACGATAGCCGACGTTACATCACG |
| lar_geneE_R2 | GGACCATTCCATCTCGGTCAGAC |
| lar_geneT1_F3 | GTCTGACCGAGATGGAATGGTCC |
| lar_geneJ_bb_R3 | TTCGATATCAAGCTTATCGACATGGAGTCGAACGCGGTTCAGT |
| lar_geneT2_bb_R | TTCGATATCAAGCTTATCGACATAGGTCGAGAACTTCGGCTGAA |
| lar_geneA_bb_F | GCGGACCTATTGCGCATATGATTCTCCATGATAAATCTATGCGAGCATC |
| lar_geneM_bb_R | TTCGATATCAAGCTTATCGACATAGGACGGCGTCACTCCATGA |
| alnβ_bb_F1 | GCGGACCTATTGCGCATATGATTGTGTATCGAACCTACCTCGAAGC |
| alnβ_R1 | TTCGATATCAAGCTTATCGACATGGTTTTCAACATTCTCAAGGCCACT |
| alnβ_F2 | TCGGTACTCCTGTTCGAGTGAGTC |
| alnβ_bb_R2 | CGGAACGCCGAATCGTATCGA |
| alnγ_bb_F1 | GCGGACCTATTGCGCATATGTCAACACTCTTTCCCACGTCCG |
| alnγ_R1 | GTGACGAACCGGATGTCAACGA |
| alnγ_F2 | GTCGTTGACATCCGGTTCGTCAC |
| alnγ_bb_R2 | TTCGATATCAAGCTTATCGACATGTTCTCCAACCACTCGACGTTG |
| medb_bb_F1 | GCGGACCTATTGCGCATATGATATCTGAGACAAGACGCTATTCGTTTAATCAG |
| medb_R1 | GCAAAAGTAATTCTCGACAACATACATGTTAAATC |
| medb_F2 | GATTTAACATGTATGTTGTCGAGAATTACTTTTGC |
| medb_bb_R2 | TTCGATATCAAGCTTATCGACATTGATATATAGTTTGATGTCAGCATATCAGACATCTG |
| cib2_bb_F1 | GCGGACCTATTGCGCATATGAAGTCGAAGTCGGCGTCGTCA |
| cib2_R1 | TTTTCTGTCGTCTCGATTACGTTCGGG |
| cib2_F2 | CCCGAACGTAATCGAGACGACAGAAAA |
| cib2_R2 | TCACGAACTGACGTGATGGGAGACG |
| cib2_F3 | CGTCTCCCATCACGTCAGTTCGTGA |
| cib2_bb_R3 | TTCGATATCAAGCTTATCGACATCGTTCAAACAGACACGCCCAACTC |
| sal_bb_R | TTCGATATCAAGCTTATCGACATGTTCTAAGCTAAATGCCATTGGTTGGATATC |
| sal_bb_F | GCGGACCTATTGCGCATATGACCTCGTACGATTTGCATTGTCAATG |
| alnα_bb_F1 | GCGGACCTATTGCGCATATGTGAGAGAGGGGTATGCCGGAC |

| | |
|---|---|
| alnα_R1 | AACTCCATCGCGTCGGTGTTT |
| alnα_F2 | AAACACCGACGCGATGGAGTT |
| alnα_bb_R2 | TTCGATATCAAGCTTATCGACATTTTTTCGCCCTCCCGTTCCAG |
| meda_geneK_bb_F1 | GCGGACCTATTGCGCATATGAACAGTTCCGCCGCGT |
| meda_geneB_R1 | CGAGTAGCCACCTGCAGC |
| meda_geneD_F2 | GGCTGCAGGTGGCTACTCG |
| meda_geneM_R2 | CACCAAGGGTAACCTCACTCCAG |
| meda_geneH_F3 | CTGGAGTGAGGTTACCCTTGGTG |
| meda_geneT1_R3 | GAGTGTGATTCGTCCGAATCAACTG |
| meda_geneT1_F4 | AGTTGATTCGGACGAATCACACTCACCAG |
| meda_geneJ_bb_R4 | TTCGATATCAAGCTTATCGACATTTAGTAGAACTCGCGGACGAGGTCCATC |
| meda_geneB_bb_F1 | GCGGACCTATTGCGCATATGCGCTTCATCTGTCAATAAGTGCCAT |
| cib1_bb_F | GCGGACCTATTGCGCATATGTCACCACGGTCGAAGACGTAGTG |
| cib1_bb_R | TTCGATATCAAGCTTATCGACATGGAAGTAAAGGACTACAGTTCGAAATCCTTC |
| lit_bb_R2 | TTCGATATCAAGCTTATCGACATGAGAGAACGTGCAATGCATCGTC |
| lit_bb_F1 | GCGGACCTATTGCGCATATGACGGTATAAGAGTGGAGCGTTCGAA |
| lit_R1 | GGTCCTACTACAAGAACCTTGTGCAAAATATG |
| lit_F2 | CATATTTTGCACAAGGTTCTTGTAGTAGGACC |
| pel_bb_R | TTCGATATCAAGCTTATCGACATGTCGATTTTCGGCGTCACGTAGG |
| pel_bb_F | GCGGACCTATTGCGCATATGGCATCGTCTACAGGTACGACGGG |

**Supplementary Table 3. NMR data for archaeal lanthipeptides: sallan A2 (2) and ciblan A1 (3) in MeOH-$d_4$, medlan A1 (13), larlan A2 (14), larlan A5 (15), and archalan β3 (20) in DMSO-$d_6$ (600 MHz $^1$H NMR, 150 MHz $^{13}$C NMR)**

| Sallan A2 (2) | | | | Ciblan A1 (3) | | | | Archalan β3 (20) | | | |
|---|---|---|---|---|---|---|---|---|---|---|---|
| AA | $\delta_H$, mult(*J* in Hz) | $\delta_C$ | Key HMBC | AA | $\delta_H$, mult(*J* in Hz) | $\delta_C$ | Key HMBC | AA | $\delta_H$, mult(*J* in Hz) | $\delta_C$ | Key HMBC |
| Ala-1 | 4.27, t (4.5) | 53.4 | 168.8 | Ala-1 | 4.10 | 50.5 | 171.1 | *N*-Me-Gly-1 | 3.73, 3.65 | 48.8 | 32.7, 165.0 |
| | 2.96, dd (5.8, 13.8); 3.31 | 34.3 | 35.9, 168.8 | | 1.54, d (7.0) | 17.9 | 171.1 | | 2.54 | 32.7 | 48.8 |
| | | 168.8 | | | | 171.1 | | | | 165.0 | |
| Thr-2 | 4.38, d (4.2) | 61.4 | 173.1 | Ala-2 | 4.45, t (6.8) | 53.5 | 171.6 | Leu-2 | 8.65, d (7.9) | | 165.0 |
| | 4.2, q (5.6) | 68.6 | 173.1 | | 2.97, 2.92 | 33.8 | 35.0, 171.6 | | 4.59 | 48.8 | 170.0 |
| | 1.25, d (6.4) | 20.5 | 61.4 | | | 171.6 | | | 1.44 | 39.7 | 170.0 |
| | | 173.1 | | Lys-3 | 4.11 | 55.5 | 171.6, 173.9 | | 1.63 | 24.8 | 48.8 |
| Arg-3 | 3.94, t (7.2) | 55.8 | 26.6, 173.9 | | 1.83 | 30.3 | 27.9, 173.9 | | 0.88, d (6.7) | 22.7 | 21.4, 39.7 |
| | 1.86 | 28.9 | 42.3, 173.9 | | 1.50 | 23.9 | 40.6, 55.5 | | 0.89, d (6.7) | 21.4 | 22.7 |
| | 1.55 | 26.6 | 55.8 | | 1.69 | 27.9 | 30.3, 40.6 | | | 170.0 | |
| | 3.16 | 42.3 | 28.9, 158.8 | | 2.95 | 40.6 | 23.9, 27.9 | Pro-3 | 4.35 | 60.0 | 171.7 |
| | | 158.8 | | | | 173.9 | | | 1.84,2.09 | 29.0 | 171.7 |
| | | 173.9 | | Leu-4 | 4.33 | 52.7 | 173.6 | | 1.89, 2.05 | 24.8 | 29.0, 60.0 |
| Tyr-4 | 4.08, dd (4.1, 11.5) | 58.6 | 127.8, 172.9 | | 1.70, 1.46 | 40.5 | 21.6, 23.4, 173.6 | | 3.76,3.55 | 47.2 | 29.0 |
| | 3.16, 3.33 | 35.2 | 127.8, 131.4, 172.9 | | 1.78 | 25.9 | 52.7 | | | 171.7 | |
| | | 127.8 | | | 0.96 | 21.6 | 40.5, 23.4 | Ala-4 | 8.27, d (7.5) | | 171.7 |
| | 7.03, d (8.2) | 131.4 | 127.8, 157.4, 131.4 | | 1.00, d (6.5) | 23.4 | 40.5, 21.6 | | 4.34 | 53.0 | 170.3 |
| | 6.74, d (8.2) | 116.6 | 127.8, 157.4 | | | 173.6 | | | 2.87,3.07 | 33.8 | 34.9, 170.3 |
| | | 157.4 | | Pro-5 | 4.48 | 61.5 | 23.3, 173.5 | | | 170.3 | |

| | | | | | | | | | | | |
|---|---|---|---|---|---|---|---|---|---|---|---|
| | | 172.9 | | | 2.28 | 32.8 | 48.1, 173.5 | Ala-5 | 7.92 | | 170.3 |
| Ala-5 | 4.42, dd (4.3, 6.9) | 53.9 | 172.3 | | 1.90, 2.13 | 23.3 | 61.5 | | 4.05, m | 50.3 | 172.0 |
| | 3.07, 3.24 | 35.9 | 34.3, 172.3 | | 3.71, 3.50 | 48.1 | 32.8 | | 1.32, d (7.1) | 17.2 | 172.0 |
| | | 172.3 | | | | 173.5 | | | | 172.0 | |
| Phe-6 | 4.63, dd (4.2, 8.3) | 55.9 | 38.6 | Ala-6 | 4.56 | 54.5 | 173.0 | Ala-6 | 7.52, d (8.1) | | 172.0 |
| | 3.15, 3.25 | 38.6 | 138.9, 130.5 | | 2.93, 3.16 | 35.0 | 173.0 | | 4.46, m | 51.9 | 169.7 |
| | | 138.9 | | | | 173.0 | | | 2.80,2.87 | 35.4 | 33.1, 169.7 |
| | 7.34, d (7.6) | 130.5 | 130.5, 127.9, 138.9 | Ala-7 | 4.80 | 53.5 | 169.7 | | | 169.7 | |
| | 7.29, t (7.6) | 129.6 | 127.9, 138.9 | | 3.19, 2.98 | 32.9 | 169.7 | Met-7 | 8.43 | | |
| | 7.21, t (7.4) | 127.9 | 130.5 | | | 169.7 | | | 4.32 | 51.3 | 173.0, 29.5 |
| | | | | Asp-8 | 4.86 | 51.2 | 171.1 | | 1.96, 1.86 | 30.9 | 29.5, 173.0 |
| | | | | | 2.78, 2.87 | 38.8 | 171.1, 174.9 | | 1.84, 2.46 | 29.5 | 51.3, 14.6 |
| | | | | | | 174.9 | | | 2.02, s | 14.6 | 29.5 |
| | | | | | | 171.1 | | | | 173.0 | |
| | | | | Abu-9 | 4.32 | 56.3 | 173.7 | *N*-Ac-Ala-8 | 1.90, s | 22.7 | 169.3 |
| | | | | | 3.86 | 41.2 | 32.9, 173.7 | | | 169.3 | |
| | | | | | 1.20, d (7.1) | 20.7 | 56.3 | | 8.01, d (8.7) | | 169.3, 34.9 |
| | | | | | | 173.7 | | | 4.56 | 52.4 | 170.1 |
| | | | | Dhb-10 | | 131.0 | | | 2.83,2.92 | 34.9 | 33.8, 170.1 |
| | | | | | 6.66, q (7.1) | 130.9 | 131, 165.9 | | | 170.1 | |
| | | | | | 1.86, d (7.1) | 13.4 | 131.0 | Ala-9 | 7.94 | | 170.1 |
| | | | | | | 165.9 | | | 4.25 | 53.2 | 171.2 |
| | | | | Asn-11 | 5.22 | 50.5 | | | 2.98 | 33.1 | 35.4, 171.2 |
| | | | | | 3.02, 2.68 | 36.7 | 174.8 | | | 171.2 | |
| | | | | | | 174.8 | | | | | |
| **Medlan A1 (13)** | | | | **Larlan A2 (14)** | | | | **Larlan A5 (15)** | | | |
| **AA** | **$\delta_H$, mult(*J* in Hz)** | **$\delta_C$** | **Key HMBC** | **AA** | **$\delta_H$, mult(*J* in Hz)** | **$\delta_C$** | **Key HMBC** | **AA** | **$\delta_H$, mult(*J* in Hz)** | **$\delta_C$** | **Key HMBC** |

| | | | | | | | | | | | |
|---|---|---|---|---|---|---|---|---|---|---|---|
| Ser-1 | 8.23 | | | Ser-1 | 4.14, t (5.3) | 53.8 | 169.0 | N-Ac-Ser-1 | 1.86 | 22.5 | 54.4/54.5, 169.9 |
| | 4.15 | 53.2 | | | 3.70, 3.89 | 60.8 | 169.0 | | | 169.9 | |
| | 3.72, q (6.0); 3.88, q (5.5) | 60.5 | 168.8 | | | 169.0 | | | 7.99, d (6.9) | | 169.9 |
| | | 168.8 | | Abu-2 | 8.01, d (5.9) | | | | 4.49 | 54.4/54.5 | 171.6 |
| Abu-2 | 8.51 | | 168.8 | | 4.21, t (6.1) | 56.4 | 169.7 | | 3.66, 3.49 | 61.6 | 171.6 |
| | 4.82 | 58.2 | 169.8 | | 2.98 | 43.5 | 29.4 | | | 171.6 | |
| | 3.50 | 46.5 | 21.8, 38.3 | | 0.85, d (5.9) | 19.8 | 56.4 | Abu-2 | 8.08, d (7.3) | | 171.6 |
| | 1.30, d (7.2) | 21.8 | 58.2 | | | 169.7 | | | 4.23 | 56.2 | 169.7 |
| | | 169.8 | | Ala-3 | 8.61, d (7.8) | | 169.7 | | 3.01 | 43.6 | 29.2 |
| Ala-3 | 7.72, d (9.8) | | 169.8 | | 4.46 | 54.3 | 169.9 | | 0.89, d (6.5) | 19.9 | 56.2 |
| | 5.56 | 52.6 | 168.8 | | 2.59, 3.04 | 36.9 | 48.5 | | | 169.7 | |
| | 2.86, 2.81 | 40.9 | 49.1 | | | 169.9 | | Ala-3 | 7.63 | | 169.7 |
| | | 168.8 | | Abu-4 | 8.66 | | 169.9 | | 4.51 | 50.1 | 46.7 |
| Abu-4 | 7.91, d (8.4) | | 41.9, 168.8 | | 4.77 | 58.2 | 170.1 | | 2.75, 3.46 | 38.0 | 171.1 |
| | 4.26, t (7.6) | 56.2 | 18.5, 169.8 | | 3.52 | 46.7 | 38.1 | | | 171.1 | |
| | 3.32, m | 41.9 | 30.1 | | 1.29, d (6.4) | 22.0 | 58.2 | Abu-4 | 9.07, d (7.5) | | 171.1 |
| | 0.99, d (6.6) | 18.5 | 56.2 | | | 170.1 | | | 5.31, d (7.8) | 57.8 | 166.8 |
| | | 169.8 | | Tyr-5 | 7.83 | | | | 2.80 | 48.5 | 40.7/40.8 |
| Tyr-5 | 8.53, d (8.3) | | 169.8 | | 4.44, m | 54.5 | 171.1 | | 1.14, d (7.2) | 15.9 | 57.8 |
| | 4.45 | 53.9 | 127.1, 172.8 | | 2.60, 2.56 | 36.9/36.4 | 128.1 | | | 166.8 | |
| | 2.79, 2.90 | 36.6/36.3 | 127.1 | | | 128.1 | | Tyr-5 | 8.62, d (8.4) | | 166.8 |
| | | 127.1 | | | 7.04, d (8.1) | 130.1 | 36.9/36.4, 155.9 | | 4.45 | 54.3 | 171.1 |
| | 7.00, d (8.4) | 130.1 | 115.0, 156.0 | | 6.67, d (7.7) | 115.0 | 128.1, 155.9 | | 2.60, 3.04 | 36.3 | 128.0 |
| | 6.65, d (8.4) | 115.0 | 127.1, 130.1, 156.0 | | | 155.9 | | | | 128.0 | |
| | | 156.0 | | | | 171.1 | | | 7.04, d (8.4) | 130.1 | 36.3, 156.8 |
| | | 172.8 | | Gly-6 | 8.24 | | 171.1 | | 6.68, d (8.1) | 115.0 | 128.0, 156.8 |

| | | | | | | | | | | | |
|---|---|---|---|---|---|---|---|---|---|---|---|
| Ala-6 | 8.28 | | | | 3.65/3.73 | 41.4 | 171.5 | | | 156.8 | |
| | 3.98 | 50.6 | 30.1 | | | 171.5 | | | | 171.1 | |
| | 3.16, 2.82 | 30.1 | 41.9, 167.0 | Ala-7 | 7.63 | | | Gly-6 | 8.29, d (5.9) | | 171.1 |
| | | 167.0 | | | 4.53, m | 50.7 | 169.7 | | 3.76, d (5.8) | 40.7 | 171.1 |
| His-7 | 8.75, d (6.6) | | 26.7, 167.0 | | 2.74, 3.55 | 38.1 | 46.7 | | | 171.1 | |
| | 4.68 | 52.0 | 170.4 | | | 169.7 | | Ala-7 | 7.74, d (9.9) | | |
| | 3.0, 3.08 | 26.7 | 116.8, 128.9, 170.4 | Pro-8 | 4.58, t (6.8) | 59.1 | 25.1, 171.7 | | 5.63, t (10.8) | 52.3 | |
| | | 128.9 | | | 2.19, 1.73 | 29.2 | 47.0, 171.7 | | 2.77, 2.93 | 40.7/40.8 | 48.5 |
| | 7.43, s | 116.8 | 133.9 | | 1.98, 1.78 | 25.1 | 29.2, 47.0, 59.1 | | | 169.9 | |
| | 9.00, s | 133.9 | 116.8, 128.9 | | 3.42, 3,65, | 47.0 | 29.2, 25.2 | Pro-8 | 4.57 | 59.1 | 25.1, 171.6 |
| | | 170.4 | | | | 171.7 | | | 1.73, 2.19 | 29.2 | 47.0, 171.6 |
| Abu-8 | 9.12, d (8.4) | | 170.4 | Abu-9 | 9.07, d (7.1) | | 171.7 | | 1.78, 2.00 | 25.1 | 59.1 |
| | 5.37, d (8.1) | 57.5 | 166.6 | | 5.33, d (7.2) | 57.8 | 166.8 | | 3.45, 3.63 | 47.0 | 169.9 |
| | 2.75 | 49.1 | 40.9 | | 2.80 | 48.5 | 36.9, 166.8 | | | 171.6 | |
| | 1.16, d (7.2) | 15.8 | 57.5 | | 1.15, d (7.0) | 16.0 | 57.8 | Abu-9 | 7.65 | | 171.6 |
| | | 166.6 | | | | 166.8 | | | 4.69 | 57.8 | 170.2 |
| Ala-9 | 7.65, d (7.2) | | 38.3, 166.6 | Ala-10 | 7.63, d (6.4) | | 166.8 | | 3.45 | 46.7 | 38.0, 170.2 |
| | 4.51 | 50.2 | 169.4 | | 4.37, m | 54.3 | 164.5 | | 1.25, d (6.9) | 21.8 | 57.8 |
| | 2.78, 3.52 | 38.3 | 46.5 | | 2.75, 2.90 | 29.4 | 43.5 | | | 170.2 | |
| | | 169.4 | | | | 164.5 | | Ala-10 | 8.19 | | 170.2 |
| Asp-10 | 7.95, d (7.9) | | 169.4 | Asp-11 | 7.96 | | 164.5 | | 4.41 | 50.1 | 171.1 |
| | 4.49 | 49.2 | 170.4 | | 5.64, t (10.7) | 52.4 | 169.5 | | 2.77, 2.93 | 29.2 | 43.6 |
| | 2.40, 2.59 | 36.3 | 171.4 | | 2.75, 2.73 | 41.0 | 164.6, 169.5 | | | 171.1 | |
| | | 171.4 | | | | 164.6 | | Asp-11 | 7.91 | | 171.1 |
| | | 170.4 | | | | 169.5 | | | 4.49 | 49.2 | 170.2 |
| Phe- | 7.98, d (7.7) | | 170.4 | Phe- | 7.81 | | | | 2.41, 2.59 | 36.2 | 170.2, 171.2 |

| | | | | | | | | | | | |
|---|---|---|---|---|---|---|---|---|---|---|---|
| 11 | | | | 12 | | | | | | | |
| | 4.31 | 53.9 | 137.5, 170.4, 172.6 | | 4.27, q (6.1) | 54.3 | 137.7, 173.0 | | | 171.2 | |
| | 2.90, 3.01 | 36.6/36.3 | 137.5 | | 2.90, 3.00 | 36.9/36.4 | 137.7, 173.0 | | | 170.2 | |
| | | 137.5 | | | | 137.7 | | Phe-12 | 7.92 | | 170.2 |
| | 7.23 | 129.2 | 126.4 | | 7.19, m | 129.3 | 36.9/36.4, 126.4 | | 4.35 | 53.8 | 137.3, 172.5 |
| | 7.28 | 128.2 | 137.5 | | 7.26, t (7.2) | 128.2 | 137.7 | | 2.90, 3.00 | 36.6 | 137.3, 172.5 |
| | 7.21 | 126.4 | 137.5 | | 7.21, m | 126.4 | 129.3 | | | 137.3 | |
| | | 172.6 | | | | 173.0 | | | 7.20/7.22 | 129.2 | 36.6, 126.5 |
| | | | | | | | | | 7.28 | 128.3 | 137.3 |
| | | | | | | | | | 7.20/7.22 | 126.5 | 129.2 |
| | | | | | | | | | | 172.5 | |

**Supplementary Table 4. Advanced Marfey's analysis of archaeal lanthipeptides**

| Sallan A2 (**2**) | | | | Ciblan A1 (**3**) | | | |
|---|---|---|---|---|---|---|---|
| Residues | L-FDLA | D-FDLA | Configuration | Residues | L-FDLA | D-FDLA | Configuration |
| Thr | 13.8 | 15.1 | L | Ala | 14.9 | 15.9 | L |
| Arg | 12.9 | 12.5 | L | Lys | 10.3 | 17.1 | L |
| Tyr-Bis | 19.3 | 20.2 | L | Leu | 16.5 | 18.7 | L |
| Phe | 16.7 | 18.2 | L | Pro | 14.9 | 15.6 | L |
| di-FDLA-lanthionine* | 17.8 | 17.8 | DL | Asp | 14.0 | 14.1 | L |
| | | | | Thr | 13.7 | 15.0 | L |
| | | | | di-FDLA-methyllanthionine* | 19.1 | 15.5 | LL |
| | | | | di-FDLA-lanthionine* | 17.7 | 17.8 | DL |
| Archalan β3 (**20**) | | | | Medlan A1 (**13**) | | | |
| Residues | L-FDLA | D-FDLA | Configuration | Residues | L-FDLA | D-FDLA | Configuration |
| Ala | 14.9 | 16.6 | L | Ser | 13.9 | 14.3 | L |
| Ser | 14.3 | 14.5 | L | Phe | 16.8 | 18.5 | L |
| Pro | 15.2 | 16.3 | L | Tyr | 19.0 | 19.6 | L |
| Leu | 17.0 | 19.5 | L | His-Bis | 15.7 | 15.8 | L |
| Met-O | 17.1 | 18.9 | L | His | 12.4 | 12.0 | L |
| mono-FDLA-lanthionine* | 12.0, 12.5 | 12.2, 12.7 | LL | Asp | 14.1 | 14.5 | L |
| | | | | di-FDLA-methyllanthionine* | 17.5, 19.4 | 15.9, 20.7 | LL & DL |
| Larlan A2 (**14**) | | | | Larlan A5 (**15**) | | | |

| Residues | L-FDLA | D-FDLA | Configuration | Residues | L-FDLA | D-FDLA | Configuration |
|---|---|---|---|---|---|---|---|
| Ser | 14.0 | 14.3 | L | Ser | 14.1 | 14.3 | L |
| Pro | 15.0 | 15.9 | L | Pro | 15.1 | 15.9 | L |
| Phe | 16.9 | 18.5 | L | Phe | 16.9 | 18.5 | L |
| Tyr-Bis | 19.5 | 20.5 | L | Tyr-Bis | 19.6 | 20.1 | L |
| Asp | 14.2 | 14.4 | L | Asp | 14.2 | 14.5 | L |
| di-FDLA-methyllanthionine* | 17.5, 19.4 | 15.9, 20.7 | LL & DL | di-FDLA-methyllanthionine* | 17.5, 19.4 | 15.9, 20.7 | LL & DL |
| Residues | L-FDLA | D-FDLA | Configuration | Residues | L-FDLA | D-FDLA | Configuration |
| Larlan A2-T2S-di-FDLA-methyllanthionine | 17.5, 19.9 | 15.9, 20.7 | LL & DL | Larlan A2-T4S-di-FDLA-methyllanthionine | 17.5 | 20.7 | LL |
| Standard-di-FDLA-DL-lanthionine | 17.8 | 17.8 | DL | Standard-di-FDLA-DL-methyllanthionine | 19.4 | 15.9 | DL |
| Standard-di-FDLA-LL-lanthionine | 17.8 | 17.0 | LL | Standard-di-FDLA-LL-methyllanthionine | 17.5 | 20.7 | LL |
| Standard-mono-FDLA-DL-lanthionine | 12.3, 12.9 | 12.3, 12.9 | DL | Standard-di-FDLA-D-*allo*-L-methyllanthionine | 16.8 | 15.9 | D-*allo*-L |
| Standard-mono-FDLA-LL-lanthionine | 12.0, 12.5 | 12.2, 12.7 | LL | Standard-di-FDLA-L-*allo*-L-methyllanthionine | NA | 13.5 | L-*allo*-L |

* Compared with standard samples. NA represents no data.

The values in the table indicate the retention time (in minutes) of the respective L/D-FDLA derivatives in LR/HR LC-MS analysis.
